# Mapping the genetic landscape of the DNA damage response with Cas12a-based combinatorial knockout screens

**DOI:** 10.64898/2026.06.07.728858

**Authors:** Samuel B. Hayward, Alina Vaitsiankova, Tomas Lama-Diaz, Juihsuan Chou, Angelo Taglialatela, Jen-Wei Huang, Yodharaudshani Wijesekarahanthi, Joshua R. Heyza, Giuseppe Leuzzi, Chuanyuan Chen, Nancy Wong, Tenzin Lhakhang, Xi Fu, Alejandro L. Buendia, Veronica Gheorghe, Nazanin Esmaeili Anvar, Jens C. Schmidt, Andre Nussenzweig, Raul Rabadan, Vincenzo Costanzo, Raphaël Guérois, Traver Hart, Alberto Ciccia

## Abstract

The DNA damage response (DDR) is a complex network of cellular pathways that ensures the faithful maintenance of our genomes upon a wide array of genomic insults. To elucidate the functional architecture of this network, we conducted unbiased genetic interaction screens using the Cas12a genome editor to disrupt 233 DDR genes frequently mutated in cancer and other genetic diseases, either individually or in pairwise combinations. This approach enabled us to assess the phenotypic effects induced by the disruption of >27,000 DDR gene pair combinations under unperturbed cell growth conditions. From this analysis, we identified over 750 high-confidence positive (buffering) or negative (synthetic lethal/sick) gene-gene interactions, along with multiple connections between previously unlinked DDR pathways and modules, allowing us to define novel aspects of the cellular response to spontaneous, DNA replication-associated DNA damage. Among the identified genetic interactions, we uncovered profound synthetic lethal interactions between genes encoding 1) the translesion polymerase REV1-Pol ζ complex and the MCM8-MCM9-HROB DNA helicase complex; 2) Fanconi Anemia (FA) proteins and the mitotic DNA repair factors GEN1, CIP2A, and RHINO; and 3) the DNA translocase SMARCAL1 and components of the FANCM complex, suggesting novel opportunities for targeted therapies in tumors carrying mutations in these genes. Additionally, we identified robust suppressor interactions between the *DCLRE1B* gene encoding the nuclease APOLLO and the core non-homologous end joining (NHEJ) genes *XRCC4*, *LIG4*, and *NHEJ1*, suggesting that NHEJ impairs the fitness of APOLLO-deficient cells. This work provides a functional map of the DDR network and demonstrates the power of Cas12a-based screens for identifying synthetic lethal and buffering interactions with therapeutic potential.

## Main

The DNA damage response (DDR) is a cellular program that has evolved to counteract a diverse range of genomic alterations occurring throughout an individual’s lifespan. It is estimated that each day a single cell can encounter ∼10^5^ spontaneous DNA lesions, ranging from minor base alterations to severe DNA double-strand breaks (DSBs)^1^. In response to DNA damage, the DDR arrests cell cycle progression and coordinates the repair of DNA lesions, or initiates cell death or cellular senescence if the damage is irreparable^2^. Failure to repair genomic lesions can result in the persistence of mutations, culminating in genomic instability and the onset of pathological phenotypes. Indeed, deficiencies in DDR genes are known to cause a variety of genetic syndromes characterized by cancer predisposition, growth retardation, immunodeficiency, neurodegeneration, neurodevelopmental defects, and/or premature aging^2^. Given the frequency and diversity of DNA lesions, the DDR has evolved into a highly sophisticated network comprising numerous specialized pathways. These DDR pathways exhibit functional overlap, share protein components, and can compensate for one another despite having unique functions.

Pairwise genetic interactions reveal functional relationships between genes through the combined loss of their functions^3–7^. A pairwise genetic interaction is observed when the phenotypic outcome from the loss of one gene is modulated by the loss of a second gene^8^. Genetic interactions can be classified as negative or positive. Pairwise negative interactions occur when a double mutant phenotype is more severe than expected based on the observed single mutant phenotypes, an example being synthetic lethality, which arises when mutations in two non-essential genes result in a non-viable phenotype. Negative genetic interactions may reflect functional redundancies, where two genes can independently perform similar tasks, or situations in which one gene inhibits cellular processes that would be toxic in the absence of the second gene^9,10^. Conversely, positive interactions occur when the loss of two genes results in a less severe outcome than expected based on the observed phenotypes of the single mutants. Positive interactions may suggest that the two genes operate in the same biological pathway (within-pathway interaction), or that the function of one gene becomes toxic in the absence of the other, such that loss of the toxic gene suppresses the observed deleterious phenotype (suppressor interaction)^9,10^.

Exploring the genetic interaction landscape of the DDR has been a longstanding pursuit, driven by the interconnected nature of the DDR network and the recognition that genomic instability is a hallmark of cancer cells, making cancer-associated alterations in DDR genes promising targets for precision oncology^11^. Consequently, genetic screens have prioritized the identification of DDR genes whose disruption induces lethality or impairs drug treatments in cancer-relevant contexts. In the present study, we employed Cas12a-based combinatorial knockout (KO) screens to generate a genetic interaction map documenting all pairwise interactions for 233 genes associated with the DDR in human cells. Screening results identified over 750 high-confidence pairwise genetic interactions, and network-level analysis highlighted synthetic lethal interactions among gene modules, including those between the MCM8-MCM9-HROB DNA helicase module and the translesion synthesis (TLS) polymerase REV1-MAD2L2-REV3L module, between the Fanconi anemia (FA) gene network and the mitotic DNA repair genes *GEN1*, *CIP2A,* and *RHNO1*, and between FANCM-associated modules (*i.e.*, FANCM-FAAP24 and CENPS-CENPX) and the *SMARCAL1* DNA translocase gene. Our analyses also uncovered robust suppressor interactions between the *DCLRE1B* gene (encoding the nuclease APOLLO) and a core non-homologous end joining (NHEJ) module containing the *XRCC4*, *LIG4*, and *NHEJ1* genes. These studies reveal the functional architecture of the DDR network and establish Cas12a-based knockout screens as a powerful approach for mapping genetic interaction networks at scale.

### Combinatorial Cas12a screening maps genetic interactions in the human DDR

To investigate genetic interactions within the DDR network, we utilized Cas12a, a class 2, type V CRISPR-Cas system that can process a compact CRISPR RNA (crRNA) array into multiple guide RNAs (gRNAs), making it optimal for combinatorial screening^12^. Unlike previous Cas9-based approaches, which require individual expression of multiple gRNAs, Cas12a reduces the need for multiple promoters and features short, sequence-flexible direct repeats between gRNAs, reducing both the difficulty of library cloning and the potential for confounding recombination during lentivirus production and sequencing^7,13–15^. In our studies, we employed the Cas12a in4mer platform, which uses constructs featuring 4-gRNA arrays, with two gRNAs targeting each gene of interest within a pair^16^ (Fig. 1a). In4mer-mediated redundant gene targeting in a single construct increases the likelihood of homozygous double knockout (dKO) of two genes, which has been a significant challenge in combinatorial KO-based screening^16^. The challenge of double gene KO using one gRNA per gene arises from the fact that, even with highly efficient gRNAs achieving editing rates of 80%, using one gRNA per gene would result in a theoretical ∼41% homozygous dKO rate in diploid cells (Extended Data Fig. 1a). However, incorporating a second 80% efficient gRNA for each gene doubles the theoretical dKO rate (Extended Data Fig. 1b). For this reason, redundant gene targeting with the in4mer system allows for more robust genetic interaction calling than single gRNA screening.

**Fig. 1.**
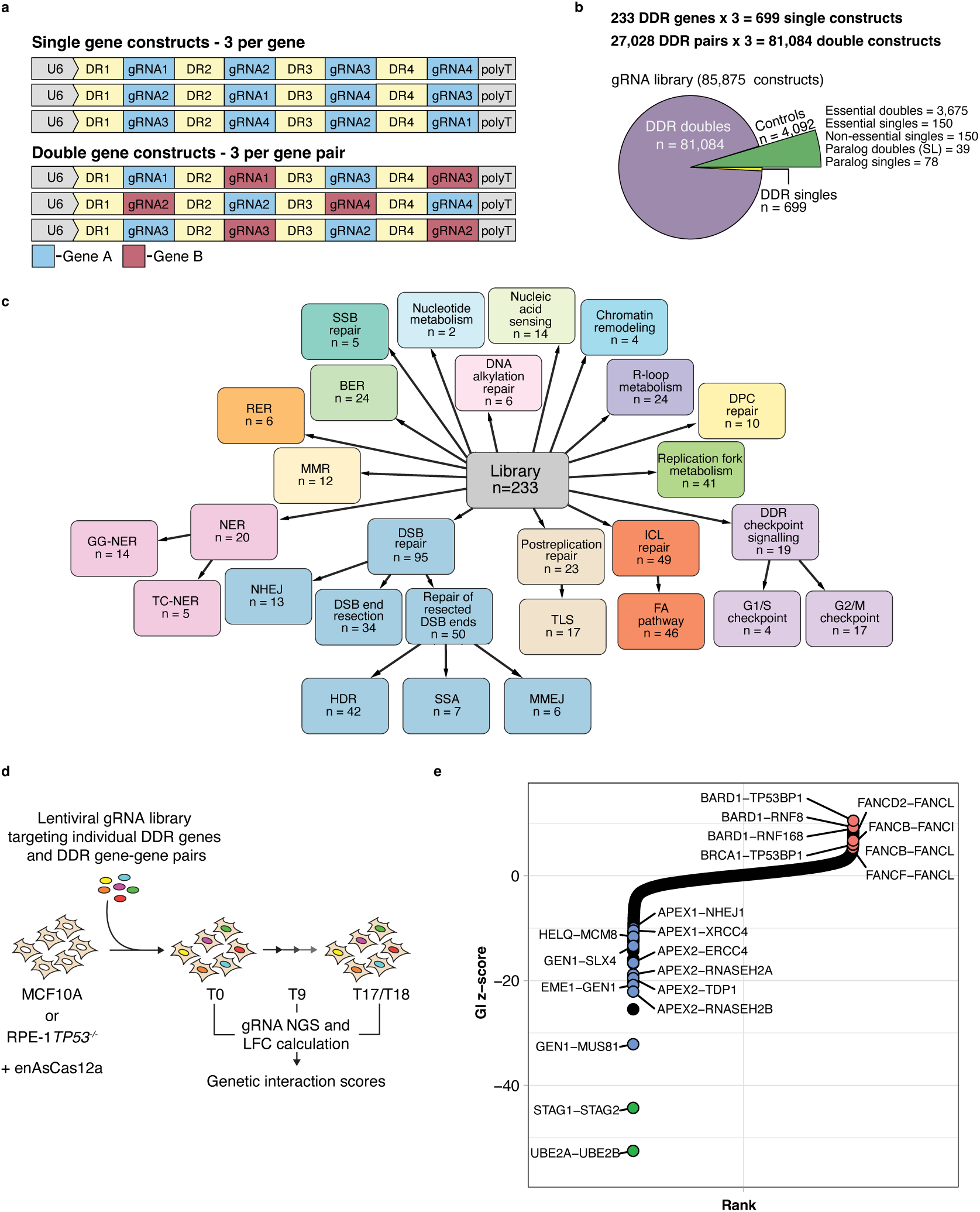
Cas12a-based combinatorial screening of DDR genetic interactions. **a**, Illustration of Cas12a in4mer gRNA constructs containing 4 gRNAs targeting single genes and gene pairs used in the screen. **b**, Summary of DDR GI library gRNA composition. Total number (n) of gRNAs targeting DDR gene pairs, DDR single genes, and controls is indicated. **c**, DDR pathway annotations for genes in the GI library. Terminal pathways and their gene members are represented in their parent node. Individual genes can appear in multiple pathways. **d**, Schematic of GI combinatorial screen carried out in MCF10A and RPE-1 *TP53 ^-/-^* clonal cell lines constitutively expressing enAsCas12a. Following selection of MCF10A or RPE-1 *TP53^-/-^* cells transduced with the lentiviral gRNA GI library (T0), cells were collected at day 9 (T9, early) and 17/18 (T17/T18, final) for sequencing and downstream analysis. **e**, GI screen results for the RPE-1/MCF10A_integrated dataset combining both time points (early and final) across the two cell lines. Known positive interactions, negative interactions, and synthetic lethal paralog pairs are highlighted in red, blue, and green, respectively.

In our in4mer DDR library, we selected four gRNAs for each gene of interest. Constructs for single gene targeting incorporate all four gRNAs, whereas those designed for gene pairs utilize two gRNAs per gene. Single and pair targets encompass three distinct constructs, each comprising permutations of the available gRNAs to control for positional effects and increase robustness (Fig. 1a). For our genes of interest, we selected a total of 233 DDR genes, ensuring representation of the diverse pathways of the DDR network (Fig. 1c and Supplementary Table 1). Additionally, we included 13 synthetic lethal paralog pairs and their corresponding single gene counterparts as positive controls for genetic interactions^16–21^ (Fig. 1b). Reference essential and nonessential gene controls were also included. Altogether, our gRNA library consists of 81,084 constructs targeting 27,028 DDR gene pairs, 699 constructs targeting 233 individual DDR genes, and 4,092 controls (Fig. 1b and Supplementary Table 2).

For the screen, we generated an MCF10A clonal cell line stably expressing an enhanced version of Cas12a from *Acidaminococcus* sp. (enAsCas12a, referred to as enCas12a) to ensure high editing efficiency^22,23^. We selected MCF10A cells, a spontaneously immortalized human mammary epithelial cell line derived from normal breast tissue, due to their stable near-diploid karyotype^24–26^, which minimizes confounding effects of copy number variation between genomic loci that could otherwise affect KO efficiency and influence phenotypic outcomes of combinatorial gRNA targeting. To complement this model, we also generated a stable enCas12a-expressing clone of hTERT-immortalized RPE-1 *TP53^-/-^* retinal pigment epithelial cells (hereafter referred to as RPE-1 in the main text), similarly chosen for their stable karyotype^27^. We employed *TP53^-/-^* cells to reduce cell cycle arrest and/or cell death resulting from p53 activation induced by loss of key DDR genes, such as *BRCA1* and *BRCA2*^28–31^.

To perform our genetic interaction screens, enCas12a-expressing MCF10A and RPE-1 cells were transduced with the DDR in4mer lentiviral library and an initial timepoint (T0) was collected after selection of the transduced cells. MCF10A and RPE-1 cells were then passaged every three days at 1000x or 500x representation for a total of 17 (T17) or 18 (T18) days following T0, respectively (Fig. 1d). Screens were performed in triplicate and in4mer constructs from T9 and T17 (or T18) were sequenced and used as an early and final timepoint in downstream analysis, respectively.

Following sequencing, gRNA log fold changes (LFCs) were calculated and compared to DepMap gene essentiality scores to determine if single gene fitness was accurately represented in our screen^32^. This analysis revealed a strong overall correlation between the LFCs of our single gene constructs in MCF10A and RPE-1 cells at the final time point and DepMap gene effect scores, indicating that our screen is capable of accurately calling single gene phenotypes (Extended Data Fig. 2a). Additionally, essential gene controls showed a significant depletion in our screens, unlike nonessential genes, further validating the use of the in4mer architecture for functional screening of single gene essentiality (Extended Data Fig. 2a). To determine the ability of our screens to call genetic interactions, a log additive model was used to predict expected pairwise phenotypes^33,34^. Utilizing this model, we noted a significant negative deviation between the expected and observed LFCs for all synthetic lethal paralog pair controls in MCF10A and RPE-1 cells, confirming the capability of our screening method to accurately identify genetic interactions (Extended Data Fig. 2b).

Genetic interaction (GI) scores were then determined using the GRAPE (Genetic Interaction Regression Analysis of Pairwise Effects) pipeline^35^. GRAPE leverages all pairwise measurements to estimate single mutant fitness (SMF) for each gene using a linear regression model, under the assumption that most gene pairs do not exhibit interactions (Extended Data Fig. 3a,b). These SMF estimates are subsequently used to model expected dKO effects, and deviations between observed and expected values are defined as candidate genetic interactions. We next applied a dynamic range filter to remove gene pairs whose expected dKO fitness falls outside the measurable range of the screen, which would otherwise generate artifactual positive interactions involving essential genes. Finally, GI z-scores were calculated using a local variance normalization approach (Extended Data Fig. 3c,d and Supplementary Table 3). Notably, this normalization further reduced the number of significant false-positive interactions, particularly buffering interactions between essential genes (Extended Data Fig. 3e,f).

The GRAPE methodology was applied to individual time points (T9 and T17/T18) (Supplementary Table 3). From these analyses, we identified 142 and 355 high-confidence interactions with FDR scores below 0.1 at the early time point, and 163 and 462 high-confidence interactions at the final time point in the MCF10A and RPE-1 datasets, respectively (Extended Data Fig. 2c and Supplementary Table 4). At the early time point, approximately 40% of significant interactions identified in MCF10A cells were also significant in RPE-1 cells, increasing to 50% at the final time point (Extended Data Fig. 2c and Supplementary Table 5). Together, the two screens yielded more than 750 high-confidence interactions (Supplementary Table 4). Stouffer’s method was used to integrate the screening data between the two time points within each cell line (MCF10A_integrated; RPE-1_integrated). Correlation analysis between the MCF10A_integrated and RPE-1_integrated datasets (FDR < 0.1) revealed strong concordance (R = 0.81), demonstrating the reproducibility of our screen across independent cellular contexts (Extended Data Fig. 2d). Given this concordance, we also generated combined cross-cell-line datasets at the early time point (RPE-1/MCF10A_T9_integrated), final time point (RPE-1/MCF10A_T18/T17_integrated), and across all time points (RPE-1/MCF10A_integrated) (Supplementary Table 3). Through this approach, we recognized known negative and positive genetic interactions in all datasets, including suppressor interactions between *BRCA1/BARD1* and *TP53BP1* pathway genes and within-pathway interactions between FA genes in the RPE-1/MCF10A_integrated dataset (Fig. 1e)^36–38^. Similarly, the synthetic lethal interactions of *GEN1* with *MUS81*, *HELQ* with *MCM8*, *APEX1* with NHEJ genes, and *APEX2* with *TDP1*, *RNASEH2A/B* and *ERCC4* were also identified in the RPE-1/MCF10A_integrated dataset (Fig. 1e)^39–42^.

To validate synthetic lethal interaction calling, we compared the GI z-scores from our RPE-1/MCF10A_T18/T17_integrated dataset to differential essentiality scores from three previously published screens conducted in APEX1-, APEX2-, and TDP1-deficient cell lines^42^. Our screens were able to identify the top hits from these isogenic background screens, showing that our pooled genetic interaction screening approach can identify similar hits as one-dimensional screens (Extended Data Fig. 2e). Additionally, our screen confirmed that *APEX2* displays a greater number of genetic interactions than *APEX1* and *TDP1*, as previously reported^42^, suggesting that our screening approach can help nominate subsets of genes that have denser sets of genetic interactions.

### Genetic interaction mapping identifies therapeutically relevant targets

A broad goal of genetic interaction screening is to identify synthetic lethal partners of cancer-associated genes, which can be exploited for precision oncology. Analysis of the frequency of pathogenic mutations within the GENIE database (AACR) highlighted the enrichment of cancer-suppressors in our library gene set (Extended Data Fig. 4a)^43^. Within our library, we assigned 45 genes a cancer-associated status based on their presence in the top 25% of mutated genes in the GENIE database (Extended Data Fig. 4a). Mutations in these genes were found in a range of cancer patient samples, making their synthetic lethal partners potential targets for precision oncology (Extended Data Fig. 4b and Supplementary Table 6). Among the negative genetic interactions with these cancer-associated genes in our RPE-1/MCF10A_integrated dataset, we identified the known synthetic lethal interactions of *PRKDC* and *XRCC1* with *ATM*, and *ERCC4* with *APEX2* (Extended Data Fig. 4c)^42,44–46^. In addition, several of the other negative interactions involved functionally related genes, in support of their validity as *bona fide* synthetic lethal interactions, including the shared interactions of both *RNASEH2A* and *RNASEH2B* with *ATM,* and those of FA genes with *GEN1* (Extended Data Fig. 4c). These interactions provide strong candidates for further validation to assess their potential as therapeutic targets.

Similarly, we examined positive interactions with DDR-associated disorder genes to identify cooperating genes that might prevent disease occurrence and uncover interactors that could be targeted to ameliorate the pathological outcomes induced by the loss of these disease genes. Analysis of the ClinVar database revealed pathogenic germline mutations in 121 of our library genes (Extended Data Fig. 5a)^47^. From this set of ClinVar mutations, 61 of the genes in our library harbored a mutation associated with hereditary cancers, while 107 genes carried mutations assigned to a non-cancer genetic disease (Extended Data Fig. 5a,b and Supplementary Table 6). Among positive interactions involving DDR-associated disease genes, we identified positive interactions between the Shu complex genes *ZSWIM7* and *SPIDR* and the Bloom syndrome helicase *BLM*, in line with observations that loss of the Shu complex member *Swsap1* prolongs *Blm*-mutant embryo survival^48^ (Extended Data Fig. 5c). We also observed a positive interaction between *ERCC6L2*, a gene mutated in patients with bone marrow failure^49^, and the ataxia telangiectasia gene *ATM*, consistent with recent studies showing that ERCC6L2 loss promotes resistance to ATM inhibition^50^. Additionally, we uncovered positive interactions between the NHEJ genes *XRCC4*, *LIG4*, and *NHEJ1* and the Hoyeraal-Hreidarsson syndrome gene *DCLRE1B*^51^, in keeping with the rescue of *Apollo* mutant lethality by *Ku70* deficiency in mice^52^. The ability of our screen to recapitulate functional interactions through differential cell viability in untreated conditions validates our approach and nominates positive interactors for further characterization.

### Genetic interaction map identifies novel DDR pathway interactions

A key advantage of high-throughput genetic interaction screening is the ability to construct genetic interaction profiles for individual genes. These profiles comprise all genetic interaction measurements for one gene against the rest of the library, providing a high-resolution phenotypic description of that gene’s function. Unlike functional profiles derived from less tractable phenotypes (*i.e.*, gene essentiality in response to drug treatments or across different cell lines), genetic interaction profiles from pairwise perturbations are directly traceable to individual gene effects, enabling the identification of interactions between gene modules^53,54^. To determine gene modules, we conducted clustering analysis of our MCF10A_integrated, RPE-1_integrated, and RPE-1/MCF10A_integrated datasets (Extended Data Fig. 6a, 7a, and 8a), as well as analysis of clustering at individual time points in both cell lines (https://ddr-genetic-interactions.ciccialab-database.com/; see also Data Availability section). Due to the sensitivity of clustering analysis to input data, gene module assignment was performed on a co-clustering frequency metric, the GI odds ratio, that considers the propensity of gene pair co-clustering across a large set of filtered input matrices. Clustering of the GI odds ratio analyses identified 53, 52, and 58 distinct modules of two or more genes in the MCF10A_integrated, RPE-1_integrated, and RPE-1/MCF10A_integrated datasets, respectively (Extended Data Fig. 6a, 7a, 8a and Supplementary Table 7).

By correlating genetic interaction profiles, it becomes possible to effectively discern information about protein complexes. Indeed, genes with highly correlated phenotypes likely participate in similar functions, indicating their potential membership in shared complexes or functional pathways^55^. Using protein complex membership data from the Complex Portal, we noted that genes with highly correlated GI profiles in our MCF10A and RPE-1 screens were enriched in known protein complexes (Extended Data Fig. 9a,b)^56^. Notably, correlation analysis of GI profiles from our combinatorial KO screens achieved similar performance to that of DepMap gene essentiality profiles, generated from over 1,000 CRISPR screens^57,58^, in categorizing protein complex members. This similarity is evidenced by comparable areas under the precision-recall curve (PR-AUC), particularly for our RPE-1_integrated dataset (Extended Data Fig. 9a,b). The capacity of our screens to pinpoint protein complex members based on the correlation of GI profiles underscores the robustness of our genetic interaction screening data.

The enrichment of correlated genes in known protein complexes prompted us to assess whether module members might also form direct physical interactions. AlphaFold2-multimer predicted enrichment in direct physical contacts between module members for 12 of the 53 modules of the MCF10A_integrated dataset, 18 of the 52 modules of the RPE1_integrated dataset, and 17 of the 58 modules of the RPE-1/MCF10A_integrated dataset (Extended Data Fig. 9c-e and Supplementary Tables 8,9), supporting a link between genetic module composition and direct physical interactions. By analyzing the correlation of module odds ratio values with AlphaFold2-multimer scores we identified 33, 53, and 49 factors predicted to directly physically interact within modules of the MCF10A_integrated, RPE1_integrated, and RPE-1/MCF10A_integrated datasets, respectively (Extended Data Fig. 10a, upper right quadrants, and Supplementary Table 10). In addition to finding known physical interactors within the same module (*e.g.*, SWSAP1-SPIDR-ZSWIM7, SHLD1-2; Extended Data Fig. 10b,c), AlphaFold2-multimer also identified module members without predicted direct physical interaction that might instead associate through bridging factors either not present in our screens or belonging to other modules (*e.g.*, BRIP1-BRCA1-FANCA, CIP2A-TOPBP1-RHINO, FIGNL1-FIRRM-MCM9; Extended Data Fig. 10d-f). Gene set enrichment analysis (GSEA) for the identified modules assigned 14 modules of the MCF10A_integrated dataset and 22 modules of the RPE-1_intergrated dataset to terms from GO:CC, CORUM, Reactome or KEGG (Extended Data Fig. 11), confirming the functional relationship of components of our modules. Next, module-module interactions were calculated as the mean of all GI z-scores between genes from each cluster (Extended Data Fig. 6c and Supplementary Table 7). Approximately 50% of all modules in the MCF10A_integrated, RPE1_integrated, and MCF10A-RPE1_integrated datasets exhibited strong interactions with at least one other module (Extended Data Fig. 6b, 7b, 8b and Supplementary Table 7), revealing genetic interactions shared among functionally related genes.

Modules of related genes that share genetic interactions are of particular interest as they represent functionally related pathways^3,59^. These modular interactions also enhance confidence in genetic interactions, given that multiple points, representing a single biological function, exhibit similar phenotypes. To increase our confidence in module-module interactions, we applied a stringent criterion, filtering for the top module-module interactions which left us with a reduced set of high-confidence modules in the MCF10A_integrated, RPE1_integrated, and RPE-1/MCF10A_integrated datasets (Fig. 2a,b and Extended Data Fig. 12a,b and 13a,b). Within this refined set of high-confidence module-module interactions, we noted groups of genes associated with known biological pathways. In our RPE-1/MCF10A_integrated dataset, these include genes implicated in the FA pathway *FANCA/B/C/D2/F/G/I/L/BRIP1/UBE2T*), TLS (*REV1/MAD2L2/REV3L*), NHEJ (*XRCC4*/*LIG4*/*NHEJ1* and *CYREN*/*ERCC6L2*), Holliday junction resolution (*SLX4*/*MUS81*/*EME1* and *SLX1A*/*SLX1B*), and mitotic DNA repair (*CIP2A/GEN1/RHNO1*), as well as genes encoding for the RNASEH2, MCM8-MCM9-HROB, PARP1-XRCC1, FANCM-FAAP24, BRCA1-BARD1, ATM-NBN, shieldin (SHLD1-SHLD2), and Shu (SPIDR-ZSWIM7) complexes (Fig. 2a,b). Module-module interactions identified known synthetic lethal interactions, such as those between *BRCA1*, *BARD1* and genes in the PARP1 and FANCM modules^60–62^, as well as suppressor interactions between the *BRCA1/BARD1* module and shieldin-containing modules^28,63–66^ (Fig. 2a,b). Additionally, we identified strong negative interactions between FA genes and the *CIP2A/GEN1/RHNO1* module that have not been previously characterized. We also observed previously unreported negative interactions between the *MCM8/MCM9/HROB* module and the *REV1/MAD2L2/REV3L* module, the RNASEH2 module and the ATM module, the *APEX1/RNASEH1* module and NHEJ modules, and the Shu complex module and the *REV1/MAD2L2/REV3L* module. These novel module-module interactions represent some of the top hits from our screens due to their redundancy in supporting evidence and their high genetic interaction scores, meriting further downstream validation.

**Fig. 2.**
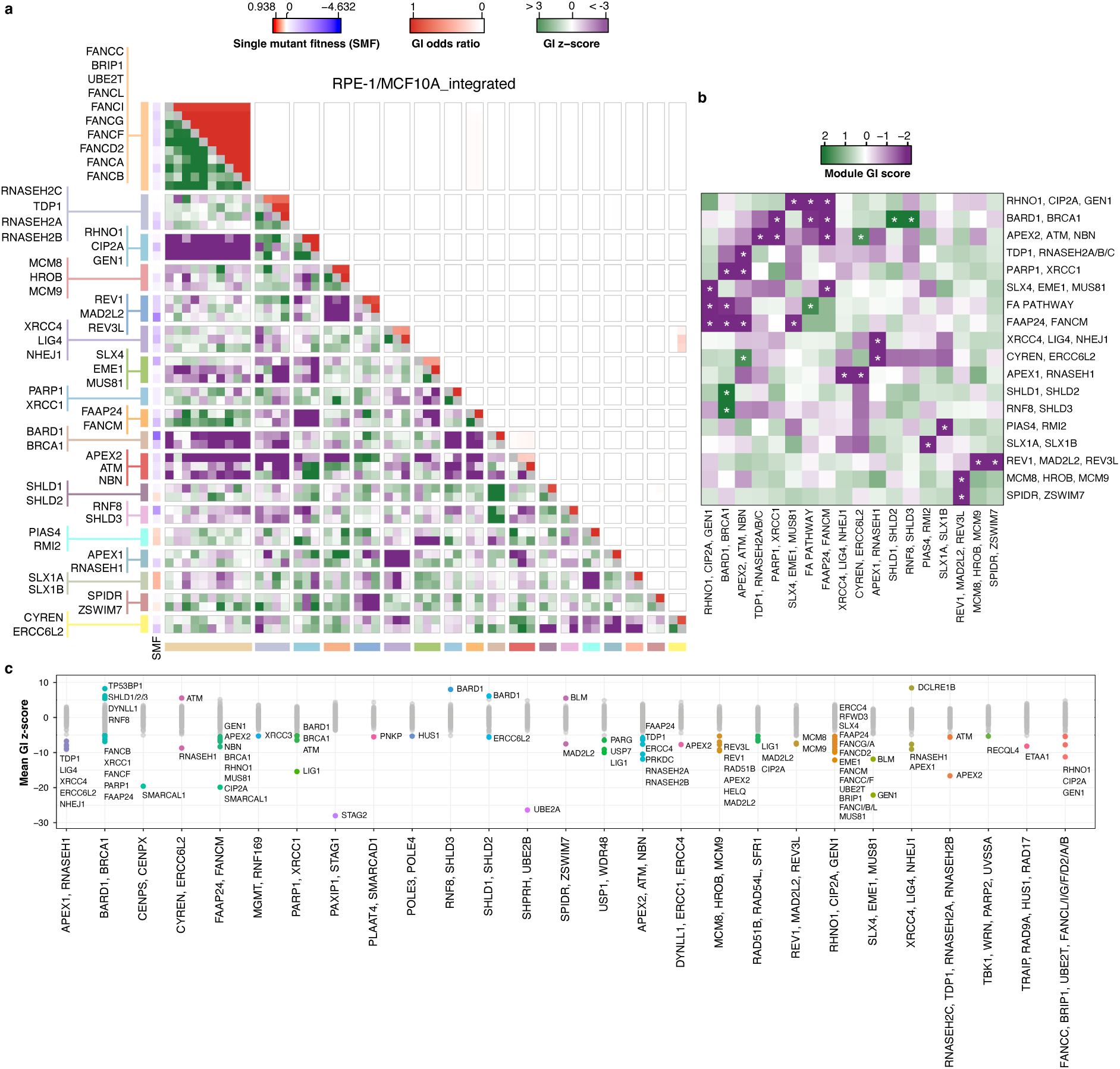
DDR network analysis and GI modules. **a**, Filtered GI map highlighting genes from 18 high-confidence modules from the RPE-1/MCF10A_integrated dataset. The 233-gene library was filtered to include genes involved in top-scoring module-module interactions. The lower heatmap triangle displays integrated GI z-scores (negative in purple, positive in green). The upper heatmap triangle displays cluster odds ratios, representing the frequency with which each two genes cluster together across multiple filtered subsets of the GI dataset (see methods). To the left of the heatmap, 18 GI modules are shown, together with each gene’s single mutant fitness (SMF). Interaction of a gene with itself is depicted as grey in the diagonal. **b**, A reduced heatmap showing the significant module-module interactions used for filtering in panel **a**. Significance is indicated (*), corresponding to a nominal two-sided p-value ≤ 0.1 under a normal approximation. **c**, Analysis of gene-module interactions. Each column represents a module (x-axis), and each point represent a single gene’s mean GI z-score with that module (y-axis). Colored datapoints indicate significant interactions (FDR ≤ 0.1). Modules without significant single gene interactors were omitted.

Finally, leveraging gene modules, we identified individual gene-module interaction patterns (Fig. 2c and Extended Data Fig 12c, 13c). In our RPE-1/MCF10A_integrated dataset, this analysis reaffirmed several interactions described above, such as those between FA genes and *CIP2A/GEN1/RHNO1* and between *REV1/MAD2L2/REV3L* and *MCM8/MCM9/HROB* (Fig. 2c and Supplementary Table 7). We also confirmed the ability of our screens to recapitulate known interactions, such as the synthetic lethal interactions between *GEN1* or *BLM* and the *SLX4/MUS81/EME1* module^39,40^, and between *LIG1* and the *USP1/WDR48* module^67^ (Fig. 2c). Additionally, we uncovered robust interactions between single genes and modules that were not apparent from module-level analysis. These interactions were missed in the module-module analysis because some genes did not cluster into defined modules, and module-level aggregation can obscure gene-specific interactions. Among those, we identified strong negative interactions of *SMARCAL1* with *FANCM/FAAP24* and *CENPS/CENPX*, consistent with recent reports^67,68^, and positive interactions between *BLM* and *SPIDR/ZSWIM7*, as previously reported in yeast and mouse models^48,69^, as well as between *DCLRE1B* and *XRCC4/LIG4/NHEJ1* (Fig. 2c). These robust modular interactions, representing clusters of shared interactions between genes with similar annotations, provide compelling evidence for epistasis between DDR functional pathways.

### Early time point analysis reveals genetic interactions with essential genes

While our DDR genetic interaction library primarily targets non-essential DNA repair genes, we deliberately incorporated a subset of core essential factors to enable the identification of suppressor interactions. Furthermore, we included an early time point (T9) to assess whether our screens could detect negative genetic interactions with essential genes before strong fitness defects at the final time point (T17/T18) obscured their pairwise interaction signatures. Comparison of high-confidence negative interactions (FDR < 0.1) across time points revealed that more than 50% of hits were unique to the early time point in both MCF10A and RPE-1 cells (Extended Data Fig. 14a,c and Supplementary Table 5). Notably, these early time point-specific interactions were enriched for gene pairs containing essential genes relative to the final time point, with ∼41% and ∼64% of T9-exclusive interactions containing at least one essential gene in MCF10A and RPE-1 cells, respectively (Extended Data Fig. 14b,d and Supplementary Tables 11, 12).

Analysis of early time point-exclusive interactions containing essential genes uncovered module-level relationships that were not apparent at the final time point (Supplementary Table 13). For example, we observed negative genetic interactions between a module containing the essential flap endonuclease *FEN1* and the DNA ligase *LIG1* and the *USP1/WDR48* PCNA deubiquitylation module (Extended Data Fig. 14e)^70–73^, consistent with a recent report from a combinatorial CRISPR interference screen^67^. In addition, we identified interactions between the essential *RAD17/HUS1/RAD9A* module that senses DNA damage and promotes ATR activation and the *BOD1L1/POLE3/POLE4/ETAA1* replication stress response module (Extended Data Fig. 14f)^74–78^. As expected, by the final time point, strong fitness defects associated with essential gene perturbations became saturated, obscuring pairwise interaction effects and diminishing meaningful clustering of essential genes. In particular, *FEN1* clustered with other essential genes rather than with its functional partner *LIG1*, and components of the *RAD17/HUS1/RAD9A* module no longer clustered together (Extended Data Fig. 14e,f). In addition, the *BOD1L1/POLE3/POLE4/ETAA1* module identified at T9 was no longer preserved at T18, when only *POLE3* and *POLE4* remained clustered together (Extended Data Fig. 14f). This pattern suggests that shared strong negative genetic interactions with the *RAD17/HUS1/RAD9A* module drove co-clustering of *BOD1L1, POLE3/POLE4*, and *ETAA1* at the early time point, with their separation at the final time point likely reflecting differential functions during DNA replication^74–77^.

Additionally, we identified significant gene-module interactions involving essential genes at the early time point (Extended Data Fig. 14g,h and Supplementary Table 13). These include genetic interactions between essential DNA damage-sensing genes that promote ATR activation (*HUS1* and *RAD17*) and *ATM* in both RPE-1 and MCF10A datasets, consistent with parallel modes of checkpoint activation and DSB signaling provided by ATR and ATM^79,80^. We also observed interactions between essential homologous recombination (HR) modules (*MRE11/RAD50* or *XRCC3/RAD51D/XRCC2*) and single-strand break repair genes (*PARP1, XRCC1*) in the RPE-1 dataset, in line with previously reported PARP inhibitor sensitivity upon loss of these HR factors^81,82^. Collectively, these results demonstrate that incorporating an early time point into combinatorial CRISPR KO screens enables detection of negative genetic interactions involving essential genes.

### Validation studies confirm the phenotype of DDR genetic interactions

To confirm the phenotype of genetic interactions identified in our screens, we selected a subset of hits from our RPE-1_T18 and MCF10A_T17 datasets for validation using mid-throughput assays. In total, we included 22 negative and 16 positive genetic interactions from the MCF10A dataset and 20 negative and 21 positive genetic interactions from the RPE-1 dataset, prioritizing interactions based on 1) therapeutic relevance (disorder- and cancer-relevant genetic interactions from GENIE- and ClinVar-based analysis), 2) module connectivity (gene pairs from significant module-module and gene-module interactions, including cases where individual gene pairs did not meet the FDR < 0.1 threshold), and 3) individual GI z-scores (top-ranking interactions involving genes of interest, including interactions with FDR > 0.1).

To validate the genetic interactions of interest, we employed cell growth-based two-color fluorescence competition assays. In these assays, the growth of individual samples carrying in4mer constructs targeting either a single gene and a control non-essential gene or a pair of genes was measured in a mixed population alongside cells carrying control gRNAs targeting non-essential genes (Extended Data Fig. 15a). GI scores were calculated by subtracting the expected fitness of double mutants, derived from the sum of single mutant fitness phenotypes, from the observed double mutant fitness (Extended Data Fig. 15b). These phenotypes were recorded across several timepoints during our validation studies.

The GI scores obtained from our validation experiments showed a strong concordance with the GI z-scores derived from our screens in both RPE-1 and MCF10A cells (Fig. 3a), further highlighting the robustness of our genetic interaction screen analyses. In total, we confirmed the phenotype of 36 negative and 23 positive genetic interactions (Fig. 3b-d, Extended Data Fig. 15c, 16a, and Supplementary Table 14). Four negative interactions in RPE-1 cells that did not validate in the dKO format were instead confirmed using single gene KO combined with a small-molecule inhibitor targeting the protein encoded by the second gene, in line with recent studies^83,84^, providing additional support for screen accuracy (Extended Data Fig. 16b-e). In our validation, we confirmed the existence of several interaction sets that span multiple clusters. For mechanistic interrogation, we chose to prioritize genetic interaction sets based on the clarity of the associated modules and the strength of the validated genetic interactions. Thus, we selected for further characterization the interactions between 1) the *REV1/MAD2L2/REV3L* module and the *MCM8/MCM9/HROB* module, 2) the FA gene module and the *CIP2A/GEN1/RHNO1* module, 3) *SMARCAL1* and the *FANCM/FAAP24* and *CENPS/CENPX* modules, as well as 4) *DCLRE1B* and the *XRCC4/LIG4/NHEJ1* module.

**Fig. 3.**
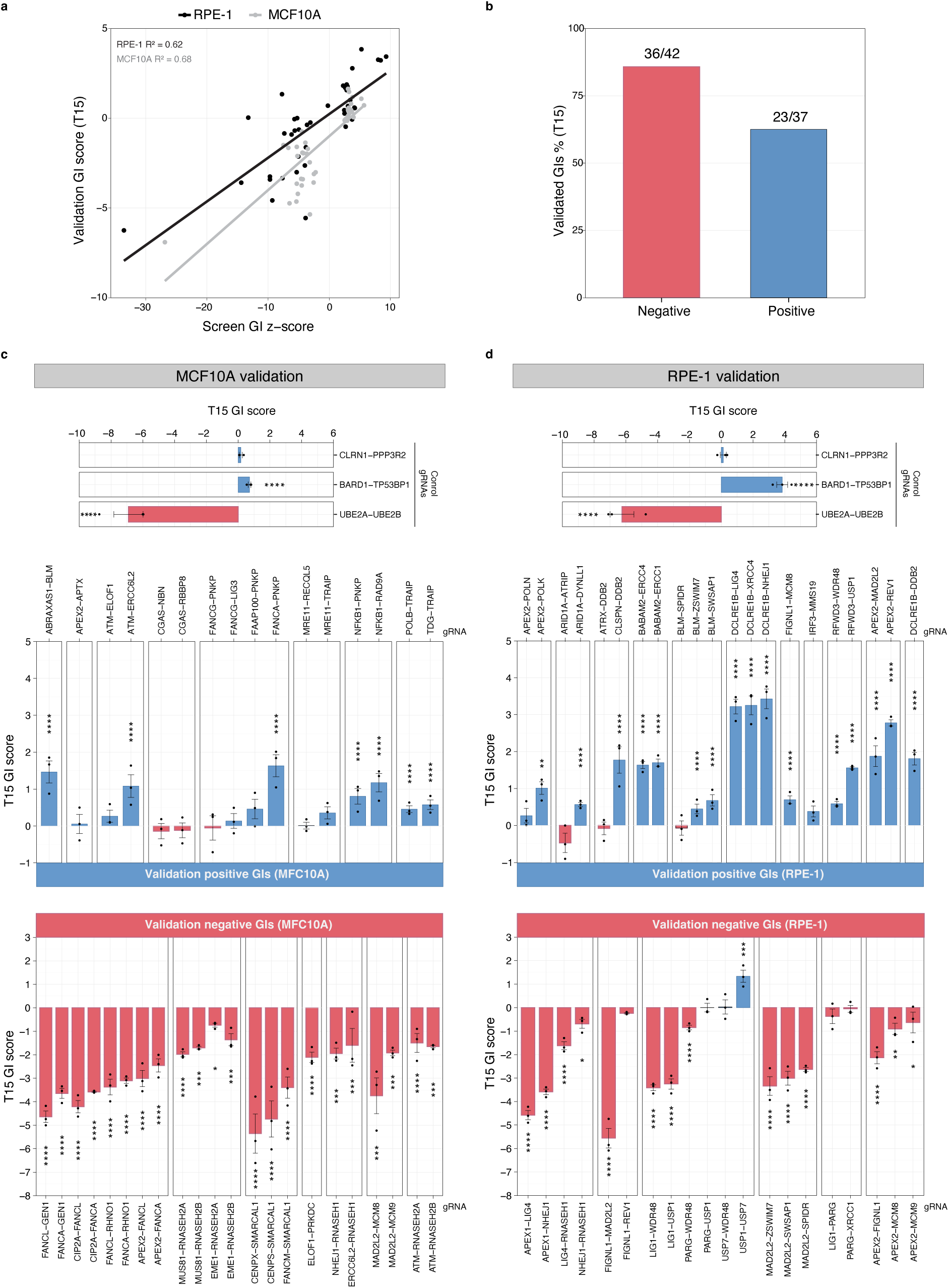
Arrayed validation of GIs among DDR genes. **a**, Comparison of validation GI scores of individual genetic interactions obtained by two-color competition assays at day 15 (T15) in MCF10A and RPE-1 *TP53^-/-^* cells and GI z-scores for the corresponding interactions obtained from the Cas12a-based GI screens at T17/T18. **b**, Combined validation rates for tested GIs in MCF10A and RPE-1 *TP53^-/-^* cells. The percent of screen hits that have a validation p-value ≤ 0.05 are shown for all validation pairs. **c**, **d**, Validation GI scores at T15 in MCF10A and RPE-1 *TP53 ^-/-^* cells, respectively, for reference gene pairs (top), including a control pair (*CLRN1-PPP3R2*), a positive interaction pair (*BARD1-TP53BP1*), and a negative interaction pair (*UBE2A-UBE2B*), and for genetic interactions of interest (bottom). Positive and negative genetic interactions are shown in blue and red, respectively. Significant GIs are called based on the difference between expected and observed log fold changes (LFCs) for dual-gene targeting gRNAs at T15 using two-way ANOVA. Points represent the mean of three technical replicates and bars show the mean ± SEM (n = 3 biological replicates). Statistically significant GI scores are indicated. Corresponding single gene effects, as well as expected and observed LFCs used to calculate validation GI scores at T15, are shown in Extended Data Fig. 15c and 16a for MCF10A and RPE-1 *TP53^-/-^* datasets, respectively. **** = p-value ≤ 0.0001, *** = p-value ≤ 0.001, ** = p-value ≤ 0.01, * = p-value ≤ 0.05.

### MCM8/9-deficient cells are dependent on the REV1-Pol ζ complex

In our RPE-1/MCF10A_integrated dataset we identified a functional module containing *HROB, MCM8,* and *MCM9* that showed strong negative interactions with a module consisting of *MAD2L2, REV1,* and *REV3L* (Fig. 2a,b and 4a). Further, *MCM8* and *MCM9* ranked as the top two negative interactors with *MAD2L2*, *REV1*, and *REV3L* (Supplementary Table 3). To confirm these findings, we selected gRNAs that efficiently disrupted *MCM8*, *MCM9*, and *MAD2L2* (Extended Data Table 1 and Extended Data Fig. 17a), and conducted two-color competition assays using MCF10A cells deficient for these genes (Fig. 3c and 4b). In these assays, disruption of *MAD2L2* alone caused a mild reduction in cell viability, while additional loss of MCM8 or MCM9 caused a synergistic reduction in cell fitness, confirming the negative interactions between *MAD2L2* and *MCM8/9* (Fig. 4b).

**Fig. 4.**
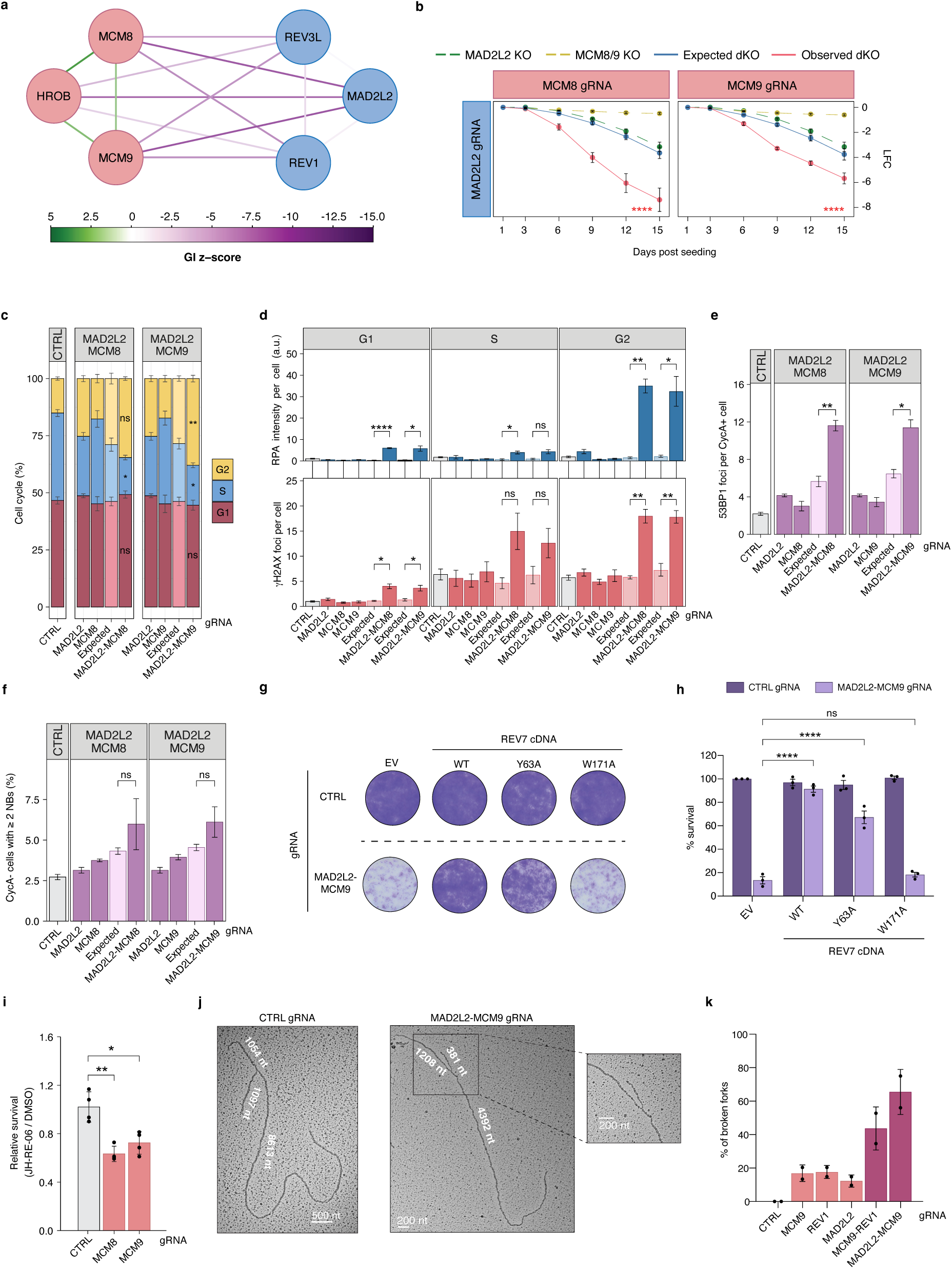
Dependency of MCM8/9-HROB complex mutant cells on REV1-Pol ζ. **a**, GI network for the *HROB*, *MCM8*, *MCM9* and *MAD2L2*, *REV1*, *REV3L* modules from the RPE-1/MCF10A_integrated heatmap. GI z-scores < 5 and > −15 are shown. Node colors are according to the module coloring scheme in Fig. 2a. **b**-**k**, Cells were targeted with the indicated in4mer gRNAs. **b**, Validation of *MAD2L2* interactions with *MCM8/9* by two-color competition assay in MCF10A cells. Points represent mean ± SEM (n = 3 biological replicates). Statistical significance was determined by two-way ANOVA comparing expected and observed values at T15. **c**-**f**, For each interaction, values are shown for individual single KOs, the dKO (observed), and the corresponding expected values for each dKO, calculated from single KO phenotypes. Expected values are plotted immediately to the left of their matched observed dKO values for each gene pair. Significant deviations between observed and expected values are indicated. Bars represent the mean ± SEM (n = 3 biological replicates), unless stated otherwise. Statistical significance was determined by unpaired t test. **c**, Cell cycle profiles for the *MAD2L2* interactions with *MCM8*/*9* in MCF10A cells. Cell cycle phases were determined based on PCNA and DAPI staining. Significant deviations of observed from corresponding expected profiles are indicated for each cell cycle stage. Bars represent the mean ± SEM of six independent experiments (3 biological replicates and 2 independent imaging experiments per replicate). **d**, Quantification of integrated RPA32 foci intensity (in arbitrary units, a.u.) (top) and γH2AX foci (bottom) in MCF10A cells across cell cycle phases (G1, S, and G2), which were distinguished by PCNA and DAPI staining. **e**, **f**, Quantification of 53BP1 foci in cyclin A-positive (CycA+, S/G2) and 53BP1 nuclear bodies (NBs) in cyclin A-negative (CycA-, G1) MCF10A cells, respectively. **g**, **h**, Representative survival assay and quantification, respectively, of control (CTRL) and *MAD2L2-MCM9* dKO MCF10A cells complemented with either empty vector (EV), wild-type (WT), or mutant (Y63A or W171A) REV7 cDNA constructs. Bars represent the mean percentage of survival (± SEM) relative to the CTRL sample complemented with EV cDNA construct (n = 3 biological replicates). Statistical significance was determined by one-way ANOVA. **i**, Sensitivity of MCM8/9-deficient MCF10A cells to the JH-RE-06 inhibitor (1.25 μM). Relative survival is calculated as survival after JH-RE-06 treatment divided by survival after DMSO treatment for each sample. Bars represent the mean ± SEM (n = 4 biological replicates). Statistical significance was determined by unpaired t test**. j**, Representative images of a normal (left) and an asymmetric (broken; right) replication fork detected by EM in CTRL and *MAD2L2-MCM9* dKO MCF10A cells, respectively. The length of the replication fork branches is indicated in nucleotides (nt). Scale bars are shown in nt. **k**, Quantification of broken replication forks detected by EM in *MCM9-REV1* and *MAD2L2-MCM9* dKO MCF10A cells and single KO cells. Bars represent the mean percentage of broken forks (± SD) relative to the total number of replication intermediates (RIs) for each sample (n = 2 biological replicates). At least 41 RIs were examined per sample per experiment. **** = p-value ≤ 0.0001, *** = p-value ≤ 0.001, ** = p-value ≤ 0.01, * = p-value ≤ 0.05.

MCM8 and MCM9 form a complex with HROB that operates in the postsynaptic stages of HR after RAD51 loading and displacement loop (D-loop) formation^41,85^. At this stage, HROB stimulates the helicase activity of the MCM8-9 complex, possibly enabling D-loop extension and migration^85^. MCM8-9 and HROB have also been implicated in promoting replication fork progression in response to replication stress^85,86^. REV3L (Pol ζ catalytic subunit), REV7 (accessory Pol ζ subunit encoded by *MAD2L2),* and REV1 function primarily in TLS as part of the REV1-Pol ζ complex^87^. To elucidate the relationship between these DDR complexes, we conducted cell cycle analyses on MCM8/9-deficient cells upon disruption of *MAD2L2*. These studies revealed that the simultaneous loss of REV7 and MCM8 or MCM9 led to an increase in G2-phase cells greater than expected from the phenotype of the single gene disruptions (Fig. 4c). This increase in G2-phase cells coincided with a decrease in S-phase cells, suggesting that cells deficient in both REV7 and MCM8/9 can progress through S-phase but are arrested in G2 (Fig. 4c), presumably due to the accumulation of DNA lesions following DNA replication. Accordingly, simultaneous disruption of *MAD2L2* and *MCM8/9* resulted in increased γH2AX focus formation and RPA staining, suggestive of ssDNA accumulation (Fig. 4d and Extended Data Fig. 18a,b). Furthermore, simultaneous loss of REV7 and MCM8/9 caused a ∼2-fold increase in 53BP1 foci, a DSB marker, in S/G2 cells (cyclin A-positive) (Fig. 4e and Extended Data Fig. 18c), and a mild, albeit not significant, increase in 53BP1 nuclear bodies, a marker of replication stress, in G1 cells (cyclin A-negative) (Fig. 4f), as compared to expected values.

Besides promoting TLS as part of Pol ζ, REV7 also acts to counteract DSB end resection as a component of the shieldin complex^28,63,64,66,88,89^. Interestingly, our screens did not identify negative interactions for *MCM8, MCM9*, or *HROB* with other components of the shieldin complex (*i.e.*, *SHLD1, SHLD2,* and *SHLD*3), suggesting that the loss of REV7’s functions in the shieldin complex is not responsible for the synthetic lethal phenotype observed in HROB- and MCM8/9-deficient cells. To directly test whether the observed synthetic lethality reflects impaired REV1-Pol ζ function rather than shieldin dysfunction, we performed complementation experiments in MCF10A cells deficient for both REV7 and MCM9. Cells were reconstituted with either wild-type REV7 or separation-of-function mutants: Y63A, which disrupts SHLD2 binding and partially impairs REV3L interaction, or W171A, which selectively impairs REV3L binding while preserving shieldin function^90,91^. The W171A mutant failed to rescue viability, phenocopying complete REV7 loss, whereas Y63A provided partial rescue (Fig. 4g,h and Extended Data Fig. 19a). These results confirm that the observed synthetic lethal interaction depends primarily on REV7’s role within Pol ζ rather than its shieldin-associated NHEJ function. Consistent with this conclusion, pharmacological inhibition of TLS recapitulated the genetic phenotype. Indeed, treatment with JH-RE-06, a small molecule that blocks REV1-Pol ζ complex formation by targeting REV1 while leaving REV7-shieldin interactions intact^92^, reduced viability by ∼37% in MCM8-deficient cells and ∼28% in MCM9-deficient cells compared to wild-type control (Fig. 4i).

We next investigated the mechanistic basis of this synthetic lethality. REV1-Pol ζ operates at replication forks to promote bypass of DNA lesions and post-replicative refilling of ssDNA gaps generated by the PRIMPOL primase/polymerase^93–95^. We previously showed that loss or inhibition of REV1-Pol ζ reduces the viability of HR-deficient breast cancer cells carrying *BRCA1* mutations, and that this lethality can be suppressed by preventing ssDNA gap accumulation through loss of PRIMPOL^93^. We observed similar effects in MCF10A cells, where combined loss of BRCA1 and REV7 or REV1 induced a loss of cell fitness that could be rescued by PRIMPOL depletion (Extended Data Fig. 19b,c). However, PRIMPOL loss did not suppress the cell fitness defect caused by combined loss of MCM9 and REV7 or REV1, indicating that these negative genetic interactions are PRIMPOL-independent (Extended Data Fig. 19d,e).

To further characterize the replication defects underlying these distinct synthetic lethal interactions, we employed electron microscopy (EM) to visualize replication intermediates. In previous studies, we observed that BRCA1-deficient cells with impaired REV1-Pol ζ function accumulated large (>1,000 nt-long) ssDNA regions at replication forks in a PRIMPOL-dependent manner^93^. In contrast, combined loss of MCM9 and REV7 or REV1 did not lead to the accumulation of large ssDNA gaps (Extended Data Fig. 19f). Instead, these cells exhibited a 2-to-3-fold increase in asymmetric replication fork intermediates in which all three fork branches were of unequal length, indicative of replication fork collapse^96,97^ (Fig. 4j,k and Extended Data Fig. 19g-j). Together, these findings highlight a novel dependency of HROB- and MCM8/9-deficient cells on the TLS polymerase complex REV1-Pol ζ and implicate replication fork instability as the underlying mechanism.

### Cells deficient in Fanconi anemia genes rely on mitotic DNA repair

In our screens, we identified a module consisting of 10 FA genes that shared strong negative interactions with a module consisting of *CIP2A, GEN1,* and *RHNO1* (Fig. 2a,b and 5a). Interestingly, analysis of DepMap data revealed that *CIP2A, GEN1,* and *RHNO1* have highly correlated essentiality scores, and cluster together in a small network of top correlated partners (Extended Data Fig. 20a), providing evidence for a shared biological function that is essential in FA-deficient backgrounds.

**Fig. 5.**
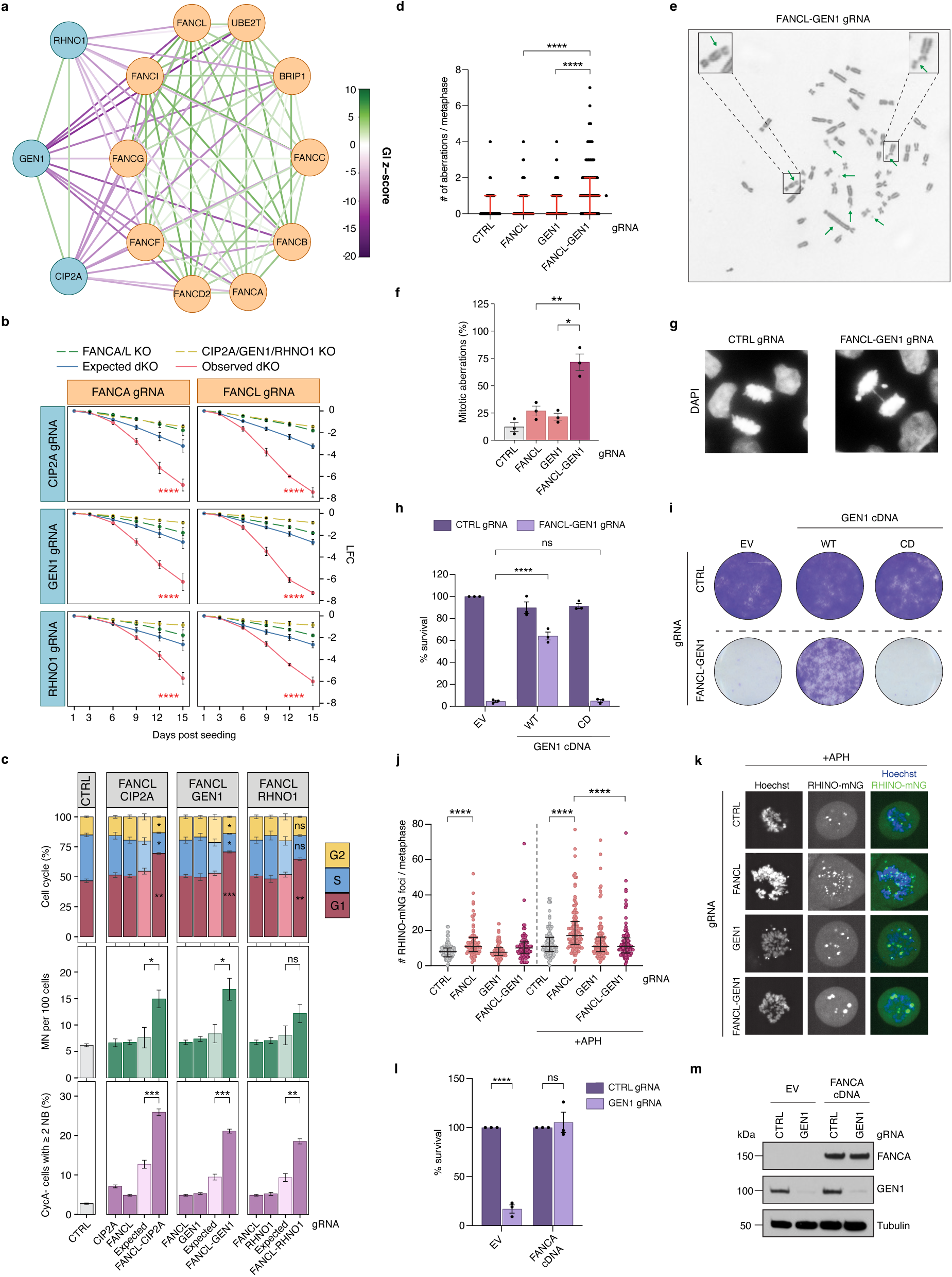
Negative interactions between the Fanconi anemia gene module and the mitotic repair genes *CIP2A*, *GEN1,* and *RHNO1*. **a**, GI network for genes in the FA module and the *CIP2A*/*GEN1*/*RHNO1* module from the RPE-1/MCF10A_integrated heatmap. GI z-scores < 10 and > −20 are shown. Node colors are according to the module coloring scheme in Fig. 2a. **b**-**m**, Cells were targeted with the indicated in4mer gRNAs, unless stated otherwise. **b**, Validation of *FANCA/L* interactions with *CIP2A, GEN1,* and *RHNO1* by two-color competition assay in MCF10A cells. Points represent the mean ± SEM (n = 3 biological replicates). Statistical significance was determined by two-way ANOVA comparing expected and observed values at T15. **c**, Cell cycle profiles (top), quantification of micronuclei (MN) (middle), and 53BP1 nuclear bodies (NBs) in cyclin A-negative (CycA-, G1) MCF10A cells (bottom) for the indicated interactions. G1, S, and G2 phases for cell cycle profiles were determined based on PCNA and DAPI staining. For each interaction, values are shown for individual single KOs, the dKO (observed), and the corresponding expected values for each dKO, calculated from single KO phenotypes. Expected values are plotted immediately to the left of their matched observed dKO values for each gene pair. Significant deviations between observed and expected values are indicated. For cell cycle profiles (top), bars represent the mean ± SEM of six independent experiments (3 biological replicates and 2 independent imaging experiments per replicate). For MN (middle) and 53BP1 NBs (bottom), bars represent the mean ± SEM (n = 3 biological replicates). Statistical significance was determined by unpaired t test. **d**, Quantification of metaphase aberrations detected by metaphase spreads in RPE-1 *TP53^-/-^*cells. Scatter plot shows individual values (n = 2 biological replicates). At least 50 metaphases were examined per sample per experiment. Bars represent the median and interquartile range. Statistical significance was determined by Mann-Whitney test. **e**, Representative metaphase image from *FANCL-GEN1* dKO RPE-1 *TP53^-/-^* cells quantified in panel **d**. Metaphase aberrations, such as chromosome gaps (*e.g.*, top left zoom section), chromosome breaks (*e.g.*, top right zoom section), chromatid breaks, and a dicentric chromosome are indicated by green arrows. **f,** Quantification of spontaneous mitotic aberrations (lagging chromosomes and anaphase bridges) in MCF10A cells. Bars represent the mean percentage of anaphases with mitotic aberrations (± SEM) relative to all anaphases scored per sample (n = 3 biological replicates). At least 30 anaphase cells were scored per sample per experiment. Statistical significance was determined by one-way ANOVA. **g**, Representative images of anaphases observed in control (CTRL) and *FANCL-GEN1* dKO MCF10A cells quantified in panel **f**. **h**, **i**, Quantification of survival assays and representative survival, respectively, of control (CTRL) and *FANCL-GEN1* dKO MCF10A cells complemented with empty vector (EV), wild-type (WT) GEN1, or catalytic-dead (CD) GEN1 cDNA constructs. *FANCL* and *GEN1* were targeted using a dual-gRNA Cas12a construct. Bars represent the mean percentage of survival (± SEM) relative to the CTRL sample complemented with EV cDNA construct (n = 3 biological replicates). Statistical significance was determined by one-way ANOVA. **j**, Quantification of mitotic RHINO-mNeonGreen (RHINO-mNG) foci in live-cell imaging experiments in U2OS cells with and without aphidicolin (APH) treatment (see Extended Data Fig. 20m for a schematic of the imaging protocol). Scatter plot shows individual values (n = 3 biological replicates). At least 24 mitotic cells were examined per sample per experiment. Bars represent the median and interquartile range. Statistical significance was determined by Mann-Whitney test. **k**, Representative images of mitotic RHINO-mNG foci in APH-treated cells quantified in panel **j**. **l**, Quantification of survival assays for OHSU-SCC-974 cells complemented with either EV or wild-type FANCA cDNA constructs upon knockout of *GEN1*. Bars represent the mean ± SEM (n = 3 biological replicates). The mean percentage of survival in GEN1-deficient samples was normalized to the corresponding CTRL samples. Statistical significance was determined by one-way ANOVA. **m**, Western blot showing FANCA and GEN1 levels in OHSU-SCC-974 cells examined in the survival assays in panel **l**. Tubulin was used as a loading control. **** = p-value ≤ 0.0001, *** = p-value ≤ 0.001, ** = p-value ≤ 0.01, * = p-value ≤ 0.05.

CIP2A, GEN1, and RHINO (encoded by *RHNO1)* all function in mitotic DNA repair processes^40,98–102^. CIP2A is a member of a complex that tethers unrepaired DSBs in mitosis, allowing for the proper segregation of acentric chromosome fragments during cell division^100,101,103^. RHINO, a 9-1-1 and TOPBP1 interacting partner involved in ATR activation, also plays a role in DSB repair during mitosis by promoting microhomology-mediated end joining (MMEJ) through its association with the DNA polymerase Pol θ^102,104^. Similarly, GEN1, a structure-specific endonuclease that functions in mitosis after nuclear envelope breakdown, promotes DSB repair by resolving Holliday junctions formed during HR^98,99,105^. GEN1 also mediates common fragile site expression, allowing mitotic DNA synthesis at difficult-to-replicate genomic sites^106^. The top genetic interactors of *CIP2A*, *GEN1*, and *RHNO1* are members of a module comprised of genes mutated in FA, a genetic disorder characterized by bone marrow failure and cancer predisposition^107^. Mutations in 23 genes involved in interstrand cross-link (ICL) repair, HR, TLS, and replication fork stability have been shown to cause FA^107^.

To validate the synthetic lethal interactions within these modules, we first disrupted two FA genes (*FANCA* and *FANCL*) in MCF10A cells using gRNAs (Fig. 3c and 5b). While FANCA/L-deficient MCF10A cells exhibited moderate viability defects, additional disruption of *CIP2A, GEN1,* or *RHNO1* caused a pronounced reduction in cell fitness, confirming the strong negative genetic interaction observed in our screens (Fig. 5b, Extended Data Fig. 17b, and Extended Data Table 1). Similarly, loss of the RHINO interactor Pol θ reduced the fitness of FANCL-deficient MCF10A and RPE-1 cells (Extended Data Fig. 21a,b). Additionally, disruption of *APEX2*, an endonuclease that promotes the repair of blocked 3’ DNA ends generated during the processing of genomic ribonucleotides^42^ and stimulates MMEJ^108^, also reduced viability in FANCA/L-deficient MCF10A cells (Fig. 3c). Notably, *APEX2* and *POLQ* clustered with *CIP2A, GEN1,* and *RHNO1* in the MCF10A_integrated and RPE-1_integrated datasets, respectively (Extended Data Fig. 12a and 13a). Loss of Pol θ, APEX2, and CIP2A has been previously shown to cause lethality in BRCA1/2-deficient cells^42,100,109–111^. In contrast, GEN1 loss did not significantly affect the growth of BRCA1-deficient RPE-1 cells (Extended Data Fig. 20b-f), consistent with previous studies that did not identify *GEN1* among *BRCA1/2* synthetic lethal partners^100^. These findings indicate that GEN1 represents a synthetic lethal partner specific to FA-deficient, but not BRCA1/2-deficient, cells.

Disruption of *FANCL* in combination with *CIP2A, GEN1,* or *RHNO1* resulted in the accumulation of MCF10A cells in G1 at a rate significantly greater than expected from their single-mutant phenotypes (Fig. 5c). Such G1 accumulation can arise from unresolved DNA damage in the previous cell cycle that has been transmitted through mitosis. Consistent with this hypothesis, MCF10A cells deficient for FANCL and CIP2A or GEN1 showed increased accumulation of micronuclei and 53BP1 nuclear bodies in G1, indicative of unrepaired DNA lesions carried over from mitosis (Fig. 5c and Extended Data Fig. 18d,e). In contrast to loss of CIP2A, RHINO, or GEN1, APEX2 loss in FANCL-deficient MCF10A cells caused only a modest increase of 53BP1 nuclear bodies in G1 and did not induce G1 arrest (Extended Data Fig. 21c,e). Instead, it resulted in the accumulation of 53BP1 foci during S/G2 (Extended Data 18f and 21d), suggesting that the mechanism underlying the synthetic lethality with FA genes differs between *APEX2* and the above interactors.

Combined loss of FANCL and GEN1 led to a ∼3-fold increase in mitotic abnormalities and a significant increase in metaphase aberrations as compared to single gene deficiencies (Fig. 5d-g and Extended Data Fig. 20g). Given the established role of GEN1 in processing DNA replication and recombination intermediates during mitosis, we then investigated whether its catalytic activity is required for the survival of FA-deficient cells. To this end, we complemented FANCL/GEN1-deficient MCF10A cells with either wild-type GEN1 or a catalytically inactive mutant (E134A/E136A)^105^. Complementation with wild-type, but not catalytically inactive, GEN1 rescued cell viability (Fig. 5h,i and Extended Data Fig. 20h), indicating that endonucleolytic processing of branched DNA intermediates by GEN1 is essential for the survival of FA-deficient cells.

Previous studies have shown that RHINO localizes to DSBs in mitosis, where it promotes Pol θ recruitment to facilitate repair by MMEJ^102,112^. To investigate whether loss of FA genes promotes accumulation of RHINO and Pol θ foci during mitosis, we endogenously tagged RHINO with an mNeonGreen fluorescent tag in U2OS cells already expressing endogenously tagged Halo-Pol θ^102^, as both proteins are difficult to detect with available antibodies (Extended Data Fig. 20i-k). *FANCL* disruption in these cells led to enhanced formation of mitotic RHINO foci under unperturbed conditions, which was further exacerbated upon aphidicolin treatment (Fig. 5j,k, Extended Data Fig. 20l,m, and Extended Data Table 1). Similarly, FANCL-deficient cells displayed increased accumulation of Pol θ foci following aphidicolin treatment (Extended Data Fig. 20m-o). Notably, GEN1 loss suppressed the elevated formation of both RHINO and Pol θ mitotic foci in FANCL-deficient cells (Fig. 5j,k and Extended Data Fig. 20n,o), suggesting that GEN1 can generate mitotic DSBs that are subsequently recognized by RHINO and Pol θ. FANCL-deficient MCF10A cells also exhibited accumulation of CIP2A foci in mitosis, consistent with CIP2A’s role in recognizing mitotic DNA lesions (Extended Data Fig. 20p,q). However, unlike RHINO and Pol θ foci, CIP2A foci formation was not suppressed by GEN1 loss, suggesting that GEN1 might act downstream of CIP2A (Extended Data Fig. 20p). Together, these data support a model in which unresolved DNA lesions accumulate in mitosis in FA-deficient cells, where they are processed by GEN1 to generate DSBs that are then recognized by the RHINO-Pol θ machinery for MMEJ-mediated repair.

To determine the relevance of the synthetic lethal interaction between FA genes and *GEN1* in the context of cancer, we obtained the head and neck squamous cell carcinoma cell line OHSU-SCC-974, which carries biallelic *FANCA* loss^113,114^. Disruption of *GEN1* caused loss of viability in OHSU-SCC-974 cells, which was suppressed by complementation with *FANCA* cDNA^114^, confirming that the observed effect is dependent on FANCA deficiency (Fig. 5l,m). These findings suggest that the synthetic lethal interaction between FA genes and *GEN1* is also conserved in cancer cells with FA pathway deficiency.

### SMARCAL1-deficient cells depend on FANCM and its associated factors

Our screens also identified two modules comprised of *CENPS, CENPX* and *FANCM, FAAP24* that interact with *SMARCAL1* (Fig. 2c and 6a), consistent with recent findings from other groups^67,68^. SMARCAL1 and FANCM are DNA translocases that remodel branched DNA structures, such as stalled replication forks^115–124^. Interestingly, CENPS, CENPX, and FAAP24 are physical interacting partners of FANCM that stimulate the DNA translocase activity of FANCM or its recruitment to stalled forks^123,125,126^. The negative interactions of *CENPS, CENPX,* and *FANCM* with *SMARCAL1* were all confirmed with two-color competition assays using gRNAs targeting these genes in MCF10A cells (Fig. 6b, Extended Data Fig. 17c, and Extended Data Table 1). Interestingly, while loss of FANCM alone markedly reduced cell fitness, individual loss of CENPS, CENPX, or SMARCAL1 had no apparent effect on cell proliferation (Fig. 6b). Cell cycle analysis did not show significant changes in cell cycle distribution in MCF10A cells deficient for SMARCAL1 and CENPX at day 7 after gRNA selection, while a significant increase in G1-phase cells and reduction in S-phase cells was observed in cells deficient for SMARCAL1 and FANCM at a similar time point (Fig. 6c). Greater than additive phenotypes were also observed for micronuclei and 53BP1 focus formation in cells deficient for SMARCAL1 and FANCM (Fig. 6d,e and Extended Data Fig. 18g,h). 53BP1 nuclear body formation was also elevated above expected levels in cells deficient for SMARCAL1 and FANCM (Fig. 6f). Additionally, cell cycle-dependent accumulation of RPA in G2 was noted in cells deficient for SMARCAL1 and FANCM (Fig. 6g and Extended Data Fig. 18i), confirming the accumulation of replication stress in these cells. Similar phenotypes were not observed in MCF10A cells deficient for SMARCAL1 and CENPX at day 7 after gRNA selection (Fig. 6d-g). Nonetheless, cells deficient for SMARCAL1 and CENPX displayed accumulation of mitotic aberrations similar to cells lacking SMARCAL1 and FANCM (Fig. 6h,i and Extended Data Fig. 22a). Interestingly, loss of FANCM alone also caused a pronounced increase in mitotic aberrations (Fig. 6i). Consistent with previous studies^67,68^, analysis of metaphase spreads from MCF10A cells deficient in SMARCAL1 and either CENPX, FAAP24, or FANCM revealed chromosome breaks and cohesion defects (Fig. 6j,k and Extended Data Fig. 22b-d).

**Fig. 6.**
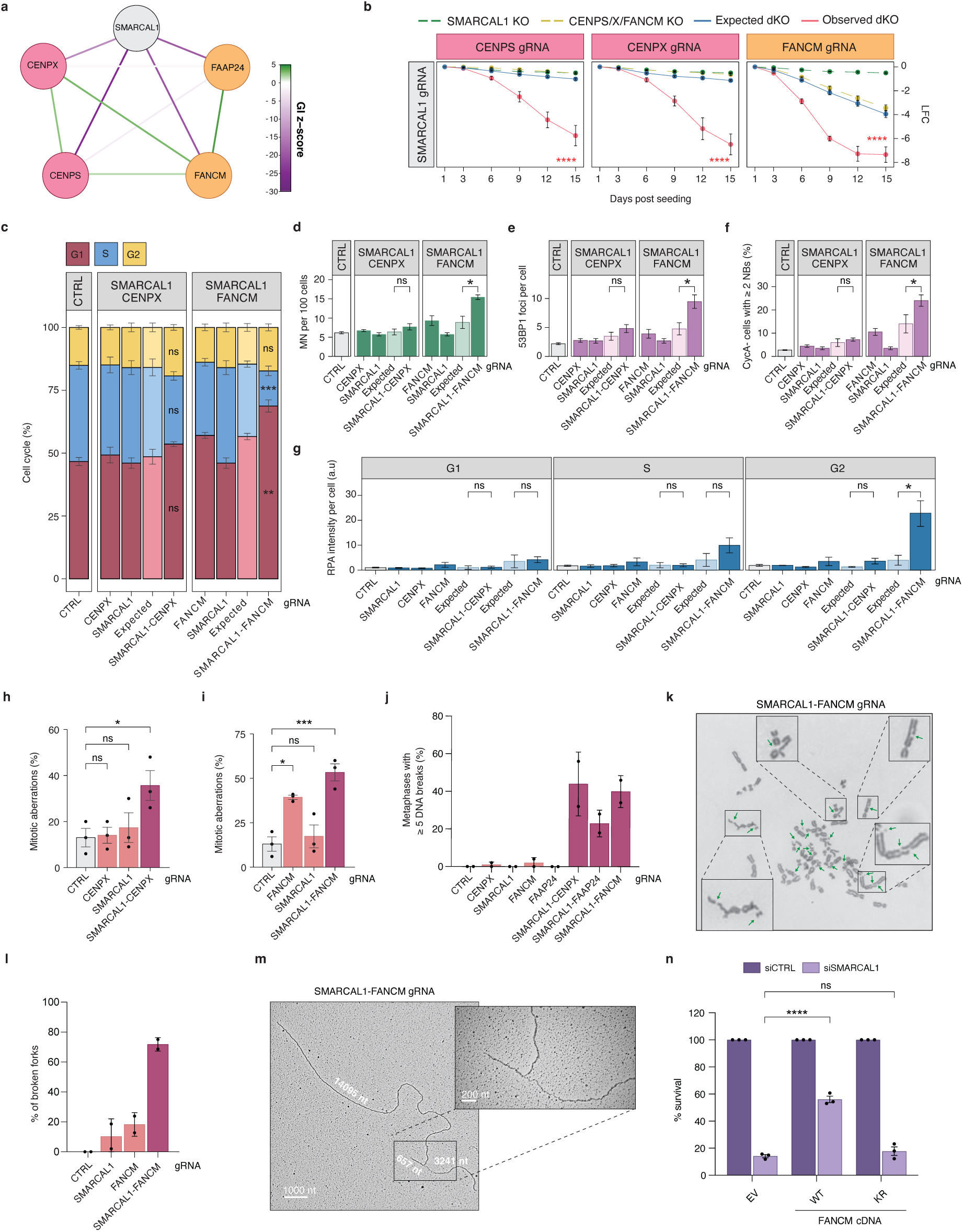
Negative interactions between *SMARCAL1* and genes encoding FANCM and its partners. **a**, GI network for *SMARCAL1*, *FANCM*, and *FANCM*-related partners from the RPE-1/MCF10A_integrated heatmap. GI z-scores < 5 and > −30 are shown. **b**-**n**, Cells were targeted with the indicated in4mer gRNAs. **b**, Validation of *SMARCAL1* interactions with *CENPS/X* and *FANCM* by two-color competition assay in MCF10A cells. Points represent the mean ± SEM (n = 3 biological replicates). Statistical significance was determined by two-way ANOVA comparing expected and observed values at T15. **c**-**g**, For each interaction, values are shown for individual single KOs, the dKO (observed), and the corresponding expected values for each dKO, calculated from single KO phenotypes. Expected values are plotted immediately to the left of their matched observed dKO values for each gene pair. Significant deviations between observed and expected values are indicated. Bars represent the mean ± SEM (n = 3 biological replicates), unless stated otherwise. Statistical significance was determined by unpaired t test. **c**, Cell cycle profiles for the indicated *SMARCAL1* interactions in MCF10A cells. Cell cycle phases were determined based on PCNA and DAPI staining. Significant deviations of observed from corresponding expected profiles are indicated for each cell cycle stage. Bars represent the mean ± SEM of six independent experiments (3 biological replicates and 2 independent imaging experiments per replicate). **d**, Quantification of micronuclei (MN) in MCF10A cells. **e**, Quantification of 53BP1 foci in cyclin A-positive (CycA+, S/G2) MCF10A cells. **f**, Quantification of 53BP1 nuclear bodies (NBs) in cyclin A-negative (CycA-, G1) MCF10A cells. **g**, Quantification of integrated RPA32 foci intensity (in arbitrary units, a.u.) in MCF10A cells across cell cycle phases (G1, S, and G2), which were distinguished by PCNA and DAPI staining. **h**, **i**, Quantification of spontaneous mitotic aberrations in MCF10A cells. Bars represent the mean percentage of anaphases with mitotic aberrations (± SEM) relative to all anaphases scored per sample (n = 3 biological replicates). At least 18 anaphase cells were scored per sample per experiment. Statistical significance was determined by one-way ANOVA. **j**, Quantification of DNA breaks detected by metaphase spreads in RPE-1 *TP53^-/-^*cells. Bars represent the mean percentage of metaphases with > 5 DNA breaks per metaphase (± SD) relative to the total number of metaphases examined per sample (n = 2 biological replicates). At least 50 metaphases were analyzed per sample per experiment. **k**, Representative image of a metaphase from *SMARCAL1-FANCM* dKO RPE-1 *TP53^-/-^* cells quantified in panel **j**. Chromatid and chromosome breaks are indicated by green arrows. **l**, Quantification of asymmetric (broken) replication forks detected by EM in MCF10A cells. Bars represent the mean percentage of broken forks (± SD) relative to the total number of replication intermediates (RIs) per sample (n = 2 biological replicates). At least 43 RIs were examined per sample per experiment. **m**, Representative image of a broken replication fork detected by EM in *SMARCAL1-FANCM* dKO MCF10A cells quantified in panel **l**. The length of the replication fork branches is indicated in nucleotides (nt). Scale bars, 1,000 and 200 nt. **n**, Quantification of survival assays of FaDu cells complemented with empty vector (EV), wild-type (WT) FANCM, or K117R (KR) ATPase-dead FANCM cDNA constructs and treated with either control (siCTRL) or SMARCAL1 (siSMARCAL1) siRNA. Bars represent the mean ± SEM (n = 3 biological replicates). The mean percentage of survival in siSMARCAL1-treated samples was normalized to the corresponding siCTRL samples. Statistical significance was determined by one-way ANOVA. **** = p-value ≤ 0.0001, *** = p-value ≤ 0.001, ** = p-value ≤ 0.01, * = p-value ≤ 0.05.

DSB mapping by END-seq^127^ confirmed the accumulation of DSBs in cells deficient for SMARCAL1 and FANCM, with breaks enriched at A/T-rich sequences (Extended Data Fig. 23a,b), as previously reported^68^. We also observed accumulation of DSBs at centromeric regions and Alu sequences (Extended Data Fig. 23a,b), suggesting that centromeric fragility, along with instability at other repetitive elements, may contribute to the lethality of cells lacking SMARCAL1 and FANCM. Notably, the palindromic sequence TAAAGCGCTTTA was enriched at or nearby END-seq peaks in both centromeric and non-centromeric regions (Extended Data Fig. 23c,d), suggesting that this palindrome could undergo preferential breakage in cells deficient for SMARCAL1 and FANCM. To investigate whether DNA breakage could occur during DNA replication, we analyzed replication fork intermediates by EM. This analysis revealed a >3-fold increase in broken forks in MCF10A cells deficient for SMARCAL1 and FANCM, implicating replication fork breakage as an underlying cause of the synthetic lethal interaction between *SMARCAL1* and *FANCM* (Fig. 6l,m and Extended Data Fig. 22e-g).

To assess the relevance of the SMARCAL1-FANCM interaction in a cancer context, we utilized FaDu cells, a head and neck squamous cell carcinoma cell line harboring biallelic loss-of-function mutations in *FANCM*^128^. FaDu cells displayed sensitivity to SMARCAL1 depletion, which was partially suppressed by expressing wild-type FANCM but not the catalytically inactive K117R^129^ (KR) mutant (Fig. 6n and Extended Data Fig. 22h,i). These findings indicate that the DNA translocase activity of FANCM is required for viability in SMARCAL1-deficient cells and demonstrate that this synthetic lethal interaction is conserved in a cancer-relevant setting.

### Loss of NHEJ genes suppresses the proliferation defect induced by APOLLO deficiency

We also identified positive interactions of the NHEJ genes *XRCC4*, *LIG4*, and *NHEJ1* with *DCLRE1B* in the RPE-1/MCF10A_integrated dataset (Fig. 2c and 7a). APOLLO (encoded by *DCLRE1B,* also known as SNM1B) is a 5′-3′ exonuclease involved in telomere maintenance^130^ and multiple aspects of the DNA damage response, including ICL repair, DSB processing, replication stress resolution, and DNA damage checkpoint signaling^131–134^. APOLLO is recruited by TRF2 to generate 3′ overhangs at leading-strand telomeres^135,136^, and its deletion causes perinatal lethality in mice, as well as leading-end telomeric fusions and proliferation defects in mouse embryonic fibroblasts (MEFs)—all of which are rescued by *Ku70* deletion^52,137^. This rescue was initially interpreted as evidence that APOLLO protects telomere ends from toxic joining by NHEJ. However, more recent work has shown that telomere fusions in Apollo-deficient MEFs are independent of Lig4 and instead require the MMEJ factors Lig3 and Pol θ, and that loss of DNA-PK (including Ku70) prevents telomere fusions by relieving suppression of Mre11-mediated end resection, thereby restoring 3′ overhang formation at telomeres^138^. In contrast, APOLLO depletion in non-transformed human cells induces telomere dysfunction and S-phase DNA damage signaling^136^ without leading to significant telomere fusions^136,139^, warranting further investigation of its interplay with NHEJ factors in human cells.

**Fig. 7.**
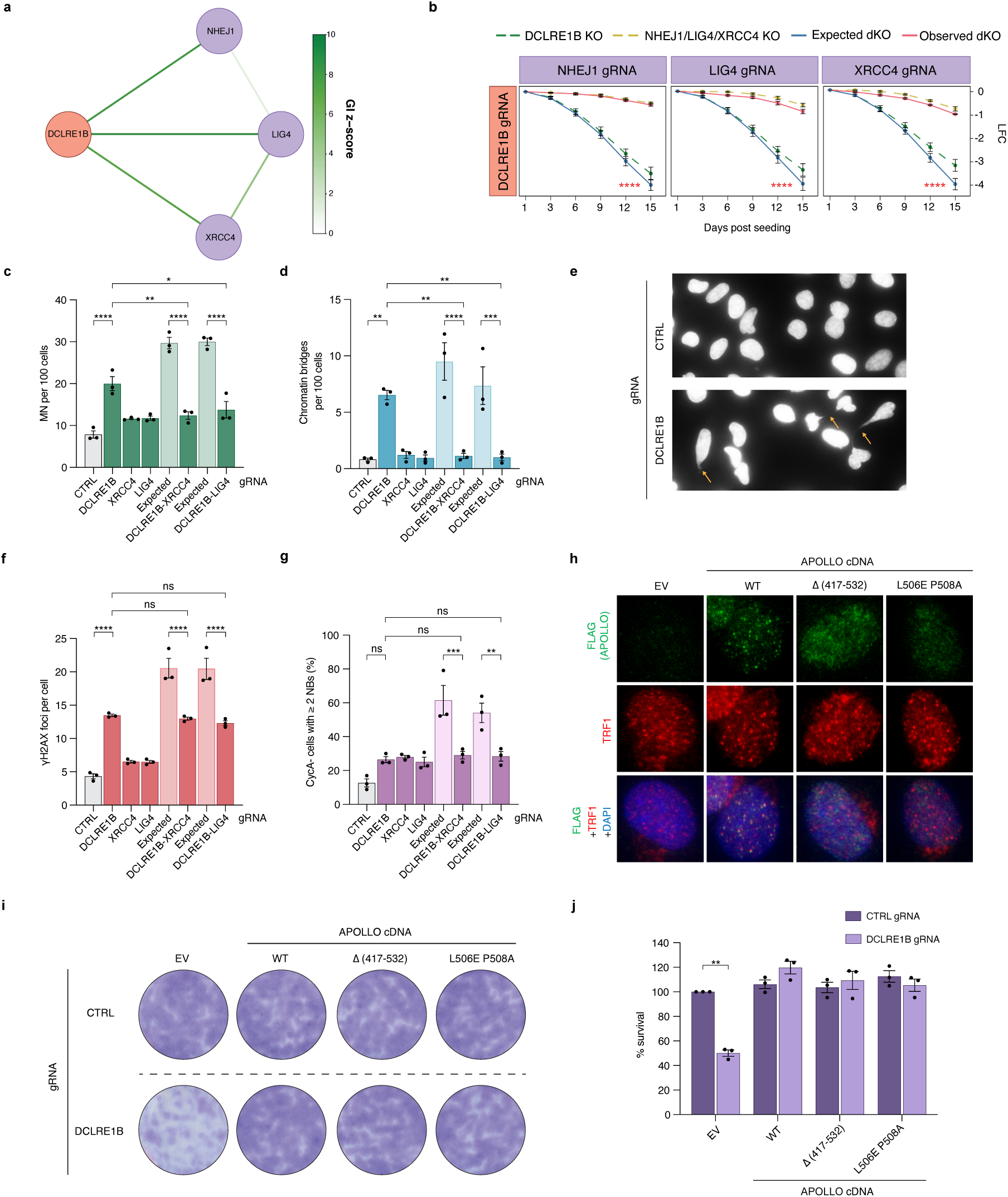
Suppressor interactions between *DCLRE1B* and the NHEJ genes *XRCC4*, *LIG4*, and *NHEJ1*. **a**, GI network for *DCLRE1B* and the NHEJ gene module (*XRCC4, LIG4, NHEJ1*) from the RPE-1/MCF10A_integrated heatmap. GI z-scores < 10 and > 0 are shown. Node colors are according to the module coloring scheme in Fig. 2a. **b**-**j**, Cells were targeted with the indicated in4mer gRNAs. **b**, Validation of *DCLRE1B* interactions with *XRCC4, LIG4,* and *NHEJ1* by two-color competition assay in RPE-1 *TP53^-/-^* cells. Points represent mean ± SEM (n = 3 biological replicates). Statistical significance was determined by two-way ANOVA comparing expected and observed values at T15. **c**, **d**, **f**, **g**, For each interaction, values are shown for individual single KOs, the dKO (observed), and the corresponding expected values for each dKO, calculated from single KO phenotypes. Expected values are plotted immediately to the left of their matched observed dKO values for each gene pair. Bars represent the mean ± SEM (n = 3 biological replicates). Statistical significance was determined by one-way ANOVA. **c**, Quantification of micronuclei (MN) in RPE-1 *TP53^-/-^* cells. **d**, **e**, Quantification and representative immunofluorescence images, respectively, of chromatin bridges in RPE-1 *TP53^-/-^* cells. **f**, **g**, Quantification of γH2AX foci and 53BP1 nuclear bodies (NBs) in cyclin A-negative (CycA-, G1) RPE-1 *TP53^-/-^*cells, respectively. **h**, Representative immunofluorescence images of FLAG-tagged APOLLO (green), TRF1 (red), or FLAG-tagged APOLLO together with TRF1 and DAPI (blue) staining of RPE-1 *TP53^-/-^*cells complemented with either empty vector (EV), wild-type (WT) APOLLO, Δ(417-532) APOLLO, or L506E P508A APOLLO cDNA constructs. **i**, **j**, Representative survival assay and quantification, respectively, of control (CTRL) and *DCLRE1B* KO MCF10A cells complemented with the indicated APOLLO cDNA constructs. Bars represent the mean percentage of survival (± SEM) relative to the CTRL sample complemented with EV construct (n = 3 biological replicates). Statistical significance was determined by unpaired t test. **** = p-value ≤ 0.0001, *** = p-value ≤ 0.001, ** = p-value ≤ 0.01, * = p-value ≤ 0.05.

To validate the genetic interactions between *DCLRE1B* and NHEJ factors, we conducted cell competition assays in RPE-1 cells with combined loss of APOLLO and XRCC4, LIG4, or NHEJ1 (Fig. 3d and 7b, Extended Data Fig. 17d, and Extended Data Table 1). While loss of APOLLO alone impaired cell fitness, this effect was rescued by disruption of *XRCC4*, *LIG4*, or *NHEJ1* (Fig. 7b). APOLLO loss also resulted in accumulation of micronuclei and chromatin bridges, phenotypes that were suppressed by disruption of *XRCC4* or *LIG4* (Fig. 7c-e and Extended Data Fig. 24a). APOLLO-deficient cells also showed accumulation of γH2AX foci, a subset of which colocalized with telomeres as telomere dysfunction-induced foci (TIFs), as well as a modest (but not statistically significant) increase in 53BP1 nuclear bodies and foci (Fig. 7f,g and Extended Data Fig. 24b-f). While the γH2AX foci and 53BP1 nuclear bodies were not rescued by disruption of *XRCC4* or *LIG4*, the corresponding phenotypes in double mutants were less pronounced than expected (Fig. 7f,g and Extended Data Fig. 24b-d). These studies suggest that the suppression of mitotic aberrations may account for the rescue of the growth defect of APOLLO-deficient cells upon loss of NHEJ factors.

To investigate this observation in greater mechanistic detail, we complemented APOLLO-deficient RPE-1 cells with wild-type APOLLO or TRF2-binding-deficient mutants carrying either a C-terminal deletion (Δ 417-532) or point mutations (L506E, P508A)^51,140^ (Extended Data Fig. 24g). As expected, the APOLLO mutants displayed defective interaction with TRF2 and impaired localization to telomeres (Fig. 7h and Extended Data Fig. 24h,i). Notably, both wild-type and TRF2-binding-deficient APOLLO mutants rescued the cell fitness defect of APOLLO-deficient cells with comparable efficiency (Figure 7i,j), indicating that this defect is independent of the APOLLO-TRF2 interaction and suggesting a potential telomere-independent function for APOLLO in this context.

## Discussion

In this study, we generated a complete pairwise GI map of 233 DDR genes by conducting combinatorial genetic interaction screens in two cell lines, MCF10A and RPE-1. From these screens, we identified over 750 high-confidence genetic interactions (FDR < 0.1) (Supplementary Table 4) and validated 59 interactions (36 negative, 23 positive) (Fig. 3 and Supplementary Table 14). Additional characterization of 4 genetic interaction sets further defined DDR pathway interactions between TLS and the MCM8/9-HROB complex, mitotic repair genes and the FA pathway, the DNA translocases *SMARCAL1* and *FANCM* (or its associated factors), and NHEJ genes and *DCLRE1B* (Fig. 4-7). Collectively, our DDR GI map highlights functional relationships among specialized DDR pathways, providing a valuable resource for future studies of the human DDR.

### The Cas12a in4mer system enables robust genetic interaction screens in human cells at scale

In our screening approach we utilized the Cas12a-based in4mer system to mediate efficient double gene disruption^16^. Previous combinatorial screening approaches have utilized CRISPRi to avoid the issue of low dKO efficiency in diploid cells^7,67,141–145^. Although CRISPRi produces homogeneous knockdown of all alleles, it has been shown to identify different categories of essential genes when compared to KO-based CRISPR screens, potentially due to phenotypes that can be masked by low-level residual expression of the target genes^146^. For this reason, our CRISPR KO-based interaction map may uncover a different set of genetic interactions that are missed in CRISPRi-based screens. While CRISPR-KO screens are generally considered less suited for essential gene interactions, inclusion of an early time point (T9) in our screens enabled their detection before fitness defects obscured pairwise interaction measurements (Extended Data Fig. 14).

Direct comparison of genetic interaction screening approaches is challenging due to the low likelihood of overlapping hits between unrelated screens, arising from the sparse landscape of genetic interactions and the vast combinatorial space of potential gene pairs. In our work, we employed 13 gold standard synthetic lethal interactions between paralogs as positive controls, providing strong evidence for the efficacy of Cas12a-based in4mer screens (Extended Data Fig. 2b and Supplementary Table 2)^16^. Additionally, strong agreement between our screen results and data obtained in isogenic CRISPR-KO screens further highlights the utility of redundant Cas12a targeting for combinatorial screening (Extended Data Fig. 2e).

Analysis of our genetic interaction screens revealed more than 750 high-confidence (FDR < 0.1) genetic interactions between individual DDR genes (Supplementary Table 4). Network-level analysis uncovered genetic interaction modules composed of gene sets sharing multiple interacting partners (Fig. 2a,b, Extended Data Fig. 6-8, 12a,b, 13a,b, and Supplementary Table 7). Many of these modules comprised functionally related genes, providing increased confidence in pathway-level interactions (Extended Data Fig. 11). Notably, correlation of GI profiles from our combinatorial screens achieved similar performance to DepMap co-essentiality data in identifying complex members (Extended Data Fig. 9a,b), and approximately 30% of modules were significantly enriched in genes encoding proteins predicted to form physical complexes by AlphaFold2 (Extended Data Fig. 9c-e). Validation assays confirmed the phenotype of ∼75% of significant individual hits and candidates nominated through expanded pathway analysis (Fig. 3b-d). Together with the recovery of multiple known interactions (Fig. 1e), this high validation rate establishes our GI maps as a robust resource for the scientific community.

### Combinatorial Cas12a-based screens identify synthetic lethal interactions within the DDR

In-depth mechanistic interrogation of 3 synthetic lethal interaction sets provided insights into the functional basis for a subset of the top genetic interaction patterns observed in our screen. We showed that cells are dependent on the TLS REV1-Pol ζ complex following loss of the MCM8/9-HROB complex (Fig. 4). This functional dependency was identified as shared synthetic lethality between two genetic interactions modules comprised of *MCM8/MCM9/HROB* and *REV1*/*MAD2L2/REV3L* (Fig. 4a). The observed increase in RPA levels and γH2AX foci in G2-phase cells upon loss of both MCM8/9 and REV7 (Fig. 4d), along with the sensitivity of MCM8/9-deficient cells to the TLS inhibitor JH-RE-06 (Fig. 4i), suggest that loss of REV1-Pol ζ-mediated TLS underlies the observed lethality induced by REV7 deficiency in MCM8/9-deficient cells. Consistent with this model, a REV7 mutant (W171A) defective in REV3L binding failed to suppress the lethality of MCM9- and REV7-deficient cells (Fig. 4g,h). We and others have previously shown that MCM8/9-HROB complex functions as a DNA helicase involved in homology-directed repair and replication fork progression under replication stress^41,85^. We also demonstrated that REV1-Pol ζ-mediated TLS is required for the filling of ssDNA gaps induced by the DNA primase-polymerase PRIMPOL in HR-deficient *BRCA1*-mutant cells^93^. However, in contrast to *BRCA1*-mutant cells with REV1-Pol ζ inactivation, loss of REV1 or REV7 in MCM9-deficient cells did not induce large (>1,000 nt) ssDNA gaps at replication forks (Extended Data Fig. 19f), which are dependent on PRIMPOL-mediated repriming and nucleolytic processing by MRE11, EXO1, and DNA2^93,147^. Moreover, unlike the synthetic lethality between *BRCA1* and *REV1*, the synthetic lethality between *MCM9* and *REV1* or *MAD2L2* was not suppressed by PRIMPOL loss (Extended Data Fig. 19b-e).

These findings indicate that the functional interaction between MCM8/9 and REV1-Pol ζ is PRIMPOL-independent and suggest that, unlike BRCA1/2, MCM8/9 may not protect ssDNA gaps from extensive nucleolytic processing. Instead, combined loss of MCM9 and REV1 or REV7 led to a 2-to-3-fold increase in collapsed replication fork intermediates (Fig. 4j,k and Extended Data Fig. 19g-j), implicating fork collapse as the underlying cause of synthetic lethality. MCM8/9 associates with nascent DNA during replication and promotes fork progression upon replication stress^86,148^, while REV1-Pol ζ has been implicated in the bypass of DNA lesions at stalled forks^149^. Thus, MCM8/9 and REV1-Pol ζ may represent alternative pathways for promoting fork progression at DNA lesions or difficult-to-replicate sites. In the absence of both pathways, persistent fork stalling may lead to nucleolytic cleavage. Notably, loss of MCM8/9 has been shown to increase MUS81-dependent cleavage of stalled forks^86^, suggesting that MUS81 or other SLX4-associated nucleases may mediate fork collapse in cells lacking both MCM8/9 and REV1-Pol ζ. These findings reveal a novel functional interplay between MCM8/9 and REV1-Pol ζ required for preserving replication fork integrity and maintaining cell viability.

Synthetic lethal interactions between a 10-gene FA module and a module composed of three mitotic DNA repair genes, *CIP2A*, *GEN1,* and *RHNO1*, revealed a novel dependency of FA-deficient cells (Fig. 5). These findings suggest that GEN1, CIP2A, and RHINO may function in a mitotic DNA repair pathway that resolves late replication intermediates at difficult-to-replicate regions, structures that FA proteins are known to bind and stabilize^150^. In our model, FA deficiency leads to incomplete replication at difficult-to-replicate loci, producing under-replicated DNA structures that persist into mitosis. Previous studies have shown that the CIP2A-TOPBP1 complex recognizes these structures and recruits SLX4-associated nucleases to promote their resolution^151–153^. However, because SLX4 (FANCP) is thought to function downstream of the FA core complex and FANCD2-FANCI^154–156^, SLX4-mediated resolution may be compromised in FA-deficient cells, causing these intermediates to accumulate. Consistent with this model, FANCL-deficient cells display increased CIP2A foci in mitosis (Extended Data Fig. 20p,q), indicative of persistent unresolved DNA structures. In the absence of functional SLX4-mediated resolution, FA-deficient cells become dependent on GEN1 to cleave under-replicated DNA intermediates, which are known GEN1 substrates^106^. Supporting this model, complementation assays demonstrated that GEN1’s nuclease activity is required for viability in FANCL-deficient cells (Fig. 5h,i). In contrast, BRCA1-deficient cells show no dependency on GEN1 (Extended Data Fig. 20b-f), in line with the observation that SLX4-associated nucleases operate with CIP2A-TOPBP1 to resolve under-replicated DNA intermediates in BRCA1-deficient cells^151,152^. GEN1-dependent cleavage of these mitotic intermediates generates DSBs that require repair to maintain chromosome stability. We propose that RHINO promotes the repair of these GEN1-generated DSBs through MMEJ, likely via its association with Pol θ, which is also required for the survival of FA-deficient cells (Extended Data Fig. 21a,b). Consistent with this model, GEN1 is required for the accumulation of RHINO and Pol θ foci in mitosis (Fig. 5j,k and Extended Data Fig. 20n,o). DSBs that escape mitotic repair could persist into the subsequent G1 phase, explaining the G1 accumulation, micronuclei formation, and 53BP1 nuclear bodies observed in cells deficient for FA genes combined with GEN1, CIP2A, or RHINO loss (Fig. 5c). Collectively, these findings uncover a mitotic DNA repair pathway dependent on GEN1 nuclease activity, CIP2A, RHINO, and Pol θ that becomes essential in FA-deficient cells.

Additionally, we characterized synthetic lethal interactions between *SMARCAL1* and genes encoding FANCM and its associated partners FAAP24 and CENPS/X (Fig. 6), which have also been recently reported by others^67,68^. Both SMARCAL1 and FANCM promote the remodeling of branched DNA structures, with FANCM functioning in complex with CENPS/X and FAAP24^118,119,123,125,126^. The synthetic lethality between these factors suggests that they play compensatory roles in preserving genomic stability at endogenous sites of replication stress. Indeed, combined disruption of *SMARCAL1* and *FANCM* resulted in pronounced DNA damage, mitotic aberrations, DNA breaks, and chromosome fragmentation (Fig. 6c-k and Extended Data Fig. 22a-d). Our observation of a ∼3-fold increase in collapsed forks upon combined SMARCAL1 and FANCM loss (Fig. 6l,m) indicates that DSBs form during DNA replication and implicate replication fork breakage as an underlying cause of the synthetic lethality. Accumulation of DSBs at A/T-rich sequences and Alu elements in SMARCAL1- and FANCM-deficient cells (Extended Data Fig. 23a,b) indicates that SMARCAL1 and FANCM are critical for resolving replication impediments caused by repetitive sequences. Notably, DSB sites in SMARCAL1- and FANCM-deficient cells are enriched for the palindromic sequence TAAAGCGCTTTA, consistent with the ability of SMARCAL1 and FANCM to resolve cruciform structures^67^ (Extended Data Fig. 23c,d). DSBs also accumulate at centromeric regions (Extended Data Fig. 23a,b), suggesting that SMARCAL1 and FANCM contribute to centromere stability. Consistent with this notion, CENPS/X are centromeric proteins that aid kinetochore assembly and promote FANCM recruitment to centromeres^157,158^. Furthermore, the *S. pombe* orthologs of CENPS/X (Mhf1/2) and FANCM (Fml1) suppress crossover events at centromeric repeats and prevent gross chromosomal rearrangements^159^. Whether SMARCAL1 compensates for FANCM loss in maintaining centromere stability warrants further investigation. Beyond centromeres, FANCM and SMARCAL1 have been shown to suppress replication stress at telomeres in ALT-positive cells^160–165^, and their combined loss enhances telomeric DNA damage in this context^165^. Whether combined FANCM and SMARCAL1 deficiency leads to unresolved replication intermediates at telomeres that undergo aberrant processing in non-ALT cells remains to be determined. Importantly, other SMARCAL1-related fork remodelers, such as ZRANB3 and HLTF, are not essential in cells deficient for FANCM, CENPS/X, or FAAP24 (Supplementary Table 3), indicating a specific and non-redundant requirement for SMARCAL1 in the absence of FANCM^67,68^. Together, these findings underscore the critical roles of SMARCAL1 and FANCM in maintaining genome stability at repetitive sequences prone to replication stress.

Among other notable synthetic lethal interactions, we identified relationships that follow several organizing principles. First, we observed synthetic lethality between factors whose loss generates specific DNA intermediates and the pathways required to repair them. For example, *RNASEH1*, whose loss causes R-loop accumulation, triggering replication-transcription conflicts and fork collapse^166^, is synthetically lethal with NHEJ factors (*LIG4*, *XRCC4*, *NHEJ1*, *PRKDC*, *ERCC6L2*, *CYREN*), which are required for DSB repair (Extended Data Fig. 25a). Second, we identified synthetic lethality between genes whose disruption causes DNA damage, such as *POLE3*/*POLE4* (whose loss reduces leading-strand processivity and increases replication stress^74^) or *BOD1L1* (whose loss exposes stalled replication forks to excessive resection^77^), and genes required for DNA damage sensing through ATR activation (*RAD17*/*HUS1*/*RAD9A*)^78^ (Extended Data Fig. 14f). Third, we observed synthetic lethality between parallel pathways with overlapping functions: *ERCC6L2* and shieldin (*SHLD1/SHLD2/SHLD3*), which independently restrict end resection to promote NHEJ^167,168^ (Extended Data Fig. 25b); POLK and REV1-Pol ζ (*REV1*/*MAD2L2*/*REV3L*), which promote TLS across DNA lesions^169^ (Extended Data Fig. 25c); and factors that activate ATR through separate mechanisms^75,76^ (*RAD17/HUS1*/*RAD9A* and *ETAA1*) (Extended Data Fig. 14f). Together, these interactions illuminate the modular architecture of the DDR, where loss of one pathway creates dependencies on parallel or downstream repair mechanisms.

### Cas12a screens identify positive DDR interactions

Our screens also identified a suppressor interaction between *DCLRE1B*, a gene encoding the DNA exonuclease APOLLO, and the NHEJ genes *XRCC4*, *LIG4*, and *NHEJ1*. APOLLO-deficient cells displayed increased micronuclei and chromatin bridges, phenotypes suppressed by *XRCC4* or *LIG4* disruption (Fig. 7c-e and Extended Data Fig. 24a). APOLLO is involved in DNA repair and the maintenance of telomeres, where it is recruited by TRF2 through its C-terminal domain^131–136^. Notably, complementation studies revealed that TRF2-binding mutants rescued the growth defect of APOLLO-deficient cells comparably to wild-type APOLLO (Fig. 7i,j), suggesting that this phenotype is independent of TRF2-mediated telomeric recruitment. Consistent with this observation, APOLLO acts redundantly with RAP1 to suppress NHEJ at telomeres in human cells, with NHEJ-dependent telomeric fusions only observed upon combined loss of APOLLO and RAP1^139^. Thus, in cells lacking APOLLO alone, NHEJ-dependent toxicity might arise at non-telomeric sites rather than at telomeres. These substrates could include DNA repair- or replication-associated intermediates, where aberrant NHEJ-mediated processing in the absence of APOLLO may generate toxic DNA structures. Together, these findings uncover a suppressor interaction between APOLLO and NHEJ factors and suggest that APOLLO may promote cellular viability through non-telomeric functions.

Other notable suppressor interactions identified in our screens reflect pathway antagonism. These include relationships between genes encoding the end resection suppressor ERCC6L2 and the pro-resection ATM kinase, an interaction recently described^50,170,171^; and the anti-recombinase BLM and the pro-recombinase Shu complex (ZSWIM7/SPIDR/SWSAP1) (Extended Data Fig. 25d), a relationship previously characterized in yeast and mice^48,69^. We also identified a suppressor interaction between genes encoding the DNA endonuclease ERCC1/ERCC4 and BABAM2, a subunit shared by the BRCA1-A and BRISC complexes^172^. However, the pattern of interactions suggests that this effect is more likely mediated through the BRCA1-A complex that inhibit DNA resection and HR^173^, because *ERCC1/ERCC4* also show mild positive interactions with other BRCA1-A components, including *UIMC1*, *BABAM1*, and *ABRAXAS1*, but not with *ABRAXAS2*, which is specific to BRISC (Extended Data Fig. 25e). This finding suggests that enhanced HR efficiency upon BRCA1-A complex disruption may compensate for defects in processing stalled forks or ICL intermediates caused by loss of ERCC1/ERCC4^155,174^. Importantly, we validated the majority of these interactions using two-color competition assays, demonstrating that our screening approach reliably identifies suppressor interactions (Fig. 3c,d and Extended Data Fig. 17e). Together, these observations reveal antagonistic relationships and suppressor interactions that shape the architecture of the DDR.

### Cas12a screens identify genetic interactions of therapeutic relevance

Given their function in maintaining genomic stability, many of the DDR genes in our library are known cancer drivers that could potentially be targeted therapeutically through their synthetic lethal interactors (Extended Data Fig. 4). The dependency of cells deficient in FA genes, often disrupted in various tumor types, including breast, ovarian, bladder and head and neck cancer^107^, on GEN1’s nuclease activity illustrates the utility of genetic interaction data for identifying clinically relevant targets (Fig. 5h,i). The reciprocal synthetic lethality between *SMARCAL1* and *FANCM* suggests therapeutic opportunities across multiple contexts: *FANCM* loss-of-function mutations occur in breast, ovarian, and head and neck tumors^128,175,176^, while SMARCAL1 deficiency has been observed in a subset of osteosarcomas and gliomas characterized by alternative lengthening of telomeres^177–180^. Sensitivity of MCM8/MCM9-deficient cells to the TLS inhibitor JH-RE-06 is similarly relevant, as *MCM8/9* mutations are associated with Lynch-like syndrome and breast cancer^181,182^. Notably, we validated the sensitivity of FANCA- and FANCM-deficient head and neck cancer cells to loss of GEN1 and SMARCAL1 (Fig. 5l and 6n), respectively, highlighting these factors as potential therapeutic targets for cancer treatment.

Additional interactions suggest further therapeutic opportunities. *ATM*, among the most frequently mutated DDR genes (∼2–10% across cancer types including hematological malignancies, lung, gastric, prostate, and pancreatic cancers^183^), exhibits synthetically lethality with *RNASEH2A/RNASEH2B* (Fig. 2c, 3c, and Extended Data Fig. 4c), suggesting that RNASEH2 could be targeted for treating ATM-deficient tumors. Notably, combined Atm and Rnaseh2 inactivation in mice causes severe genome instability through aberrant NHEJ^184^. Together, these examples demonstrate how our genetic interaction map can guide identification of targetable vulnerabilities in DDR-deficient cancers.

In summary, our high-quality map of genetic interactions among 233 DDR-associated genes, obtained through Cas12a-based combinatorial screens, enables the identification of functional relationships among DDR genes and pathways, while pinpointing potential targets for the treatment of cancer and other diseases associated to DDR gene mutations. Applying similar strategies to other cellular pathways would provide additional insights into the inner workings of cellular networks and lead to novel approaches for targeted therapies of cancer and other diseases.

## Methods

### Cell lines and cell culture

Human mammary epithelial MCF10A cells (ATCC, CRL-10317) were cultured in a 1:1 mixture of Dulbecco’s Modified Eagle’s Medium and Nutrient Mixture F-12 (Ham) containing GlutaMAX (DMEM/F-12, Gibco/Thermo Fisher Scientific), supplemented with 5% horse serum (Gibco/ Thermo Fisher Scientific), 10 mg/ml insulin (Sigma-Aldrich), 0.5 mg/ml hydrocortisone (Sigma-Aldrich), and 25 ng/ml human epidermal growth factor (PeproTech). Human epidermal growth factor was added to media just before use and media for MCF10A cells was replaced every 3 days. Immortalized human embryonic kidney HEK293T cells (ATCC, CRL-11268) were cultured in DMEM containing 2 mM L-glutamine (Gibco/Thermo Fisher Scientific), supplemented with 10% Fetalgro (Rmbio). hTERT-immortalized human retinal pigment epithelial RPE-1 *TP53^-/-^* cells^185^ were cultured in DMEM/F-12 containing GlutaMAX (Gibco/Thermo Fisher Scientific), supplemented with 10% fetal bovine serum (FBS, Corning). Human osteosarcoma U2OS cells expressing Halo-Pol θ and RHINO-mNeonGreen were cultured in RPMI medium containing 2 mM L-glutamine (Gibco), supplemented with 10% FBS (Corning). The generation of U2OS Halo-Pol θ RHINO-mNeonGreen cells is described below. Head and neck squamous cell carcinoma (HNSCC) cell line OHSU-SCC-974 [Fanconi Anemia Research Materials (FARM) repository], derived from a biopsy of a primary lesion from a *FANCA*^-/-^ Fanconi Anemia patient, was cultured in Eagle’s Minimum Essential media containing 2 mM L-glutamine (MEM, Corning), supplemented with 10% FBS. Squamous cell carcinoma cell line FaDu (ATCC, HTB-43) was cultured in DMEM containing 2 mM L-glutamine (Gibco/Thermo Fisher Scientific), supplemented with 10% FBS. All growth media was supplemented with 100 U/ml penicillin and 100 mg/ml streptomycin (Pen/Strep, Thermo Fisher Scientific) and all cells were grown in a humidified atmosphere at 37° C and 5% CO_2_.

### Plasmids

For mid-throughput two-color competition validation experiments we generated pHAGE-mCherry-NLS or pHAGE-GFP-NLS lentiviral vectors using Gateway recombination with LR Clonase II (Thermo Fisher) (Supplementary Table 15-5). In brief, mCherry-NLS or GFP-NLS were amplified from LentiGuide-NLS-GFP (Addgene #185473) or LentiGuide-NLS-mCherry (Addgene #185474) using the primers SAM470 and SAM471 (Supplementary Table 15-2) and integrated into the pDONR223 donor vector. Donor vectors were then recombined into the pHAGE-Ct-FLAG-HA-HYGRO-DEST and pHAGE-Ct-FLAG-HA-NEO-DEST vectors by Gateway cloning, as previously described^85^. pHAGE-mCherry-NLS-HYGRO (pSAM148) and pHAGE-GFP-NLS-HYGRO (pSAM149) were used for validation experiments in MCF10A cells, while pHAGE-mCherry-NLS-NEO (pSAM558) and pHAGE-GFP-NLS-DEST (pSAM557) were employed for two-color competition experiments in RPE-1 *TP53^-/-^* cells.

For two-color competition experiments in MCF10A cells, lentiviral plasmids containing in4mer gRNA arrays of interest were cloned using Golden Gate assembly (pSAM387-482; Supplementary Table 15-1). Due to the length (168 bp) and the number (90) of gRNA arrays needed for validation tests, we ordered validation gRNAs as an oligo pool (Twist Bioscience). gRNA oligos were designed as follows: 5’-[forward common primer binding]-[forward unique primer binding]-CGTCTCgagat-[4-gRNA array]-ttttttgaatgGAGACG-[reverse unique primer binding]-[reverse common primer binding]-3’, with BsmBI/Esp3I recognition sites in capital letters. The complete gRNA oligo pool was amplified using primers SAM844 and SAM845 (Supplementary Table 15-2), followed by amplicon purification with PCR DNA Fragments Extraction Kit (IBI Scientific) according to manufacturer’s instructions. Then, specific oligos were amplified from the oligo pool using primers designed for unique primer sites included in each oligo (mid-throughput PCR primers, Supplementary Table 15-2) and purified with PCR DNA Fragments Extraction Kit (IBI Scientific) according to manufacturer’s instructions. Individually amplified oligos (Supplementary Table 15-1) were cloned into the lentiviral enCas12a gRNA expression backbone pRDA_052 (Addgene #136474; pSAM112) (Supplementary Table 15-5) using the Golden Gate assembly protocol with Esp3I (New England Biolabs) and T4 ligase (New England Biolabs) as follows: 37 °C 30 min; 30x cycles of [37 °C 5 min, 16 °C 5 min]; 37 °C 1 h; 65 °C 20 min.

For two-color competition experiments in RPE-1 *TP53^-/-^* cells and follow-up validation experiments, in4mer gRNA vectors (pSAM486-673) were generated (Supplementary Table 15-1). These gRNA sequences were ordered as individual eBlocks (IDT) and cloned directly into pRDA_052 using the same Golden Gate assembly approach. eBlocks contained 66 bp of random DNA sequence 5’ and 3’ of the Esp3I sites to accommodate the minimum required size by IDT. eBlock sequences were designed as follows: 5’-[66 bp arbitrary DNA sequence]-CGTCTCgagat-[4-gRNA array]-ttttttgaatgGAGACG-[66 bp arbitrary DNA sequence]-3’, with BsmBI/Esp3I recognition sites in capital letters (Supplementary Table 15-1).

Additional Cas12a 2-gRNA vectors targeting *OR1E1-OR52A5* (CTRL, pSAM134) and *FANCL-GEN1* (pSAM321) were cloned using two oligos joined together by overlap extension. These constructs contained one gRNA for each gene in the array allowing for the annealing and extension of two oligos. SAM454 and SAM459 were annealed (Supplementary Table 15-3) to generate pSAM134 (Supplementary Table 15-1), and SAM659 and SAM724 were annealed (Supplementary Table 15-3) to generate pSAM321 (Supplementary Table 15-1). After annealing and overlap extension, products were amplified with SAM463 and SAM464 (Supplementary Table 15-2) to make a 103 bp dsDNA fragment containing both gRNAs (Supplementary Table 15-1). These dsDNA fragments were then inserted into pRDA_052 via the same Golden Gate assembly protocol. A similar cloning approach was used for the mid-throughput validation control (CTRL) guide construct targeting *OR1E1* and *OR52A5* genes (pSAM483, Supplementary Table 15-1). Here SAM875 and SAM876 (Supplementary Table 15-3) were annealed and extended before amplification with SAM877 and SAM878 (Supplementary Table 15-2), followed by insertion into pRDA_052.

To generate the screening backbone, pSAM112-UMI (Supplementary Table 15-5), a 6 bp degenerate sequence was cloned downstream of the 3’ BsmBI site of the pRDA_052 plasmid. In brief, two oligos (UMI_forward_anneal/UMI_reverse_anneal; Supplementary Table 15-2) were annealed and ssDNA overhangs were filled via end extension. This dsDNA product was then digested with BbsI and ligated into the BsmBI linearized pRDA_052 backbone.

For GEN1 complementation assays, we generated a catalytic dead GEN1 mutant (CD GEN1, pSAM512) and a wild-type GEN1 (WT GEN1, pSAM516) expression constructs (Supplementary Table 15-5). To generate these constructs, WT GEN1 cDNA was amplified from MCF10A cells using SAM751 and SAM752 (Supplementary Table 15-2). This cDNA was integrated into the pDONR223 donor vector. The catalytic inactivating mutations were then installed by PCR using the SAM818 and SAM819 primer pair (Supplementary Table 15-2). The WT and CD cDNA donors were then recombined into the pHAGE-Ct-FLAG-HA-HYGRO-DEST vector by Gateway cloning. In these experiments, we used a single gRNA targeting the intron-exon 4 junction of GEN1 to knockout (KO) the endogenous *GEN1* gene without affecting GEN1 cDNA cloned in pHAGE destination plasmids.

For REV7 complementation assays, we generated gRNA-resistant REV7 mutants that either abolish the REV7-SHLD2 interaction (Y63A REV7, pSAM696) or REV7-REV3L interaction (W171A REV7, pSAM697), as well as wild-type REV7 (WT REV7, pSAM695) expression constructs (Supplementary Table 15-5). REV7 cDNA was amplified from MDA-MB-436 using primers J744 and J745 (Supplementary Table 15-2) and cloned into pDONR223. Stop codon (J1618-J1716) and gRNA resistant mutations were added in subsequent rounds of mutagenesis by inverse PCR with oligo pairs J1717-J1718 and J1719-1720 (Supplementary Table 15-2). The Y63A and W171A separation-of-function alleles were also generated by inverse PCR using primer pairs SAM1073-SAM1074 and SAM1077-SAM1078, respectively (Supplementary Table 15-2). Guide-resistant versions of REV7 WT and separation-of-function alleles from pDONR223 were recombined into the pHAGE-Nt-FLAG-HA-NEO vector with LR reactions.

For FANCM complementation assays, wild-type FLAG-FANCM (WT FANCM, pSAM701) and ATPase-dead FLAG-FANCM mutant K117R (KR FANCM, pSAM702) were amplified from pFastBac backbones (a gift from Andrew Deans) using SAM1021-SAM1022 and cloned into pDONR223 by BP reaction (Supplementary Table 15-2 and 15-5). FLAG-tagged WT FANCM and KR FANCM cDNA donors were recombined into the pHAGE-NEO-DEST vector by Gateway cloning.

For APOLLO complementation assays, we generated gRNA-resistant wild-type APOLLO (WT APOLLO, pSAM686) and two TRF2-interaction deficient mutants (Δ(417-532) APOLLO, pSAM688; L506E P508A APOLLO, pSAM689)^51,140^ (Supplementary Table 15-5). Briefly, pDONR223-APOLLO from the ORFeome collection^186^ was mutated in a single Golden Gate reaction to add simultaneously a stop codon and two gRNA resistant mutations. A custom DNA string including the desired modifications was ordered from Twist Biosciences and used to amplify by PCR the fragments before Golden Gate reactions. Detailed information for Golden Gate reactions will be provided upon request. Δ(417-532) APOLLO and L506E P508A APOLLO were generated by inverse PCR using primer pairs OTL103-OTL104 and OTL105-OTL106, respectively (Supplementary Table 15-2).

To tag *RHNO1* with mNeonGreen, RHNO1 gRNA was cloned into BpiI-digested px330-U6-Chimeric_BB-CBh-hSpCas9 backbone (Addgene #42230)^187^ using standard procedures with ssDNA oligos purchased from IDT (Supplementary Table 15-3). The homology-directed repair (HDR) donor plasmid for RHINO-mNeonGreen was cloned by Gibson Assembly into pFastBac Dual backbone (Thermo Fisher Scientific #10712024) after linearization with HpaI. Gibson Assembly consisted of three inserts including a left homology arm, right homology arm and an intervening sequence containing the mNeonGreen sequence. Homology arms consisted of 449 bp homologous to the genomic DNA directly upstream and downstream of the Cas9 cleavage site and 25 bp overlaps on both ends for Gibson Assembly cloning. Homology arms were ordered as double-stranded gene fragments from IDT (Supplementary Table 15-3). The mNeonGreen fragment was PCR amplified from a plasmid encoding mNeonGreen-RPA32^102^ (Supplementary Table 15-3). All plasmids were validated by Sanger Sequencing. These plasmids will be made available from Addgene.

### Cell line generation

To generate MCF10A, RPE-1 *TP53^-/-^*, OHSU-SCC-974, and U2OS Halo-Pol θ RHINO-mNeonGreen cells constitutively expressing enCas12a, these cells were transduced with lentivirus carrying pRDA_174 (Addgene #136476; pSAM113)^22^ (Supplementary Table 15-5). 24 h later, cells were treated with blasticidin at 10 μg/ml for 72 h to select transduced cells. Then, pooled enCas12a cell lines were maintained in the media containing 2.5 μg/ml blasticidin. For MCF10A and RPE-1 *TP53^-/-^* cell lines, high-editing enCas12a clones were recovered. MCF10A cells were seeded at low density in 10 cm dishes, while RPE-1 cells were sorted as single cells into 96-well plates by flow cytometry (BD Influx Cell Sorter), and outgrown until cell populations were recovered. Single clones were screened for the level of enCas12a expression by western blotting using anti-HA antibody and clonal editing efficiency was subsequently assessed as described in the “Analysis of enCas12a gene editing” section using a Cas12a 2-gRNA construct targeting the olfactory receptor gene *OR1E1* (pSAM134, Supplementary Table 15-1). Single clones exhibiting the highest editing efficiencies were selected for further experiments.

To generate pooled KO cell lines, pooled enCas12a cell lines or high-efficiency clonal enCas12a cell lines were seeded in 12-well plates and, the next day, were transduced with the appropriate lentiviral in4mer gRNA constructs. 24 h after transduction, puromycin was added to select for transduced cells (8 μg/ml for 24 h for MCF10A cells, 20 μg/ml for 72 h for RPE-1 *TP53^-/-^* cells, 8 μg/ml for 48 h for OHSU-SCC-974 cells, 2.5 μg/ml for 24 h for U2OS Halo-Pol θ RHINO-mNeonGreen cells). Then, pooled KO cell lines were maintained in the media containing 2.5 μg/ml blasticidin and puromycin (0.5 μg/ml for U2OS cells, 2.5 μg/ml for MCF10A and OHSU-SCC-97 cells, and 5 μg/ml for RPE-1 *TP53^-/-^*cells). The efficiency of pooled KOs was assessed as described in the “Analysis of Cas12a gene editing” section (Extended Data Table 1), as well as by western blotting (Extended Data Fig. 17).

To generate U2OS Halo-Pol θ RHINO-mNeonGreen cells, previously published U2OS Halo-Pol θ cells^102^ were transfected in 6-well plates with 1 µg gRNA/Cas9 and 1 µg repair donor plasmid using FuGene 6. mNeonGreen positive cells were enriched by fluorescence activated cell sorting. Proper insertion of mNeonGreen in the RHNO1 locus was confirmed by genomic PCR and Sanger Sequencing (Supplementary Table 15-2 and Extended Data Fig. 20k). Expression of Rhino-mNeonGreen was validated by western blotting using an anti-mNeonGreen antibody (Extended Data Fig. 20j).

### Genetic interaction (GI) library design

The 233 DDR library genes were selected based on our understanding of the DDR, with the goal of maintaining representation of the numerous DDR pathways (Fig. 1c and Supplementary Table 1). A subset of essential DDR genes was not included in this library due to library size constraints and the limited utility of essential genes in KO-based GI scoring. To enhance the design of the overall library, multiple control groups were included to ensure a more accurate assessment of screening quality. These control groups consist of: 1) single control genes from both essential and nonessential sets^188,189^. We chose 50 genes from the essential and nonessential sets, leveraging the selection criteria previously used in^18^, which focused on genes with 1:1 orthologs in rat and mouse. 2) Essential gene pairs from the above essential genes. 3) A gold standard set of paralog pairs and their 26 individual genes that have been shown to exhibit synthetic lethality in different backgrounds^16^. Notably, *ERCC2* was used as a reference essential gene and gene pairs with *ERCC2* were dropped from the downstream analysis of GI scores.

We utilized CRISPick to select the top four predicted enCas12a gRNAs for each gene. Selected gRNAs were then shortened to 20 bp by removing 3 nt from the 3’ ends. For each pair of genes in the library, A and B, we created three arrays targeting either gene A, gene B or genes A and B. Single gene arrays contain permutations of the four available gRNAs targeting the specific gene (Fig. 1a). For gene pairs, three arrays were designed using unique combinations of two gRNAs for each gene (Fig. 1a).

After gRNA selection, library oligos were designed by adding 5’ direct repeats (DR) to each gRNA. For the first gRNA in the array, the DR is included in pRDA_052 and was not included in the library oligos. Four unique direct repeats were selected to reduce sequence similarity that can lead to unwanted recombination within arrays or low sequence diversity during next-generation sequencing^13,22^. 5’ and 3’ sequences were added to each array to allow for amplification and restriction digest cloning into a modified version of the pRDA_052 plasmid (pSAM112-UMI).

### GI library cloning

Final oligo sequences represented as: 5’-[forward primer binding site]-CGTCTCgagat-[4-gRNA array]-ttttttgaatgGAGACGat-[reverse primer binding site]-3’, with BsmBI/Esp3I recognition sites in capital letters (Supplementary Table 2), were ordered as a pool synthesized by Agilent (Santa Clara, CA). Primers SAM775 and SAM776 (Supplementary Table 15-2) were used to amplify the oligonucleotide pool in 12 x 25 μl PCR reactions employing Herculase II Fusion polymerase (Agilent) following the manufacturers protocol, with 1 ng of oligonucleotide pool and 2x (0.5 μM) primer concentration used for each reaction. Following PCR cycling conditions were used: 98 °C 2 min; 12 cycles of [98 °C 10 s, 69 °C 20 s, 72 °C 30 s]; 72 °C 3 min. PCR products were pooled and purified with PCR DNA Fragments Extraction Kit (IBI Scientific) according to manufacturer’s instructions before being assembled into a version of the pRDA_052 backbone (pSAM112-UMI) that was modified to contain a 6 bp unique molecular identifier (UMI) downstream of the most 3’ BsmBI restriction site. Cloning was done via Golden Gate assembly using Golden Gate Enzyme Mix (BsmBI-v2, New England Biolabs) in 10 x 20 μl reactions with 0.05 pmol of vector and 0.1 pmol of insert as follows: 60x cycles of [42 °C 5 min, 16 °C 5 min]; 60 °C 1 h. The Golden Gate product was then isopropanol precipitated and electroporated into Endura electrocompetent cells (DUOs) and grown at 30 °C overnight on agar plates supplemented with 100 μg/ml carbenicillin (GoldBio). An estimate of 6000 colonies per library construct was recovered. Plasmid DNA was extracted via column purification using ZymoPURE™ II Plasmid Maxiprep Kit (Zymo Research) according to manufacturer’s instructions.

### Lentivirus production

Screen library lentiviruses were generated using Transporter 5 transfection reagent (Polysciences) according to manufacturer’s protocol. In brief, the night before transfection 4 x 10^6^ HEK293T cells were plated in 10 cm plates. On the day of transfection, fresh antibiotic-free DMEM supplemented with 2% Fetalgro (Rocky Mountain Biologicals) was added to the cells. For transfection of each plate, 6 μg of the plasmid library along with plasmids encoding the lentiviral packaging components gag-pol, VSV-G, rev, and tat (0.6 μg, 1.2 μg, 0.6 μg, and 0.6 μg, respectively) were mixed in 0.5 ml of 150 mM NaCl solution. 48 μl of Transporter 5 was added to the mixed plasmids and vortexed. This solution was incubated for 20 min at room temperature before being added dropwise to cells. The morning after transfection, cell media was replaced with fresh antibiotic-free media supplemented with 10% Fetalgro. 48 h later, viral supernatant was collected, and cellular debris was removed by filtration through a 0.45 μm filter. Virus was stored at −80 °C. Small scale lentivirus was produced using the same protocol in 6-well plates.

### Determination of pooled screening lentiviral titer

Lentiviral titer was estimated by serial dilutions of virus followed by infection of cells in the appropriate plate format for the given experiment. Infections were done in media supplemented with 8 μg/ml polybrene (Sigma-Aldrich). One day after infection, the corresponding antibiotic selection was added to cells. Once selection was completed, as determined by total cell death in a control plate that received no virus, cells were counted for each of the samples receiving a different lentiviral dose. Cell counts from infected cells were compared to a second control plate that was not infected or selected. The viral dose closest to 30% survival compared to the unselected control was then used in Cas12a combinatorial KO screens.

### GI screening

Screening was conducted in three replicates starting from independent infections in two clonal cell lines constitutively expressing enCas12a, MCF10A and RPE-1 *TP53^-/-^* cells. For infection of each replicate, 8 x 10^6^ MCF10A cells per plate were seeded in 33 x 15 cm plates and 5 x 10^6^ RPE-1 *TP53^-/-^* cells were seeded in 35 x 15 cm plates. Density and plate numbers were chosen to ensure 1000 MCF10A cells per gRNA and 500 RPE-1 *TP53^-/-^* cells per gRNA following infection with a transduction efficiency of 30%. The morning after plating, cells were infected with the lentiviral library at a volume predetermined to yield a 30% transduction efficiency. Infections were done in media supplemented with 8 μg/ml polybrene (Sigma-Aldrich). For MCF10A screen, 24 h after infection, virus containing media was replaced with fresh media. 9 h later, puromycin was added directly to the plates at a final concentration of 8 μg/ml. MCF10A cells were collected at 1000x representation 15 h after selection was added.

For RPE-1 *TP53^-/-^*screen, cells were split into 20 μg/ml puromycin selection 24 h after viral transduction and collected at 500x representation 72 h later, when selection was finished. MCF10A and RPE-1 *TP53^-/-^*cell pellets were frozen at −80 °C and used as the screen T0. During the screen, MCF10A cells were passaged every 3 days at >1000x coverage into media supplemented with 2.5 μg/ml puromycin and 2.5 μg/ml blasticidin, while RPE-1 *TP53^-/-^* cells were passaged at >500x coverage into media with 5 μg/ml puromycin and 2.5 μg/ml blasticidin. At the end of the screen, genomic DNA (gDNA) was extracted from cell pellets for samples collected at T0, T9, and T17 (MCF10A)/T18 (RPE-1 *TP53^-/-^*) (Fig. 1d). gDNA was extracted at either >1000x or >500x representation for MCF10A and RPE-1 *TP53^-/-^*cells, respectively, using Blood & Cell Culture DNA Maxi Kit (Qiagen) as per the manufacturer’s protocol.

Screen gDNA was sequenced using a two-stage PCR library protocol. The first PCR (PCR1) amplified gRNA constructs added variable length (1-5 nt) random nucleotide sequences to increase sequencing library diversity and appended a constant region of the Illumina sequencing adapters. Primers for PCR1 (Forward – SAM557-561; Reverse – SAM795, SAM800-SAM803, Supplementary Table 15-2) have the following design: forward primer: 5’-ACACTCTTTCCCTACACGACGCTCTTCCGATCT-[N 1-5 nt]-tctagttacgccaagcttgc-3’; reverse primer: 5’-AGACGTGTGCTCTTCCGATCT-[N 1-5 nt]-gtggaaaggacgaaacaccg-3’. For each MCF10A sample, 570 μg of gDNA was used as starting material to maintain a representation of 1000x based on the consideration that the genome of 10^6^ diploid cells weights ∼6.6 μg. For RPE-1 *TP53^-/-^*samples, the amount of gDNA was reduced by half to maintain 500x representation. For each sample, PCR1 was run in 57 (MCF10A) or 29 (RPE-1 *TP53^-/-^*) x 100 μl reactions using 10 μg of gDNA, 1X Ex Taq Buffer (Mg2+), 800 μM dNTPs, 0.5 μM forward primer, 0.5 μM reverse primer, 7.5 μl DMSO and 7.5 units Ex Taq polymerase (Takara Bio). The following PCR cycling conditions were used: 18 cycles of [98 °C 10 s, 62 °C 30 s, 72 °C 40 s]; 72 °C 3 min.

The second PCR (PCR2) appended the rest of the Illumina adapter sequence, as well as unique Illumina indexes. Primers for PCR2 have the following design: forward primer: 5’-CAAGCAGAAGACGGCATACGAGAT-[i7 index]-gtgactggagttcagacgtgtgctcttccgatct-3’; reverse primer: 5’-AATGATACGGCGACCACCGAGATCTACAC-[i5 index]-acactctttccctacacgacg-3’ (Supplementary Table 15-2). For each sample, PCR2 was run in 3 x 100 μl reactions using 2 μl of PCR1 product, 1X Ex Taq Buffer (Mg2+), 800 μM dNTPs, 0.25 μM forward primer, 0.25 μM reverse primer and 2.5 units Ex Taq polymerase (Takara Bio). The following PCR cycling conditions were used: 8 cycles of [98 °C 10 s, 67 °C 30 s, 72 °C 40 s]; 72 °C 3 min. After PCR2, each sample was run on 1% agarose gels and the resulting 361-369 bp product was excised and gel purified (Qiagen) according to manufacturer’s instructions. 150 bp paired end sequencing of pooled libraries was conducted by CD Genomics at a target sequencing depth of 1000x (MCF10A) or 500x (RPE-1 *TP53^-/-^*) library representation. Read counts for each sample were quantified using bowtie2 alignment to a reference file of all possible gRNA arrays present in the library. gRNA arrays with 1 or fewer mismatches were considered in the final counts.

### Simulation of homozygous double knockout (dKO) rates in diploid cells

Homozygous dKO rates for diploid cells were modeled based on the editing efficiency of individual gRNAs targeting the genes of interest (Extended Data Fig. 1a). Editing efficiencies represent the fraction of gRNA-induced KO alleles versus WT alleles in a theoretical population of cells. For example, an editing efficiency of 0.8 implies 80% of target alleles have been disrupted in the population. In the simulation, the considered alleles are assigned either KO status or WT status randomly at a frequency determined by the tested editing efficiency (editing efficiency of 0.8 means 80% chance that an allele will be assigned KO). In the 4-gRNA system in Extended Data Fig. 1b, an allele has two opportunities to be disrupted, and homozygous KO is determined if both alleles are assigned KO status by either of the two gRNAs. Homozygous dKO is achieved when both alleles of the two genes are lost at any position (gRNA1 or gRNA2). Simulations were run 10^6^ times for each combination of indicated editing efficiencies.

### GI screen data analysis

The analysis begins by processing raw read count data from each sample, where a pseudocount of one was added to all counts and normalized to the total read count per sample. Guide-level log2 fold change (LFC) was calculated as the ratio of the read count at time endpoint vs T0. To measure the assay efficacy, the in4mer library included single KO arrays targeting 50 essentials and 50 nonessential genes from the Hart reference sets (Supplementary Table 2)^188,189^. We examined the LFC density plots of the guides targeting the essential and nonessential gene controls to assess if these guides effectively separate the two distributions. Guide-level LFCs were averaged across guides targeting the same gene to obtain gene-level LFCs, which were then mode-centered to zero.

To calculate genetic interaction (GI) scores, we used the GRAPE (Genetic Interaction Regression Analysis of Pairwise Effects) pipeline^35^, which estimates the fitness effects of single gene KOs via a regression-based model. The predictor was a binary matrix A(i,j) that described the presence and absence of specific genes within the arrays. The columns j of this matrix corresponded to the unique genes present across all pairwise arrays, while the rows i represented the individual arrays themselves, and A(i,j)=1 if array i contains a gRNA targeting gene j, or 0 otherwise. The response vector for the regression was the mode-centered observed gene-level LFC data for all arrays, as defined earlier. The resulting regression coefficients β correspond to the single gene KO fitness effects learned from the model and are referred to throughout the manuscript as single mutant fitness (SMF) (Extended Data Fig. 3a).

Utilizing these coefficients, we then calculated the expected dKO fold change for each gene pair, assuming additivity (Extended Data Fig. 3b). A filter was then applied to remove pairs with expected LFC smaller than the minimum of the observed LFC. This filter improved the reliability of our results by excluding gene pairs whose expected LFC values fell outside the dynamic range of the assays. After filtering, the regression model was refit using the remaining observed LFCs to obtain updated SMF and expected LFC values. The raw GI score was calculated as the difference between observed and expected LFCs, representing the deviation from the additive expectation, and served as an initial estimate of genetic interaction. To account for the increasing variance at lower expected LFCs, we applied empirical Bayes normalization. For each gene pair, the local standard deviation was estimated from the 500 nearest neighbors from the expected LFC (Extended Data Fig. 3c). GI Z-scores were then measured as the raw GI divided by the local standard deviation (Extended Data Fig. 3d). Finally, statistical significance was assessed by calculating p-values for both synthetic sick/lethal (negative) and suppressor/within-pathway (positive) interactions based on the Z-distribution. These p-values were corrected for multiple hypothesis testing using the Benjamini-Hochberg procedure, yielding adjusted p-values for interaction calls. Finally, GI z-scores were multiplied by −1, so negative GI z-scores represent synthetic sick/lethal interactions, while positive GI z-scores point to suppressor/within-pathway interactions (Supplementary Table 3).

To integrate GI z-scores across time points and cell lines, we applied Stouffer’s method. For each gene pair, the GI z-score from each time point or cell line was combined into a single integrated GI z-score using the formula 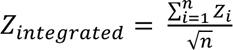, where *Z_i_* is GI z-score in each time point or cell line and *n* is the number of cell lines or timepoints being integrated.

### Single gene validation and paralog control validation

Integrated guide-level LFCs were calculated by averaging guide-level LFCs from the RPE-1 and MCF10A screens, which were obtained as described in the previous section. In Extended Data Fig. 2a, DepMap 23Q4 gene essentiality scores are compared against the integrated single gene LFC from the RPE-1 and MCF10A dataset at the late timepoints (T18 ant T17, respectively). In Extended Data Fig. 2b, expected LFCs for each gene-pair were calculated as the sum of the single gene LFCs for each gene-pair. The expected LFC and observed LFC were then plotted (Extended Data Fig. 2b).

### Correlation between RPE-1 and MCF10A screens

GI z-scores from RPE-1_integrated and MCF10A_integrated datasets were matched by gene pair to assess cross-cell line reproducibility. Pearson correlations were computed on the full dataset and on the subset of 405 pairs reaching FDR < 0.1 in both cell lines (Extended Data Fig. 2d).

### TDP1, APEX1, APEX2 screen analysis

JACKS scores from^42^ were used to calculate Δ JACKS z-scores. For each knockout background (APEX1, APEX2 and TPD1), Δ JACKS scores were calculated as the difference between the isogenic cell line’s JACKS score and the paired wild-type cell line’s JACKS score. The resulting ΔJACKS scores were then z-normalized across all target genes to produce Δ JACKS z-scores. To assess concordance between in4mer’s combinatorial screening, Δ JACKS z-scores were compared to z-normalized GI z-scores using Pearson correlation (Extended Data Fig. 2e).

### Identification of cancer-relevant genes in the GENIE database

GENIE data was filtered to include samples with deleterious mutations predicted by the SIFT (Sorting Intolerant From Tolerant) algorithm within the coding sequence of a gene^190^. All SIFT predictions with the term deleterious (i.e. “deleterious” or “deleterious low confidence”) were considered. To determine the number of patients with a mutation in each gene, patient IDs from GENIE were matched with mutation IDs and a gene was counted a single time regardless of the number of mutations present in a single patient. Cancer-associated DDR library genes were selected if they were in the top quartile of genes mutated in patients within the GENIE database (Extended Data Fig. 4a). Top cancer types in Extended Data Fig. 4b were selected based on the percentage of patients with a mutation in a DDR GI library gene (Supplementary Table 6). The total number of patients with a specific cancer annotation and a mutation in a DDR GI library gene were divided by the total number of patients with the same type of cancer. Next, cancer types with less than 50 patients assigned to them were removed and the top cancer type for each cancer associated DDR GI library gene was selected. GENIE clinical sample data and mutation data was obtained from release 15.1-public.

### Identification of disorder and cancer relevant genes in the ClinVar database

ClinVar variants were filtered to contain pathogenic germline mutations and unique MONDO IDs for each gene in the DDR GI library. After the identification of pathogenic germline mutations in the DDR library genes, MONDO ID names were used to stratify the ClinVar terms based on their association with cancer or a non-cancer genetic disorder (Extended Data Fig. 5a,b and Supplementary Table 6). After sorting of the ClinVar terms, the total numbers of unique terms per gene were tallied. The ClinVar database was accessed on 7/2/2024.

### Identification of synthetic lethal and buffering interactions for GENIE- and ClinVar-associated genes

Cancer-associated genes identified from GENIE database and disorder-related genes identified from ClinVar were used to define query gene sets for synthetic lethal and suppressor interactions plots, respectively (Extended Data Fig. 4c and 5c). A significance threshold of FDR < 0.001 was applied to both plot types, with the top 70 interactions labeled. For the suppressor interactions plot, two extra filters were applied. First, to remove hub genes with many positive interaction partners, the number of positive interactions (FDR < 0.1) per gene was z-score normalized and genes with a z-score ≥ 1 were removed. Second, interactions between genes within the same protein complex/pathway (epistatic) were excluded using cluster odds ratio > 0.

### Precision-recall analysis

Pairwise GI profiles correlations were calculated using the Pearson correlation between each gene’s interactions z-score profiles across all library genes. Essential genes (SMF below the 10^th^ percentile) were excluded from the analysis. As a baseline, pairwise Pearson correlations were also computed from DepMap dependency profiles (24Q4) across cell lines for all the genes included in our in4mer library. Protein complex membership was annotated from the Complex Portal dataset for human protein complexes (data accessed on 1/22/2026). Precision-recall curves were generated using the R yardstick package, with gene pairs ranked from highest to lowest GI profile correlation. At each rank position, precision (fraction of top-ranked pairs annotated as protein-complex) and recall (fraction of all protein complexes recovered up to that rank) were computed. The area under the precision-recall curve (PR-AUC) was computed using the yardstick::pr_auc() as a summary statistic of overall ranking performance.

### GI map and module detection

Gene modules were identified by clustering genes with similar GI profiles. To improve robustness to perturbations in the input data, we adopted a parameter sweep approach to quantify the consistency of gene co-clustering. Rather than relying on a single clustering solution, we measured how frequently gene pairs co-clustered across multiple filtered subsets of the GI dataset. This co-clustering frequency, or GI odds ratio, was used as a measure of clustering stability.

Filtered datasets were generated by systematically varying two filtering parameters: GI strength (z-score thresholds from 1.5 to 4.0; step size = 0.005) and GI connectivity (minimum number of significant interactions per gene from 1 to 30; step size = 1). Each parameter combination yielding a distinct filtered gene set with more than 50% of the original gene library was retained, resulting in a collection of overlapping but distinct representations of the interaction landscape.

For each filtered gene set, the symmetric GI z-score matrix was independently clustered using HDBSCAN^191^ (metric = cosine; min_cluster_size = 2; max_cluster_size = 15; alpha = 1). For each gene pair, we recorded the number of times the genes were assigned to the same cluster across all filtered datasets. This value was normalized to generate a GI odds ratio ranging from 0 (never co-clustered) to 1 (always co-clustered), defined as:

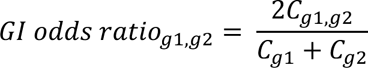

Where *C_g1,g2_* denotes the number of co-clustering events for genes *g_1_* and *g_2_*, and *C_g1_* and *C_g2_* denote the total number of clustering assignments for each gene. This procedure yielded a gene-by-gene co-clustering matrix.

The resulting co-clustering matrix was used for hierarchical clustering (Ward’s linkage, Euclidean distance) to define gene modules. The number of modules was selected based on silhouette analysis over k=2–100 clusters. Resulting heatmaps displaying the GI z-score matrix, cluster odds ratio, and SMF are shown in Extended Data Fig. 6, 7, and 8.

### Module GI score calculations

Module GI scores were calculated as the mean of all GI z-scores between the two sets of modules (Supplementary Table 7). As an example, the module GI score between the *GEN1, CIP2A,* and *RHNO1* module and the 10 gene FA module shown in Fig. 2a is the mean of all 30 (3 x 10) GI z-scores between the two sets of modules (Extended Data Fig. 6c). For the calculation of gene-module GI scores in Fig. 2c and Extended Data Fig. 12c, 13c, and 14g,h, the same approach was used but with the relevant single gene’s GI z-scores considered (Supplementary Table 7). For example, the negative gene-module interaction score for *SMARCAL1* with the *CENPS, CENPX* module is the mean of the two interactions in the module with *SMARCAL1*. For module-module and gene-module interactions, significance was assessed by standardizing GI scores to z-scores across all pairs and converting these to two-sided p-values assuming a normal distribution. For gene-module interactions, resulting p-values were adjusted for multiple testing using the Benjamini–Hochberg method, and interactions with a false discovery rate (FDR) ≤ 0.1 were considered significant. For module-module interactions multiple testing correction was not performed, and raw p-values ≤ 0.1 were considered significant.

### Module-module interaction filtering

To define a filtered module-module interaction set with a controlled number of high-scoring interactions, we implemented an iterative thresholding procedure on the module-module GI score. For each candidate threshold, interactions were filtered based on the absolute module-module GI score, and the number of unique clusters represented was quantified. We then performed a stepwise search to identify cutoff bounds that yielded a target number of modules. Starting from an initial threshold of 0, the cutoff was incrementally increased until the approximate number of desired modules was reached. Unless otherwise specified, a target of 15 clusters and a step size of 0.01 were used. This approach ensures that the filtered interaction set contains a controlled and comparable number of contributing clusters across analyses while prioritizing the strongest interactions.

### Structural predictions using AlphaFold2

Sequences of genes identified in the genetic modules defined for the RPE-1_integrated, the MCF10A_integrated, and RPE-1/MCF10A_integrated datasets were used as inputs for mmseqs2 homology search program^192^ to generate a multiple sequence alignment (MSA) against the UniRef30 clustered database^193^. Homologs sharing less than 25% sequence identity with their respective query and less than 50% of coverage of the aligned region were not kept. In every MSA, in case several homologs belonged to the same species, only the one sharing highest sequence identity to the query was kept. Full-length sequences of the orthologs were retrieved and re-aligned with mafft^194^. In total, 8,305 pairwise interactions were calculated using AlphaFold2 (AF2) between proteins inside and outside the modules (Supplementary Table 8). For this, their individual MSAs were concatenated following a paired and unpaired mode^195^ and used as inputs to generate 5 structural models using a local version of the ColabFold v1.5 interface^196^ running 3 iterations of the AF2 v2.3 algorithm^197^ trained on the multimer dataset^198^. To rate the quality of the models, four scores were provided by AF2, the pLDDT, the pTM score, the ipTM score and the model confidence score (weighted combination of pTM and ipTM scores with a 20:80 ratio)^198^. An average ipTM score was calculated for each protein pair across the 5 AF2 models (Supplementary Table 8 and at https://ddr-genetic-interactions.ciccialab-database.com/). For genes assigned to clusters with high GI odds ratios, but for which AlphaFold2 did not predict any potential direct physical interaction, we expanded the search for candidate partners from PDB or using the BIOGRID database^199^. For each of these singleton genes, we applied AlphaFold2, following the protocol described above, to screen the potential interaction partners of all cluster members and assess whether any factors absent from the experimental genetic screen could act as bridging factors (Extended Data Fig. 10b-f).

To evaluate whether proteins in genetic modules were enriched in high-confidence predicted physical interactions, we computed p-values based on a score-informed null model using the amplitude of the AlphaFold2 ipTM score averaged over 5 models. Under the null hypothesis, we defined the probability of a true positive by chance as a function of this predicted score, using a control dataset sampling 2,968 pairs of genes outside genetic modules. The relationships between average ipTM score and null probability is as follows (0.45: 2.6%; 0.5: 1.6%; 0.55: 1.1%; 0.6: 0.0.5%; 0.65: 0.4%; 0.7: 0.2%; 0.75: 0.17%; 0.8: 0.1%; 0.85: 0.07%; 0.9: 0.03%). For each module, we counted the number of observed positive interactions and estimated the probability of obtaining this count or greater under the null model using Monte Carlo simulation. Specifically, we simulated 10^5^ random realizations of the module, where each pair was assigned a success probability according to its score and counted the number of simulated hits. The p-value was defined as the proportion of simulations with a number of hits greater than or equal to the observed count (Extended Data Fig. 9c-e and Supplementary Table 9).

### Gene over-representation analysis

Gene over-representation analysis (ORA) was performed using the g:Profiler toolset (g:GOSt) from the gprofiler2 R package. For each cluster, member genes were queried against 4 different annotation sources, including: Gene Ontology Cellular Component GO:CC, CORUM protein complexes, REAC, and KEGG. Enrichment was calculated versus a custom background gene set comprised of all DDR genes in the GI screen library. Multiple testing correction was performed using Bonferroni method, and only the most significant term per cluster was selected for reporting. Cluster cohesion was quantified by silhouette analysis after generating a distance matrix using cluster odds ratio data (distance = 1 – odds ratio). ORA pipeline was applied to MCF10A_integrated and RPE-1_integrated datasets (Extended Data Fig. 11).

### Two-color competition growth assays

To generate MCF10A enCas12a cell lines stably expressing either mCherry-NLS-HYGRO (pSAM148) or GFP-NLS-HYGRO (pSAM149) (Supplementary Table 15-5), cells were transduced with lentiviruses containing these plasmids, followed by 48 h selection with 100 μg/ml hygromycin. RPE-1 *TP53^-/-^* cells were transduced with mCherry-NLS-NEO (pSAM558) or GFP-NLS-NEO (pSAM557) (Supplementary Table 15-5). 24 h later, cells were split into 2 mg/ml G418 selection, which was complete 48 h later. MCF10A and RPE-1 *TP53^-/-^* cell populations expressing either GFP or mCherry were then selected by fluorescence-activated cell sorting (top 20%).

MCF10A and RPE-1 *TP53^-/-^*enCas12a cells expressing mCherry-NLS were plated in 10 cm dishes at a density of 2 x 10^6^ and 5 x 10^5^ cells per plate, respectively. MCF10A and RPE-1 *TP53^-/-^* enCas12a cells expressing GFP-NLS were plated in 12-well plates at a density of 7 x 10^4^ and 3.5 x 10^4^ cells per well, respectively. The next day, mCherry-NLS cells were transduced with the control lentivirus containing in4mer gRNA array targeting two olfactory receptor genes, *OR1E1* (Ctrl A) and *OR52A5* (Ctrl B) (pSAM483, Supplementary Table 15-1). GFP-NLS expressing cells were transduced with lentiviruses containing gene pair relevant in4mer gRNA arrays (Supplementary Table 15-1). For each gene pair three separate GFP pools were made: in the first pool gene 1 and the control gene *OR52A5* (Ctrl B) were targeted, in the second pool gene 2 and the control gene *OR1E1* (Ctrl A) were targeted, and in the third pool gene 1 and gene 2 were targeted (Extended Data Fig. 15a). Each in4mer gRNA array had two gRNAs targeting each gene. 24 h after lentiviral transduction puromycin was added at a final concentration of 8 μg/ml for MCF10A (24 h selection) and 20 μg/ml for RPE-1 *TP53^-/-^* (72 h selection) to select for transduced cells. After puromycin selection, GFP- and mCherry-expressing cells were mixed at a 1:1 ratio and seeded on black clear-bottom 96-well plates (Greiner Bio-One) at a collective density of 1 x 10^4^ cells per well. The next day, considered T1, plates were imaged to determine the ratio between GFP and mCherry cells. Next, MCF10A and RPE-1 *TP53^-/-^* cells were maintained and passaged for 15 days in the media containing antibiotics (MCF10A: 2.5 μg/ml puromycin, 2.5 μg/ml blasticidin, and 25 μg/ml hygromycin; RPE-1 *TP53^-/-^*: 5 μg/ml puromycin, 2.5 μg/ml blasticidin, and 500 μg/ml G418), with images acquired at T3, T6, T9, T12 and T15 (Extended Data Fig. 15a). Image acquisition was performed with an ImageXpress Nano Automated Imaging System microscope (Molecular Devices) with a 10X objective. Image analyses for segmentation and counting of GFP and mCherry positive cells was performed using the MetaXpress imaging software. Each experiment was performed in three biological replicates (independent viral transductions), with three technical repeats (independent plating of mCherry/GFP cell mixtures) per biological replicate.

For analysis of genetic interactions, a pseudo-count of 5 cells was added to each mCherry or GFP population. For each sample the ratio of GFP cells in the population was calculated [ratio = GFP cells / (GFP cells + mCherry cells)]. LFC calculations for each sample at a given timepoint were then calculated as log_2_(ratio_Timepoint_/ratio_T1_). The mean of technical replicate LFCs was used as the LFC for a given biological replicate. Each sample was then normalized to the mean LFC of control samples at the appropriate time point. Control samples were infected with in4mer gRNA arrays targeting the nonessential gene combinations *OR1E1-PPP3R2*, *CLRN1-OR52A5*, or *CLRN1*-*PPP3R2* (pSAM436, pSAM437, pSAM482, Supplementary Table 15-1). Expected GI scores were calculated from the sum of relevant single gene LFCs (Extended Data Fig. 15b). GI scores were calculated as the difference between the observed LFC and the expected LFC (Extended Data Fig. 15b). LFC values were corrected by subtracting the mean LFC of control pairs (*CLRN1*, *PPP3R2*, *CLRN1-PPP3R2*) matched by timepoint and biological replicate. A two-way ANOVA test was used to calculate p-values for each gene pair comparing the expected and observed LFCs of the three biological replicates at T15.

### Antibodies

Primary antibodies used for immunofluorescence included mouse monoclonal anti-CIP2A (Santa Cruz Biotechnology, sc-80659, 1:500), rabbit polyclonal anti-RPA32 (Bethyl Laboratories, A300-244A, 1:10,000), rabbit polyclonal anti-γH2AX (Bethyl Laboratories, A300-081A, 1:5,000), mouse monoclonal anti-γH2AX (Millipore, 05-636, 1:2,000), rabbit polyclonal anti-53BP1 (Bethyl Laboratories, A300-272A, 1:2,000), mouse monoclonal anti-cyclin A (Santa Cruz Biotechnology, sc-271682, 1:1,000), mouse monoclonal anti-PCNA (Invitrogen, MA5-11358, 1:500), mouse polyclonal anti-TRF1 (Abnova, H00007013-B01P, 1:500), rat monoclonal anti-FLAG (BioLegend, 637319, 1:500). Primary antibodies used for western blotting included mouse monoclonal anti-HA (Sigma-Aldrich, H3663, 1:2,000), rat monoclonal anti-FLAG (BioLegend, 637319, 1:1,000), rabbit monoclonal anti-FLAG (Cell Signaling Technology, 14793S, 1:1,000), rabbit polyclonal anti-MCM8 (Proteintech, 16451-1-AP, 1:2,000), rabbit polyclonal anti-MCM9 (Millipore, ABE2603, 1:5,000), mouse monoclonal anti-REV7 (Santa Cruz Biotechnology, sc-135977, 1:200), mouse monoclonal anti-REV1 (Santa Cruz Biotechnology, sc-393022, 1:500), rabbit monoclonal anti-CIP2A (Cell Signaling Technology, 14805S, 1:1,000), mouse monoclonal anti-CIP2A (Santa Cruz Biotechnology, sc-80659, 1:200), rabbit polyclonal anti-GEN1 (a gift from Steve West, 1:500), rabbit polyclonal anti-FANCA (Proteintech, 11975-1-AP, 1:1,000), mouse monoclonal anti-BRCA1 (Santa Cruz Biotechnology, sc-6954, 1:500), rabbit polyclonal anti-PRIMPOL (a gift from Juan Mendez, 1:500), rabbit polyclonal anti-mNeonGreen (Proteintech, 29523-1-AP, 1:5,000), rabbit polyclonal anti-APEX2 (Novus Biologicals, NBP2-58257, 1:500), mouse monoclonal anti-SMARCAL1 (Santa Cruz Biotechnology, sc-376377, 1:500), mouse monoclonal anti-FANCM (Novus Biologicals, NBP2-50418, 1:1,000), rabbit monoclonal anti-BABAM2 (Cell Signaling Technology, 12457T, 1:1,000), mouse monoclonal anti-ERCC1 (Santa Cruz Biotechnology, sc-17809, 1:200), rabbit polyclonal anti-ERCC4 (Fortis Life Sciences, A301-315A, 1:2,000), rabbit monoclonal anti-LIG4 (ABclonal, A11432, 1:25,000), mouse monoclonal anti-XRCC4 (Santa Cruz Biotechnology, sc-271087, 1:10,000), mouse monoclonal anti-TRF2 (Millipore, 05-521, 1:1,000), mouse monoclonal anti-Vinculin (Sigma-Aldrich, V9131, 1:100,000), rat monoclonal anti-alpha tubulin (Novus Biologicals, NB600-506, 1:20,000), mouse monoclonal anti-GAPDH (Genetex, GT239, 1:20,000).

Secondary antibodies used for immunofluorescence included goat anti-mouse Alexa Fluor 488 (Thermo Fisher Scientific, A11001, 1:500), goat anti-rabbit Alexa Fluor 488 (Thermo Fisher Scientific, A11008, 1:500), goat anti-mouse Alexa Fluor 594 (Thermo Fisher Scientific, A11005, 1:500), goat anti-rabbit Alexa Fluor 594 (Thermo Fisher Scientific, A11012, 1:500), and goat anti-rat Alexa Fluor 488 (Thermo Fisher Scientific, A11006, 1:500).

### siRNA transfection

Cells were reverse transfected with the indicated siRNAs or control firefly siRNA (Supplementary Table 15-3) using lipofectamine RNAiMAX reagent (Thermo Fisher Scientific) according to manufacturer’s instructions. Cells were collected 48 h after siRNA transfection for western blot analysis.

### Immunofluorescence

Immunofluorescence was performed 7 days after puromycin selection following lentiviral transduction of in4mer constructs. Cells were seeded in black clear-bottom 96-well plates (Greiner Bio-One) at a density of 3 x 10^4^ (MCF10A) or 2 x 10^4^ (RPE-1 *TP53^-/-^*) cells per well and immunofluorescence was carried out either the next day (for RPA32, γH2AX, 53BP1, TRF1, cyclin A, and PCNA) or 2 days after seeding (for CIP2A staining and analysis of mitotic aberrations), to increase the number of mitotic cells. When cyclin A was used as a cell cycle marker, cells were washed with PBS, followed by fixation with 4% paraformaldehyde in PBS for 10 min at room temperature and subsequent wash with PBS. Next, cells were permeabilized using 0.5% Triton X-100 in PBS for 5 min at room temperature and washed with PBS. When PCNA was used as a cell cycle marker as well as for anti-CIP2A and anti-TRF1 immunofluorescence, cells were washed with PBS, followed by incubation with ice-cold pre-extraction buffer (25 mM HEPES pH 7.4, 50 mM NaCl, 1 mM EDTA, 3 mM MgCl_2_, 0.3 M sucrose, 0.5% Triton X-100) for 5 min on ice. Next, cells were fixed with cold 4% paraformaldehyde in PBS for 15 min at room temperature, washed with PBS, and subsequently permeabilized with ice-cold methanol/acetone solution (1:1) for 5 min at room temperature, followed by incubation in PBS containing 0.5% Triton X-100 for 5 min at room temperature and wash with PBS. The subsequent steps were the same for all immunofluorescence experiments. Cells were incubated in blocking solution (5% BSA in PBS) for 1 h at room temperature, followed by overnight incubation with primary antibodies diluted in blocking solution at 4 °C. The next day, cells were incubated with secondary antibodies for 1 h at room temperature. Following both primary and secondary antibody incubations, cells were washed 3 time in PBS for 5 min each. Next, cells were counterstained with DAPI (1 μg/ml in PBS) for 5 min at room temperature and washed several times with PBS.

### Image acquisition and analysis

Immunofluorescence images were acquired with an ImageXpress Nano Automated Imaging System microscope (Molecular Devices) with a 40x Plan Apo objective at a single autofocus-directed z-position. Image analyses for segmentation and counting of DAPI nuclei, cyclin A positive (CycA+) and cyclin A negative (CycA-) nuclei, 53BP1 foci, 53BP1 nuclear bodies (NBs), RPA32 foci, γH2AX foci, TRF1 foci, and DAPI micronuclei (MN) was performed using the MetaXpress High-Content Image Acquisition and Analysis Software. Three independent biological replicates were carried out and ∼2,000 cells per sample per experiment were analyzed, unless stated otherwise. Statistical significance was assessed using an unpaired t test when comparing expected and observed values, and one-way ANOVA when more than two groups or comparisons were analyzed.

To determine the distribution of DDR foci in G1 and S/G2 cell cycle phases, cells were simultaneously stained with the anti-cyclin A antibody, where CycA-nuclei mark G1 phase and CycA+ nuclei mark S/G2 phases. Large 53BP1 foci in CycA-cells were scored as 53BP1 NBs. For analysis of micronuclei (MN) counts, small DAPI objects were scored as MN and large DAPI objects were scored as cells, based on the average area of DAPI. The results were plotted as the number of MN per 100 cells.

For cell cycle analysis, PCNA and DAPI intensities were used to determine G1, S, and G2 cell cycle phases. PCNA and DAPI intensities were calculated as the log_2_ of their integrated intensity and the hdbscan function from the dbscan R library was used to detect unique clusters. For hdbscan cluster detection, the minimum number of points in a cluster was set to 100. After assigning cells to a cell cycle phase, the percent of cells in each phase of the cell cycle was calculated as the number of cells in a cell cycle cluster divided by the total number of cells assigned to any cluster. Each experiment was run in six replicates [three independent viral transductions (biological replicates) and 2 imaging experiments per each transduction (technical replicates)].

For each experiment, expected values for cell cycle analysis and analysis of all DDR markers for gene pairs of interest were determined from fold changes for single genes within a gene pair relative to control samples. Particularly, fold changes for single gene constructs were calculated by dividing the value seen for a given gene by the mean of values observed in the control samples (*OR1E1-PPP3R2*, *CLRN1-OR52A5*, and *CLRN1-PPP3R2*). The expected fold change was then calculated as multiplication of fold changes for single gene constructs within a gene pair. Finally, the expected value was calculated as multiplication of the corresponding expected fold change by the mean of values in the control samples (*OR1E1-PPP3R2*, *CLRN1-OR52A5*, and *CLRN1-PPP3R2*).

CIP2A foci in pre-anaphase cells, spontaneous mitotic aberrations, and chromatin bridges in post-mitotic cells were quantified manually using ImageJ software (NIH, https://imagej.net). For CIP2A foci counts, at least 50 pre-anaphase cells were analyzed per sample per experiment for three independent biological replicates. Statistical significance was assessed by Mann-Whitney test. For quantification spontaneous mitotic aberrations, which included anaphase bridges and lagging chromosomes, at least 18 anaphase cells were scored per sample per experiment for three independent biological replicates. Statistical significance was assessed by one-way ANOVA test. Chromatin bridges in interphase cells were identified by the presence of DAPI-positive DNA threads connecting two nuclear bodies. Due to their variable length (∼50–200 μm)^200^, nuclei displaying unilateral DNA protrusions without an identifiable connected nucleus were also scored as chromatin bridges. Additionally, cells containing fragmented bridge material or residual DNA protrusions consistent with resolved bridges were included in the analysis. The results were plotted as the number of chromatin bridges per 100 cells. ∼2,000 cells were examined per sample per experiment for three independent biological replicates. Statistical significance was assessed by one-way ANOVA test.

### Live-cell imaging of Halo-Pol θ and RHINO-mNeonGreen

7 days prior to live-cell imaging, U2OS Halo-Pol θ RHINO-mNeonGreen cells expressing enCas12a were seeded in 6-well plates at a density of 1.25 x 10^5^ cells per well. The next day, cells were transduced with lentiviruses containing relevant in4mer gRNA arrays (CTRL (pSAM483), FANCL (pSAM402), GEN1 (pSAM417), FANCL-GEN1(pSAM465)) (Supplementary Table 15-1). Approximately 24 hours post-transduction, 2.5 µg/ml puromycin was added to select for transduced cells. The following day, cells were split into 10 cm tissue culture plates and the puromycin selection was reduced to 0.5 µg/ml. 3 days prior to live-cell imaging, transduced cells were trypsinized and seeded onto 24-well glass bottom plates. 2 days later, cells were synchronized for 16 h in G2/M with 9 µM CDK1 inhibitor (CDK1i; RO-3306) in the presence or absence of 0.4 µM aphidicolin (Extended Data Fig. 20m). The following day CDK1i was removed, Halo-Pol θ was labeled with 100 nM Halo-JFX650 for 10 minutes while DNA was simultaneously labeled with Hoechst (1 µg/ml). Samples were washed three times with complete medium after removing Hoechst and Halo-JFX650 and cells were arrested in mitosis with 10 µM S-trityl-L-cysteine (Enzo Life Sciences). Live-cell imaging of Pol θ and RHINO was performed at 37 °C in the presence of 5% CO2 and 95% humidity. Z-stack images were acquired at 240 nm intervals using a 3i spinning-disc confocal microscope with SoRa modality equipped with environmental controls, four laser lines (445, 488, 561, and 638 nm), a Zeiss C PlanApo 63x/1.42 NA objective and a Hamamatsu Orca-Fusion BT sCMOS camera. Analysis of mitotic Halo-Pol θ and RHINO-mNeonGreen foci was performed in ImageJ and foci were manually counted by visual inspection through each Z-frame. Halo-Pol θ analysis was performed as previously described^102^, where a focus was defined as static HaloTag signal persisting through at least four consecutive Z-frames that overlapped with Hoechst signal. At least 24 mitotic cells were analyzed per sample per experiment for three independent biological replicates. Statistical significance was assessed by Mann-Whitney test.

### Immunoprecipitation

Co-immunoprecipitation was performed as reported previously^116^. Briefly, RPE-1 *TP53^-/-^* cells complemented with either wild-type APOLLO (WT APOLLO, pSAM686) or TRF2-interaction-deficient mutants (Δ (417-532) APOLLO, pSAM688; L506E P508A APOLLO, pSAM689) were collected by trypsinization, washed with ice-cold PBS, and incubated in lysis buffer (50 mM Tris-HCl pH=7.5, 0.5% Triton X-100, 150mM NaCl) supplemented with protease and phosphatase inhibitors (Roche) for 30 min at 4 °C. Cell lysates were then cleared by centrifugation and cell pellets were incubated in high-salt lysis buffer (50 mM Tris-HCl pH=7.5, 0.5% Triton X-100, 500mM NaCl) supplemented with protease and phosphatase inhibitors (Roche) for 1 h at 4°C. After centrifugation, the concentration of high-salt lysates was adjusted to 150 mM NaCl and they were mixed with the corresponding low-salt lysates. The combined lysates were then incubated with anti-FLAG M2 magnetic beads (Millipore) for 4 hours at 4°C for immunoprecipitation. The beads were subsequently washed tree times with lysis buffer and bound proteins were eluted in 1x Laemmli buffer (Bio-Rad) supplemented with 2-mercaptoethanol to a final concentration of 710 mM (5%).

### Western blotting

Western blot analysis was carried out 6-9 days after puromycin selection following lentiviral transduction of in4mer gRNA constructs for all cell lines except for OHSU-SCC-974 cells, which grow slower and were collected 12 days after gRNA selection was complete. Cells were collected by trypsinization, washed with PBS, and lysed in 2x Laemmli buffer (Bio-Rad) supplemented with 2-mercaptoethanol to a final concentration of 710 mM (5%), followed by sample denaturation for 5 min at 95 °C and sonication. Then, samples were resolved either on Bis-Tris SDS-PAGE casted gels in MOPS running buffer (pH=7.7) or on precast Mini-PROTEAN TGX 4%–15% gels (Bio-Rad) in Tris-Glycine-SDS running buffer, followed by wet transfer (1 h 30 min, 400 mA) onto nitrocellulose membrane (Bio-Rad). Next, membranes were incubated for 1 h at room temperature in blocking solution (5% milk in 1x TBS containing 0.1% Tween20), followed by overnight incubation with primary antibodies diluted in 3% BSA in 1x TBS containing 0.1% Tween20 at 4 °C. The next day, cells were incubated with the horseradish peroxidase (HRP)-conjugated secondary antibodies for 1 h at room temperature. Following both primary and secondary antibody incubations, membranes were washed 3 times in 1x TBS containing 0.1% Tween20 for 10 min each. Membranes were visualized with SuperSignal West Pico PLUS Chemiluminescent Substrate (Thermo Fisher Scientific) or SuperSignal West Femto Maximum Sensitivity Substrate (Thermo Fisher Scientific) using ultra-high contrast western blotting film (MTC Bio).

For detection of Halo-Pol θ, cells were labeled with 100 nM Halo-JFX650 for 30 minutes at 37°C, followed by three washes with complete medium and incubation for 10 minutes at 37 °C, to allow any unbound ligand to diffuse out of the cells. Fluorescence was detected using the Cy5.5 filter on a BioRad Chemidoc.

### Complementation experiments

To functionally complement OHSU-SCC-974 cells, they were transduced with lentivirus carrying either wild-type FANCA (pCR1316, Addgene #111126) or empty cDNA expression vector (EV) (pCR1265, Addgene # 111094)^201^ (Supplementary Table 15-5). 24 h later, transduced cells were select with 1 mg/ml G418. Next, we generated enCas12a expressing OHSU-SCC-974 complemented cells as described in the “Cell line generation” section. Finally, these cells were transduced with lentiviruses carrying either CTRL (pSAM483) or GEN1 (pSAM417) in4mer gRNA arrays (Supplementary Table 15-1). For GEN1 complementation, MCF10A enCas12a cells were transduced with lentivirus carrying either WT GEN1 (pSAM516), CD GEN1 (pSAM512), or same cDNA expression vector with a small peptide (EV, pSAM367) (Supplementary Table 15-5). 24 h after transduction, hygromycin was added at a final concentration of 100 μg/ml to select for transduced cells. MCF10A cells stably expressing WT and CD GEN1 proteins as well as EV MCF10A cells were then infected with the CTRL Cas12a 2-gRNA construct (pSAM134) or construct targeting *FANCL*-*GEN1* (pSAM321, Supplementary Table 15-1). For REV7 complementation, MCF10A enCas12a cells were transduced with lentivirus carrying either EV (pSAM553), WT REV7 (pSAM695), Y63A (pSAM696), or W171A (pSAM697) point mutant forms of REV7 (Supplementary Table 15-5). 24 h later, cells were selected with 1 mg/ml G418. Then, EV, WT, Y63A, and W171A MCF10A cells were infected with either CTRL in4mer gRNA construct (pSAM483) or in4mer construct targeting *MAD2L2-MCM9* (pSAM470, Supplementary Table 15-1). For FANCM complementation, FaDu cells were transduced with either EV (pSAM553), WT FANCM (pSAM701), or FANCM-KR (pSAM702) lentivirus. 24 h after transduction, cells were selected with 1 mg/ml G418. After selection was finished, complemented FaDu cells were transfected with control or SMARCAL1 siRNAs. For APOLLO complementation, RPE-1 *TP53^-/-^* enCas12a cells were transduced with either EV (pSAM553), WT APOLLO (pSAM686), Δ(417-532) APOLLO (pSAM688), or L506E P508A APOLLO (pSAM689) lentivirus (Supplementary Table 15-5). 24 h after transduction, cells were selected with 1 mg/ml G418, followed by lentiviral transduction with either CTRL (pSAM483) or DCLRE1B (pSAM563) in4mer gRNA arrays (Supplementary Table 15-1). Cell viability of complemented cells was assessed by survival assays as described below.

### Survival assays

Survival assays were carried out 6 days after puromycin selection following lentiviral transduction of Cas12 gRNA constructs, unless stated otherwise. In experiments where siRNAs were employed together with Cas12a pooled KO cell lines, siRNA transfection was carried out 4 days after lentiviral transduction with Cas12a gRNA constructs and cells were seeded for survival 2 days later. For survival assays with PARG inhibitor (PARGi; PDD0017273) and USP1 inhibitor (USP1i; KSQ-4279), RPE-1 *TP53^-/-^* cells were treated on the day of seeding (5 h post-seeding) with the indicated doses of PARGi or 3.125 µM USP1i, along with the DMSO as a vehicle control (equivalent to the highest drug dose). Media containing inhibitors was refreshed after 4 days. MCF10A and RPE-1 *TP53^-/-^*cells were seeded in 6-well plates at a density of 1-2 x 10^4^ and 2 x 10^3^ cells per well, respectively, and allowed to grow for 6-8 days. Then, plates were fixed (10% methanol and 10% acetic acid in water) for 10 min, followed by staining with crystal violet (1% crystal violet in methanol) for 10 min, and thorough washes with water. After plates were fully dry, the absorbed dye was resolubilized with methanol containing 0.1% SDS for 1-2 h and the resuspended solution transferred to a 96-well plate for quantification with a spectrophotometer (λ = 595 nm). Following background subtraction, cell viability was calculated by normalizing the absorbance of each sample to the control sample.

For survival assays with JH-RE-06 inhibitor, MCF10A cells were seeded in 6-well plates at a density of 1.5 x 10^4^ per well. 24 h after seeding, cells were treated with either 1.25 μM JH-RE-06 inhibitor or the same volume of DMSO as vehicle control. After incubation for 3 days, the cells were split at 1:6 ratio and incubated with the same treatment for additional 3 days. Next, the number of viable cells per each condition was estimated by hemocytometer counts and plotted as percentage of viable cells relative to the DMSO control.

For OHSU-SCC-974 cells, survival assays were carried out 5 days after puromycin selection following lentiviral transduction of in4mer gRNA constructs. Cells were seeded in 6-well plates at a density of 5 x 10^4^ per well and were allowed to grow for approximately 3 weeks. During this time, media was replaced every 3-4 days and cells were split once 2 weeks after seeding at 1:4 ratio. FaDu cells stably expressing WT FANCM and mutant KR FANCM proteins were seeded in 6-well plates at a density of 3.5 x 10^5^ cells per well and were transfected with either control or *SMARCAL1* siRNAs. Cells were split at 1:4 ratio 2 and 4 days after seeding and were collectively grown for 7 days. The number of viable cells per each condition was estimated by hemocytometer counts and plotted as percentage of viable cells relative to the control samples. All survival data are representative of three independent biological replicates, unless stated otherwise. Statistical significance was assessed by either unpaired t test or one-way ANOVA test.

### Metaphase spreads

RPE-1 *TP53^-/-^* cells were arrested in metaphase with 50 ng/ml KaryoMAX colcemid (Gibco) for 2 h. Then, cells were harvested, washed with PBS, and incubated in 75 mM KCl for 10 min at 37 °C. Next, cells were fixed in ice-cold methanol:acetic acid (3:1) solution and cell suspension was dropped onto glass slides before staining with Giemsa solution (2 ml Giemsa Stain Stock Solution (Gibco) in 98 ml Gurr buffer (Gibco)). Slides were dried and mounted with DPX Mountant for histology (Sigma-Aldrich). Metaphase spreads were imaged using Zeiss Axio Imager Z2 microscope equipped with CoolCube 1 camera. Metafer software (MetaSystems) was used for the automated search (with 10x objective) and capture (with 63x/1.30 oil objective) of metaphases. Metaphase aberrations were quantified manually using ImageJ software (NIH, https://imagej.net) in at least 50 metaphases (containing at least 44 of the expected 46 clearly identifiable chromosomes) per sample per experiment for two independent biological replicates. Statistical significance was assessed by either Mann-Whitney test or one-way ANOVA test.

### Electron microscopy

Visualization of DNA replication intermediates by electron microscopy was performed as reported previously^202^, with some modifications. Briefly, 2.5-5.0 x 10^7^ MCF10A cells were collected, washed, and resuspended in 10 ml of ice-cold PBS containing calcium and magnesium. To cross-link genomic DNA in vivo, cell suspension was incubated with 10 µg/ml trimethylpsoralen (TMP, Sigma-Aldrich) for 5 min in the dark on ice, followed by 5 min irradiation with 365 nm wavelength UV light on a pre-cooled metal surface in a Stratalinker. This step was repeated four times. Then, genomic DNA was extracted with chloroform/isoamyl alcohol, precipitated with isopropanol, and resuspended in TE buffer (10 mM Tris-HCl pH 8.0, 1 mM EDTA). The integrity of genomic DNA was confirmed by agarose gel (0.8%). Next, 30 µg of genomic DNA were digested with 150 U of PvuII HF (New England Biolabs) for 3.5 h at 37 °C, followed by enrichment of DNA replication intermediates using BND-cellulose resin (Sigma-Aldrich) and subsequent DNA purification using Amicon Ultra-0.5 Centrifugal Filter Unit with Ultracel-100 (Millipore). Finally, DNA was processed for rotary shadowing and platinum coating using a Med20 evaporator (Leica). Image acquisition was performed at the IFOM EM facility using TECNAI12 electron microscope equipped with a GATAN camera run by Digital Micrograph software. ssDNA gap lengths were quantified at the forks and behind the forks (in nm). Values were converted to nucleotides by dividing by 0.36. At least 41 replication forks were analyzed per sample per experiment for two independent biological replicates. Statistical significance was assessed by one-way ANOVA test.

### END-seq

For END-seq, 2 plugs containing 5 x 10^6^ MCF10A cells each were collected per genotype, mixed with 1 x 10^6^ spike-in cells and embedded in agarose plugs. Plugs were processed as previously reported^127,203^. For END-se q in analysis, raw sequencing reads were trimmed using Trim Galore! (v.0.6.7) followed by adjusting the parameter “--stringency 3”. Trimmed reads were aligned to the human genome (T2T) using bowtie (v.1.3.1)^204^ with the parameters “-n 3 -l 50 -k 1”. For spike-in normalization, reads from the samples were aligned to both human and mouse genomes. The bamCoverage function from deepTools (v.3.5.5)^205^ was used to normalize read density with the parameters “-bs=1 -normalizeUsing BPM”. The scaling factor for the spike-in normalization was calculated using the number of reads mapped to the spike-in locus (mm10, chr6:41,554,106-41,556,020) of the mouse genome and the total number of reads mapped to the human genome. To generate the spike-in normalized reads, the normalized read density was divided by the scaling factor.

### END-seq peak calling and motif analysis

Following a previous study^68^, significant peaks for each sample were found for the positive strand and negative strand separately, on spike-in-normalized reads. MACS2 was run with parameters --keep-dup all --nomodel --nolambda -p 1e-5, and peaks with fold enrichment greater than 10 were kept. Peaks were defined as DSBs if the negative-strand peak was within 1,000 bp upstream of the positive-strand peak, with the summit of the two peaks between 30 bp and 2,500 bp apart. Recurrent motifs near peaks were identified with MEME. Identified motifs were located and scored using FIMO. Palindromic motif was annotated using annotatePeaks in Homer with parameter “--size 1000”.

### Analysis of Cas12a gene editing

To measure editing efficiency of Cas12a gRNA constructs, cells were collected by trypsinization 6-9 days after puromycin selection following lentiviral transduction of in4mer constructs, washed with PBS, and QuickExtract DNA Extraction Solution (Biosearch Technologies) was used to extract genomic DNA according to manufacturer’s instructions. Next, genomic regions containing gRNAs of interest were PCR amplified using primers listed in Supplementary Table 15-4 and sent for Sanger sequencing. DNA samples from cells transduced with the control in4mer construct (pSAM483, Supplementary Table 15-1), which targets two olfactory receptor genes (*OR1E1* and *OR52A5*), were used to derive wild-type reference sequences. Finally, Inference of CRISPR Editing (ICE) online tool (Synthego Performance Analysis, ICE Analysis. 2019. v3.0. Synthego) was employed to calculate the frequencies of indels and KO scores for gRNAs of interest (Extended Data Table 1).

## Extended data figures and tables

Extended data includes 25 figures and legends and 1 table.

## Data availability

All screening data are provided as supplementary material and can be additionally enquired on a dedicated website available at https://ddr-genetic-interactions.ciccialab-database.com/. AlphaFold datasets for prediction of physical interactions across all integrated datasets are also provided as supplementary material and can be enquired on the same website. All unique reagents and scripts generated in this study will be made available upon request. Sequencing data will be uploaded into the Sequencing Read Archive (SRA) upon publication.

## Acknowledgements

We thank Richard Baer for comments on the manuscript. The GEN1 antibody was kindly provided by Steve West (Crick Institute). The PRIMPOL antibody was kindly provided by Juan Mendez (Spanish National Cancer Research Centre, CNIO). FaDu cells were kindly provided by Alison Taylor (Columbia University Irving Medical Center), and OHSU-SCC-974 cells were a kind gift from the FARM repository. We thank Hiroshi Nakagawa for kindly providing pCR1316 and pCR1265 constructs, Feng Zhang for the kind gift of px330-U6-Chimeric_BB-CBh-hSpCas9 construct, and Andrew Deans for kindly providing FANCM pFastBac constructs. This work was supported by the NIH grants R01CA197774, R01CA304254, and P01CA174653 to A.C., the CRI Lloyd J. Old STAR award to A.C., the TL1TR001875 precision medicine predoctoral grant to S.B.H., the Terri Brodeur Breast Cancer Foundation 2026 grant to T.L.-D., the Warren Alpert Distinguished Scholar Fellowship to A.V., and NIH grants R35GM130119 and U01CA275886 to T.H.. R.G. was granted access to the HPC resources of IDRIS under the allocation AD010314343R1 made by GENCI. Studies conducted in the Flow Cytometry Resource of the Herbert Irving Comprehensive Cancer at Columbia University were supported by the NIH grant P30 CA013696. V.C. is funded by the Italian Association for Cancer Research (AIRC) AIRC-IG-28275 and the Italian Ministry of University and Research (MUR) under the Fondo Italiano per la Scienza (FIS 3), project DNAMET FIS-2024-05295.

Y.W.H. was supported by the 26596 AIRC fellowship for Italy. The authors would like to acknowledge the American Association for Cancer Research and its financial and material support in the development of the AACR Project GENIE registry, as well as members of the consortium for their commitment to data sharing. Interpretations are the responsibility of study authors.

## Author contributions

Conceptualization: S.B.H., A.V., T.L.-D., A.C.; Investigation: S.B.H., A.V., T.L.-D., A.T., J.-W.H., Y.W., G.L., J.R.H., N.W.; Methodology: S.B.H, A.V., T.L.-D., J.C., N.E.A., T.H., A.C.; Formal Analyses: S.B.H., A.V., T.L.-D., J.C., V.G., C.C, R.G., T.H.; Software: A.L.B., T.L.-D., X.F., T.L., R.R.; Writing-Original Draft: S.B.H., A.V., A.C.; Writing-Review & Editing: S.B.H., A.V., T.L.-D., J.C., A.T., J.-W.H., Y.W., J.R.H., N.W., C.C., T.L., G.L., V.G., N.E.A., A.L.B., X.F., A.N., R.R., R.G., J.C.S., V.C., T.H., A.C.; Supervision: R.R., J.C.S., A.N., V.C., R.G., T.H., A.C.; Funding Acquisition: S.B.H., A.V., T.L.-D., A.C.

## Ethic declarations

The authors declare no competing interests.

## Extended Data

**Extended Data Table 1.**
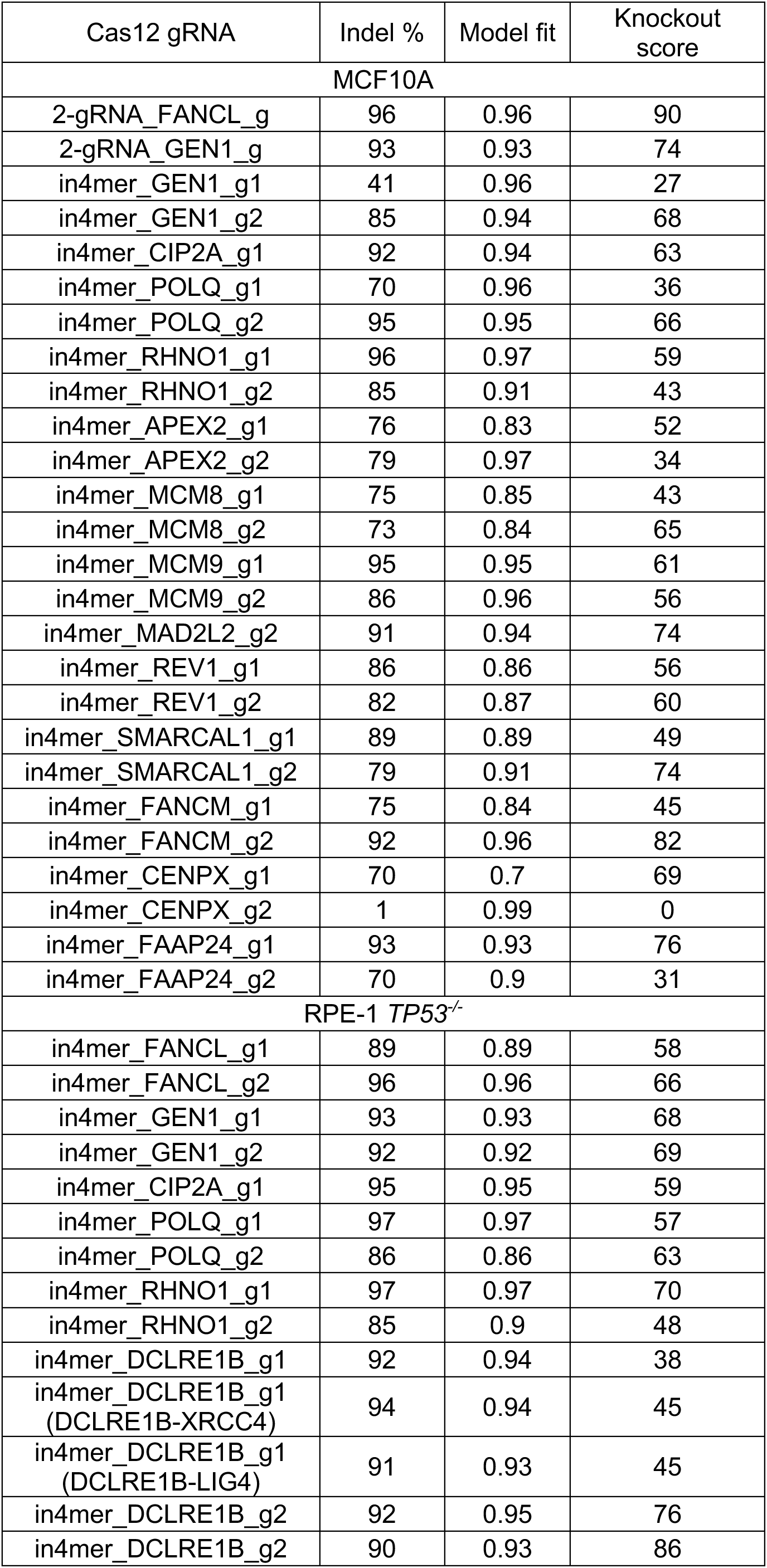
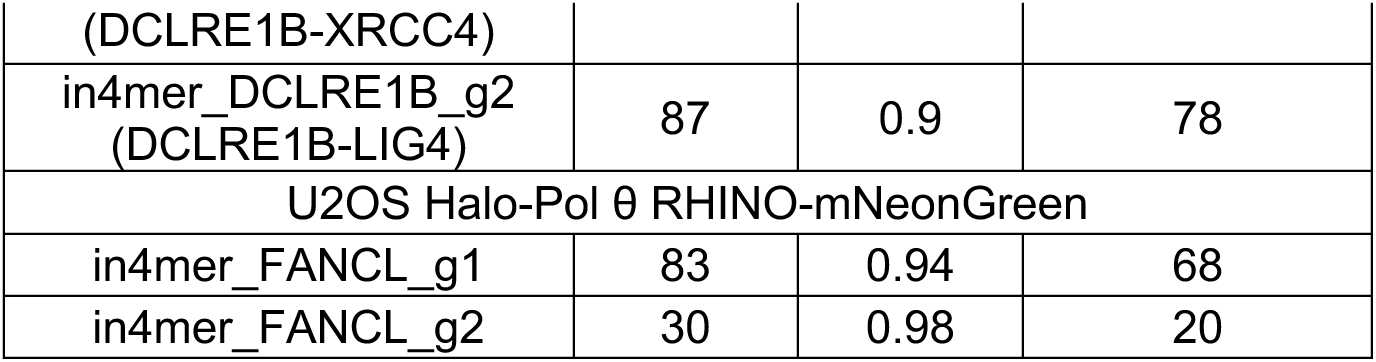
Editing efficiencies of Cas12a in4mer gRNAs. Analysis of editing efficiencies in MCF10A, RPE-1 *TP53^-/-^*, and U2OS Halo-Pol θ RHINO-mNeonGreen cells constitutively expressing enCas12a induced by the indicated gRNAs at their target loci 5-9 days after selection, as determined by Sanger sequencing and ICE analysis. “Model fit” column shows how well the distribution of indels in the sample proposed by ICE online tool matches the Sanger sequencing data. The *FANCL-GEN1* Cas12a construct consisting of two gRNAs (2-gRNA construct, one gRNA per gene) was employed in *GEN1* complementation experiments. All in4mer constructs employed for validation experiments contain 2 gRNAs per gene. The editing efficiencies for CIP2A_g2 and MAD2L2_g1 are not displayed because of the inability to amplify the genomic region targeted by CIP2A_g2 gRNA and inability to sequence the genomic region targeted by MAD2L2_g1 due to the presence of homo-polymeric regions. For *DCLER1B*, editing efficiencies of the gRNAs in the double knockouts are also displayed.

**Extended Data Fig. 1.**
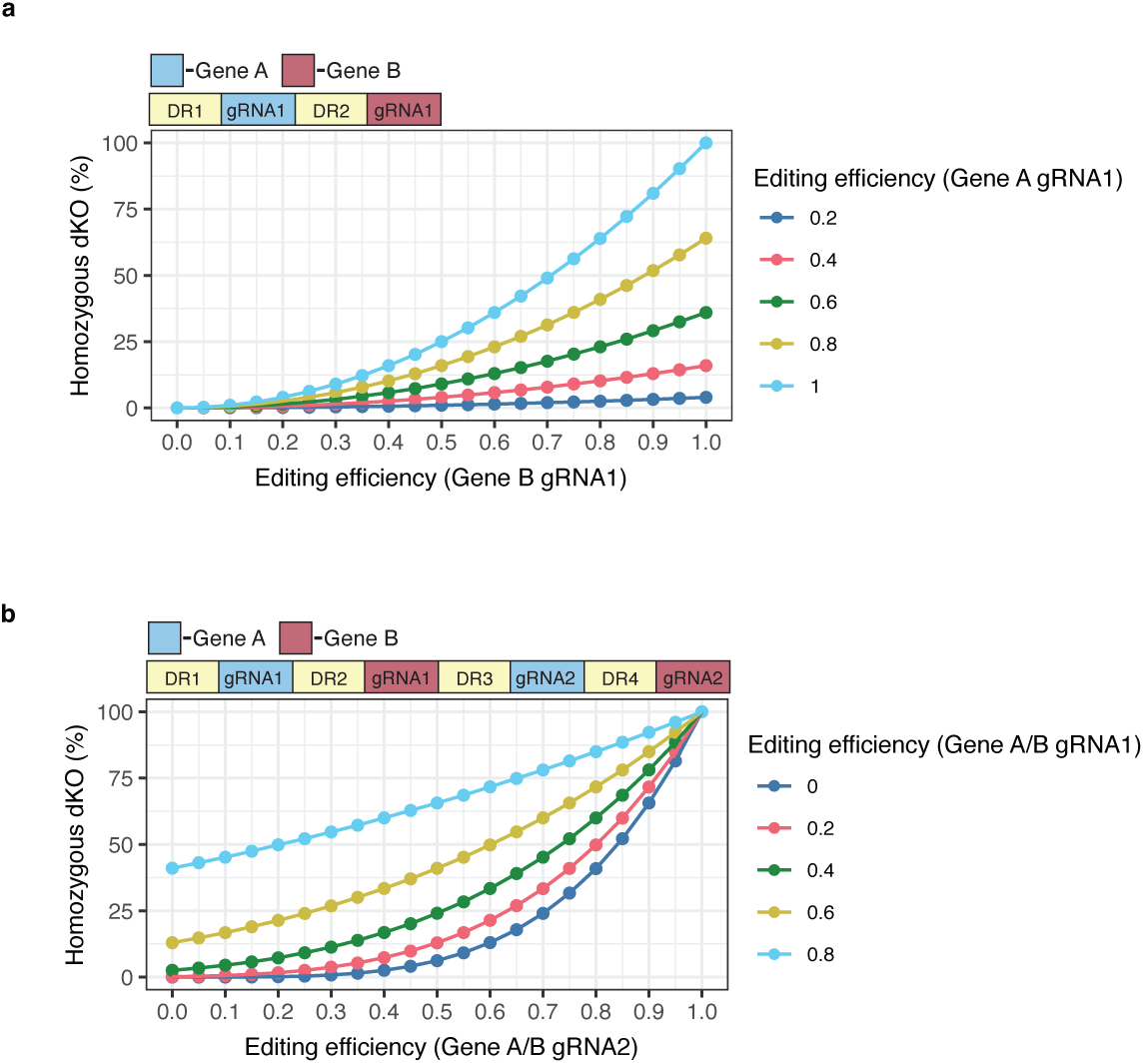
Theoretical homozygous double KO efficiency obtained using Cas12a guide RNA arrays. **a**, Simulation-based dKO rates using single gRNA targeting of two genes in diploid cells. Editing efficiency for the gRNA targeting ‘Gene A’ is shown as different color lines and editing efficiency for the gRNA targeting ‘Gene B’ is shown on the x-axis. **b**, Simulation-based dKO rates using two gRNAs targeting each gene in a gene pair. Editing efficiency for the first gRNA targeting each gene is shown as different color lines and editing efficiency for the second gRNA targeting each gene is shown on the x-axis.

**Extended Data Fig. 2.**
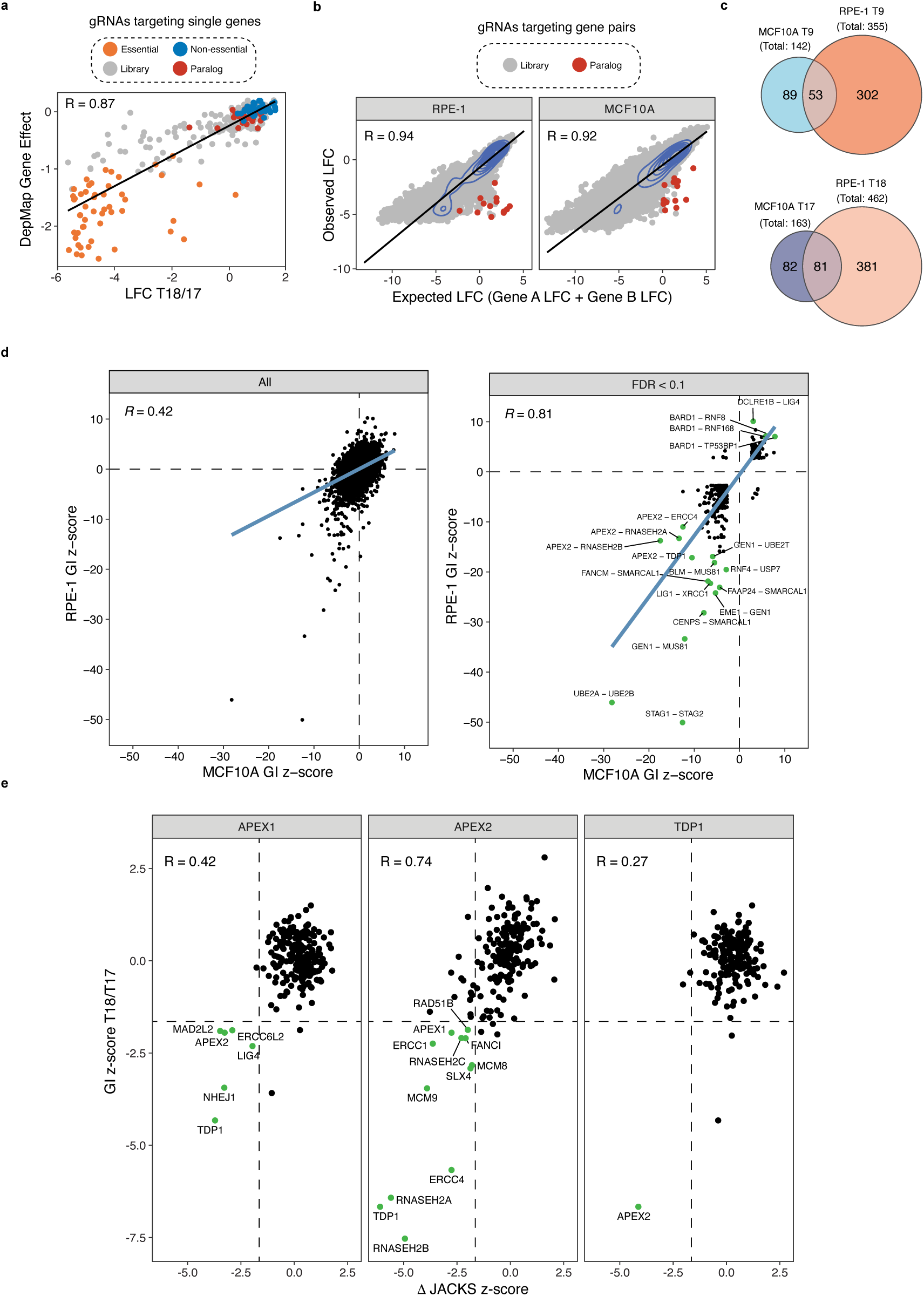
Known single gene phenotypes and genetic interactions captured by the in4mer GI screen. **a**, Correlation of single gene construct average log fold changes (LFCs) from the RPE-1 and MCF10A datasets at T18/T17 time points of GI screens with the corresponding DepMap gene effect scores. Essential, nonessential, and paralog controls from the screens are depicted in orange, blue, and red, respectively. DDR library genes are shown in grey. **b**, GI screens phenotypes of control gRNAs targeting known synthetic lethal paralog gene pairs in RPE-1 *TP53^-/-^* (left) and MCF10A (right) cell lines. Expected and observed LFCs are plotted with 13 control paralog pairs highlighted in red. DDR library gene pairs are shown in grey. **c**, Venn diagrams displaying the number of common and unique genetic interactions (FDR < 0.1) between MCF10A and RPE-1 *TP53^-/-^* cell lines at early (T9, top panel) and final (T17/T18, bottom panel) time points of the GI screens. **d**, Correlation of GI z-scores between the RPE-1_integrated and MCF10A_integrated datasets. Left: correlation computed across all gene pairs without filtering. Right: correlation after applying a threshold of FDR < 0.1. Green dots indicate the top 15 most negative and the top 4 most positive gene pairs, ranked by their GI z-score. **e**, Comparison of negative interactions identified in previously published CRISPR-Cas9 KO screens in isogenic *APEX1^-/-^*, *APEX2^-/-^*, or *TDP1^-/-^* cell lines and in our GI screens (RPE-1/MCF10A_T18/T17_integrated dataset) for the respective genes^42^. Δ JACKS score represents the difference between a gene’s phenotype in WT cells and the indicated isogenic background (Δ JACKS = JACKS score KO - JACKS score WT)^42^. Dashed lines represent the top 5% of hits in our GI screens and Cas9 KO screens. Green points indicate shared hits between the two screening approaches.

**Extended Data Fig. 3.**
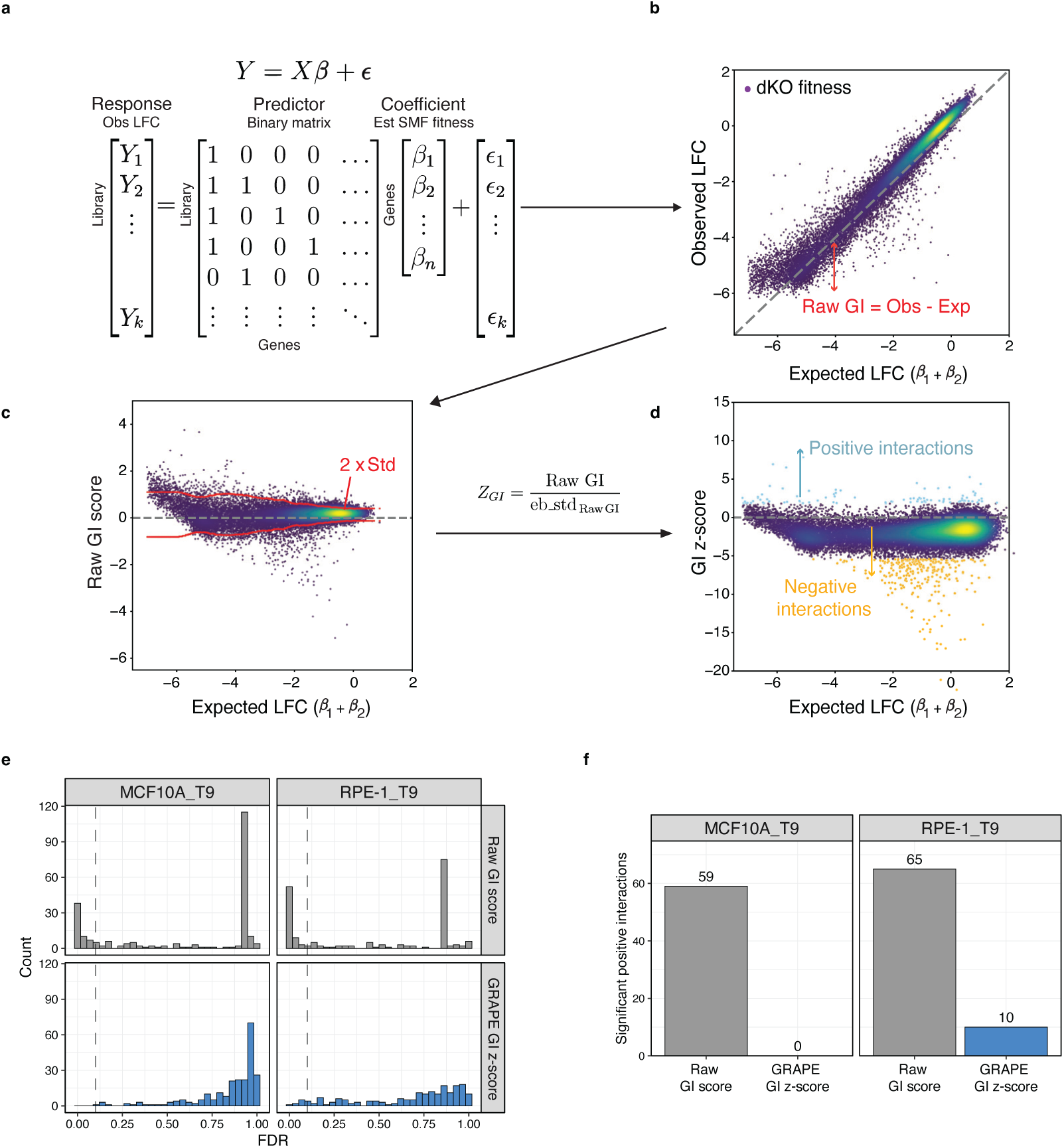
GI z-score calculation method using the GRAPE pipeline. **a**, Regression model for determining estimated (est) single gene KO fitness effects [β coefficients, or single mutant fitness (SMF) scores] from GI screening data, used to calculate expected (exp) dKO fitness effects. The predictor is a binary matrix indicating whether each target gene in the GI library is present (1) or absent (0) in each gene pair construct, and the response vector is the observed (obs) log fold change (LFC) values for gene pair constructs. k and n denote the number of gene pairs and target genes in the library, respectively. **b**, Scatter plot of observed LFCs versus expected LFCs for all gene pairs in the GI screen. Raw GI scores are computed as the difference between observed and expected LFCs. **c**, Scatter plot of raw GI scores versus expected LFCs. Red lines indicate the local standard deviation (Std) estimated from the 500 nearest neighbors in expected LFC space. Lower expected LFC is associated with greater variance. Raw GI scores were normalized using this Std, resulting in GI z-scores. **d**, Scatter plot of the final GI z-scores versus expected LFCs. **e**, Distribution of FDR values for positive genetic interactions among dual-essential gene pairs (both genes with SMF ≤ 10th percentile), comparing raw GI scores (grey) and GRAPE GI z-scores (blue) in MCF10A (T9) and RPE-1 *TP53^-/-^* (T9) datasets. Dashed vertical lines indicate FDR = 0.1. **f**, Number of artifactual positive interactions between two essential genes identified using raw GI scores compared to after local variance normalization (GRAPE analysis).

**Extended Data Fig. 4.**
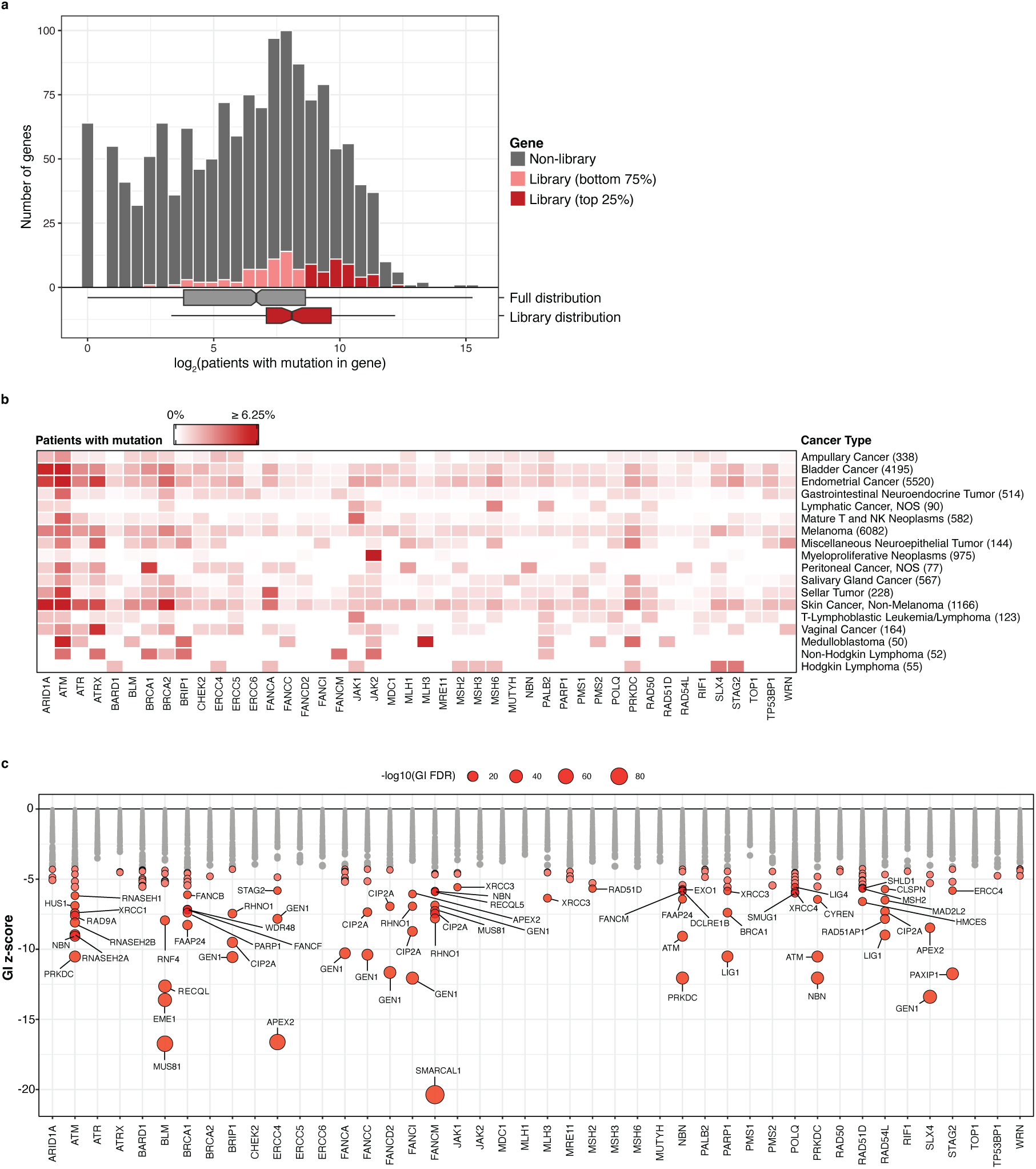
Identification of cancer-relevant synthetic lethal interactions. **a**, Histogram showing the distribution of somatic mutations across all genes in the GENIE database (AACR). Number of patients with an assigned deleterious mutation within the coding region of a given gene is shown on the x-axis. 45 DDR GI library genes among the top 25% (dark red) of mutated genes within the overall GENIE distribution (grey) are considered as cancer-relevant. Remaining DDR library genes are shown in light red. Box plots below the histogram compare the distribution of mutation frequencies for all genes in GENIE (grey) and for genes in the DDR library subset (red). Notches indicate 95% confidence intervals around the median. **b**, Heatmap showing the percentage of patients carrying somatic mutation in each of 45 cancer-relevant genes included in the in4mer DDR library across 18 cancer types from GENIE. For each gene, the cancer types with the highest mutation prevalence were identified. Only cancer types with at least 50 patients are shown (indicated between parenthesis next to the cancer type). **c**, Dot plot showing synthetic lethal interactions from the RPE-1/MCF10A_integrated dataset with cancer-relevant genes (x-axis) within our DDR GI library. Each dot represents a pairwise interaction. Dot size reflects statistical significance (-log10 FDR). Synthetic lethal interactions with FDR < 0.001 are shown in red. Gene names are only displayed for the top 70 ranked interactions.

**Extended Data Fig. 5.**
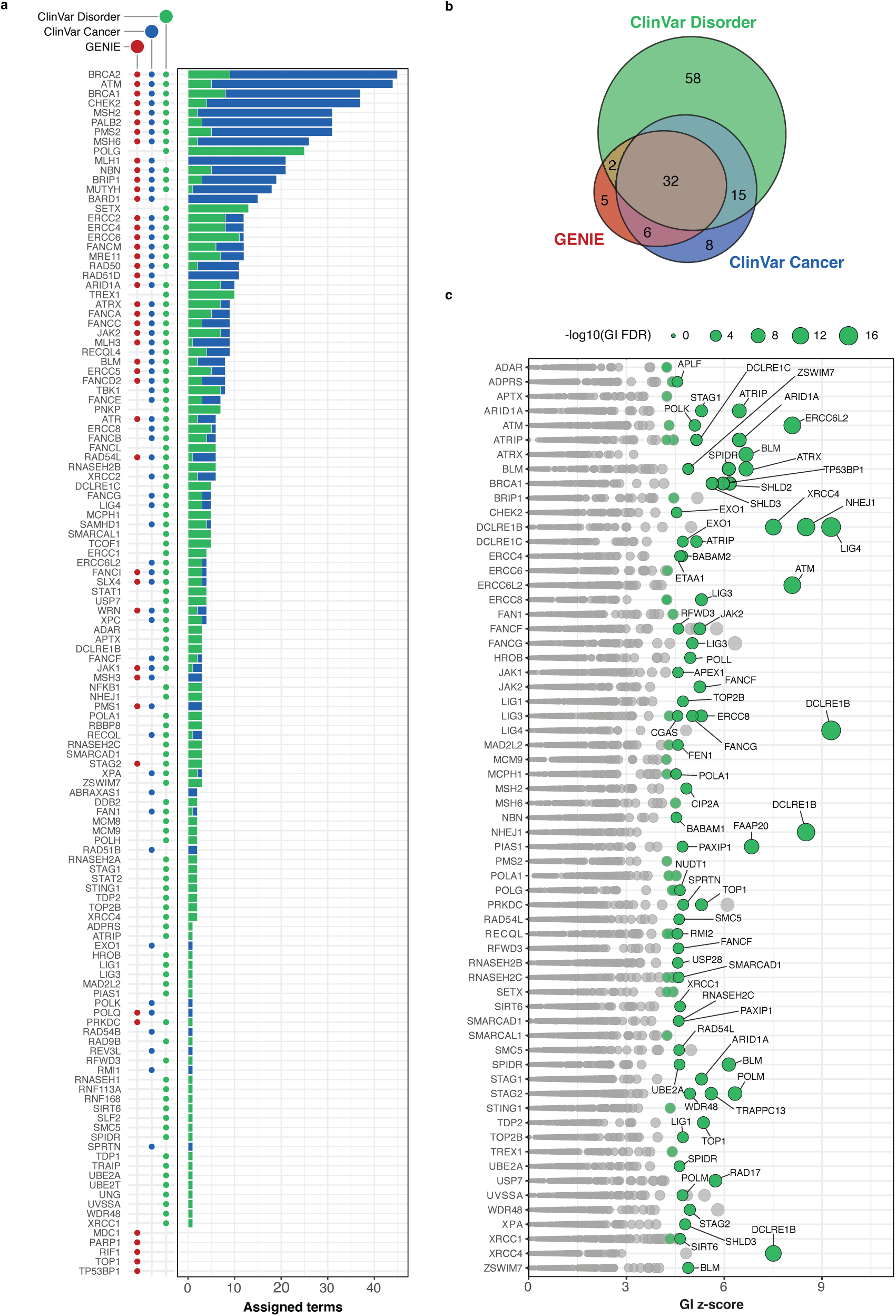
Identification of disorder-relevant positive interactions. **a**, Clinical annotations for DDR GI library genes. Y-axis lists all disorder-associated library genes. Previously identified genes within the top 25% of the GENIE database are indicated with red dots, genes with annotated germline pathogenic mutations from the ClinVar database associated with cancer and other disorders are indicated with blue and green dots, respectively. The total number of ClinVar terms representing cancer (blue) or disorder (green) are shown on the x-axis as stacked horizontal bar chart. **b**, Venn diagrams showing DDR GI library gene annotations from either GENIE or ClinVar cancer/disorder terms. **c**, Dot plot showing positive interactions from the RPE-1/MCF10A_integrated dataset with ClinVar disorder-associated genes (x-axis) within our DDR GI library. Each dot represents a pairwise interaction. Dot size reflects statistical significance (-log10 FDR). Positive interactions with FDR < 0.001 are shown in green. Gene names are only displayed for the top 70 interactions. Hub genes and same-module interactions were excluded (see methods).

**Extended Data Fig. 6.**
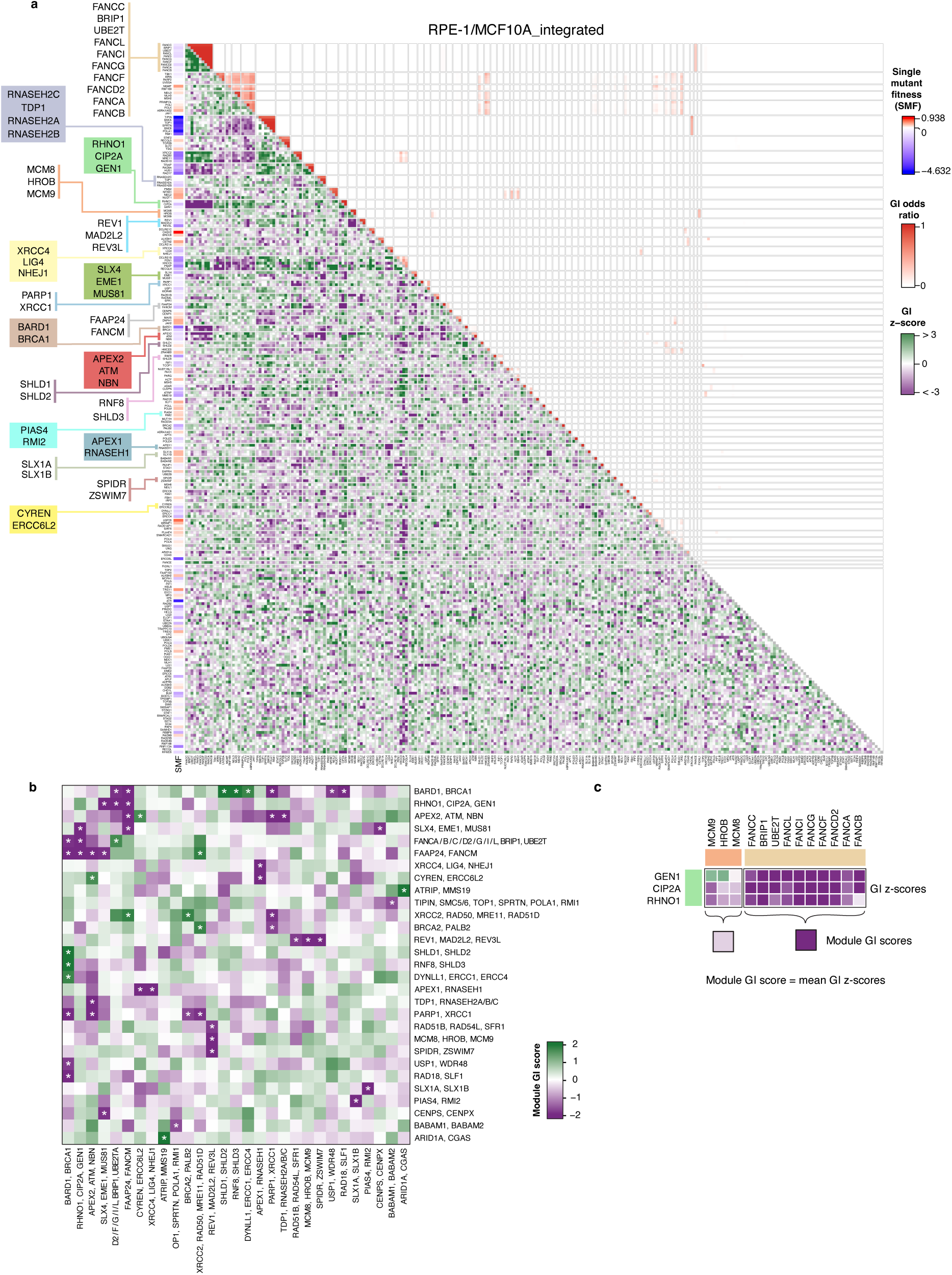
GI z-score and cluster odds heatmap for the RPE-1/MCF10A_integrated dataset. **a**, GI map for 232 DDR genes showing gene modules with shared genetic interactions in the RPE-1/MCF10A_integrated dataset. The lower heatmap triangle displays integrated GI z-scores (negative in purple, positive in green). The upper heatmap triangle displays cluster odds ratios, representing the frequency with which each two genes cluster together across multiple filtered subsets of the GI dataset (see methods). To the left of the heatmap, gene names and single mutant fitness (SMF) values are shown, and modules from the filtered GI map in Fig. 2a are highlighted. Interaction of a gene with itself is depicted as grey in the diagonal. **b**, A reduced heatmap showing the significant module-module interactions from the GI map in panel **a**. Significance is indicated (*), corresponding to a nominal two-sided p-value ≤ 0.1 under a normal approximation. **c**, Schematic illustrating that module GI scores are calculated by taking the average of all GI z-scores between two tested modules.

**Extended Data Fig. 7.**
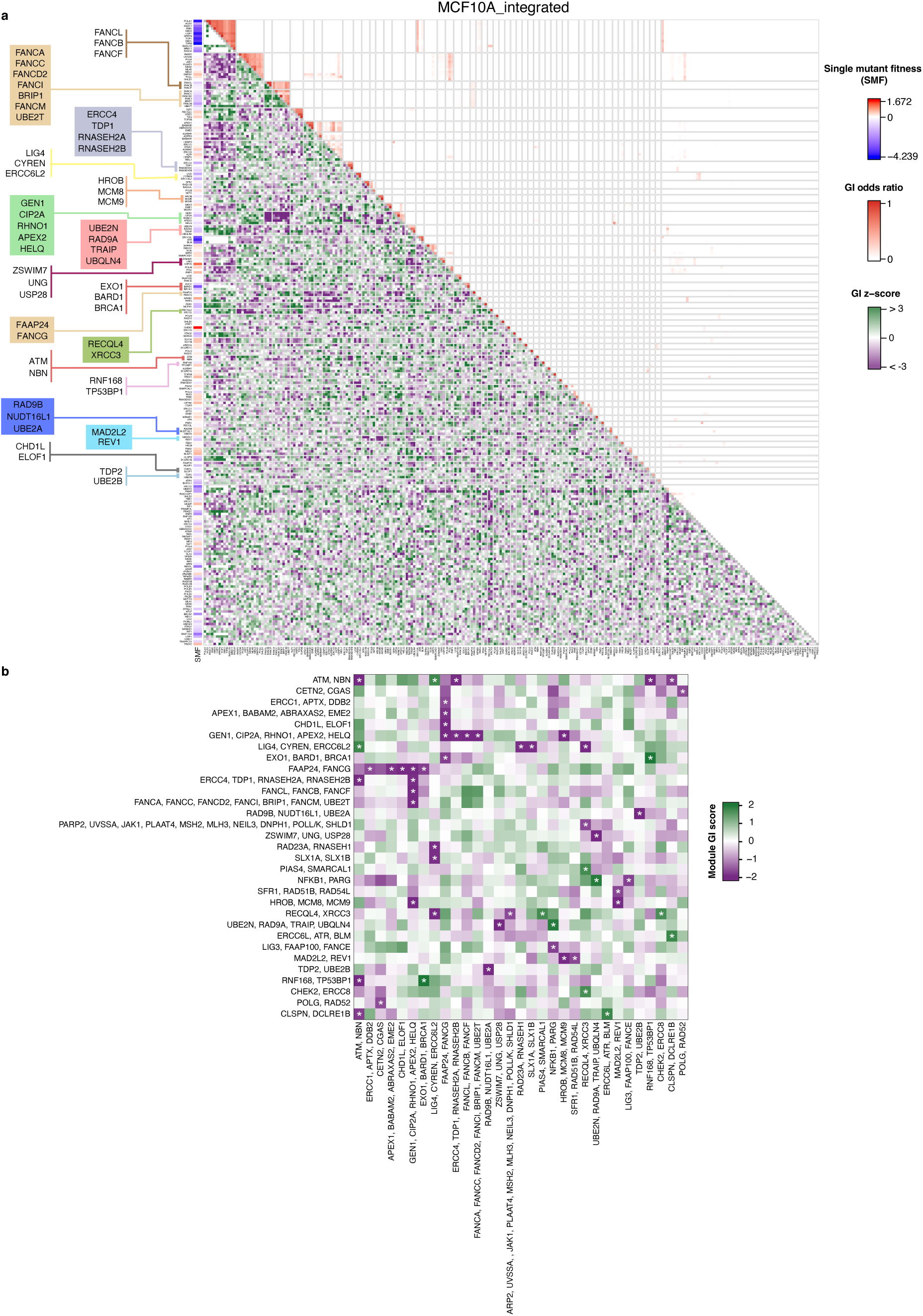
GI z-score and cluster odds heatmap for the MCF10A_integrated dataset. **a**, GI map for 232 DDR genes showing gene modules with shared genetic interactions in the MCF10A_integrated dataset. The lower heatmap triangle displays integrated GI z-scores (negative in purple, positive in green). The upper heatmap triangle displays cluster odds ratios, representing the frequency with which each two genes cluster together across multiple filtered subsets of the GI dataset (see methods). To the left of the heatmap, gene names and single mutant fitness (SMF) values are shown, and modules from the filtered GI map in Extended Data Fig. 12a are highlighted. Interaction of a gene with itself is depicted as grey in the diagonal. **b**, A reduced heatmap showing the significant module-module interactions from the GI map in panel **a**. Significance is indicated (*), corresponding to a nominal two-sided p-value ≤ 0.1 under a normal approximation.

**Extended Data Fig. 8.**
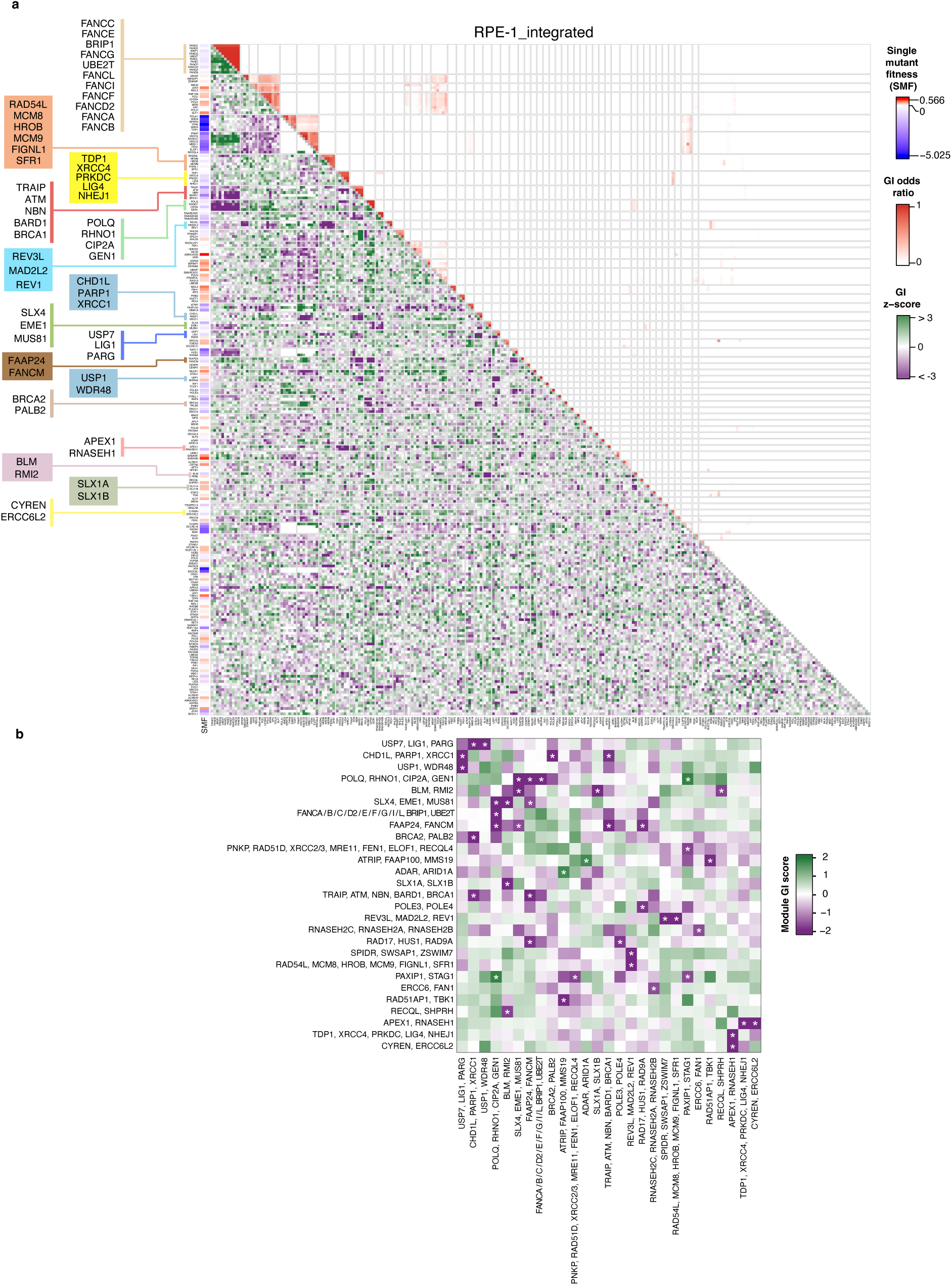
GI z-score and cluster odds heatmap for the RPE-1_integrated dataset. **a**, GI map for 232 DDR genes showing gene modules with shared genetic interactions in the RPE-1_integrated dataset. The lower heatmap triangle displays integrated GI z-scores (negative in purple, positive in green). The upper heatmap triangle displays cluster odds ratios, representing the frequency with which each two genes cluster together across multiple filtered subsets of the GI dataset (see methods). To the left of the heatmap, gene names and single mutant fitness (SMF) values are shown, and modules from the filtered GI map in Extended Data Fig. 13a are highlighted. Interaction of a gene with itself is depicted as grey in the diagonal. **b**, A reduced heatmap showing the significant module-module interactions from the GI map in panel **a**. Significance is indicated (*), corresponding to a nominal two-sided p-value ≤ 0.1 under a normal approximation.

**Extended Data Fig. 9.**
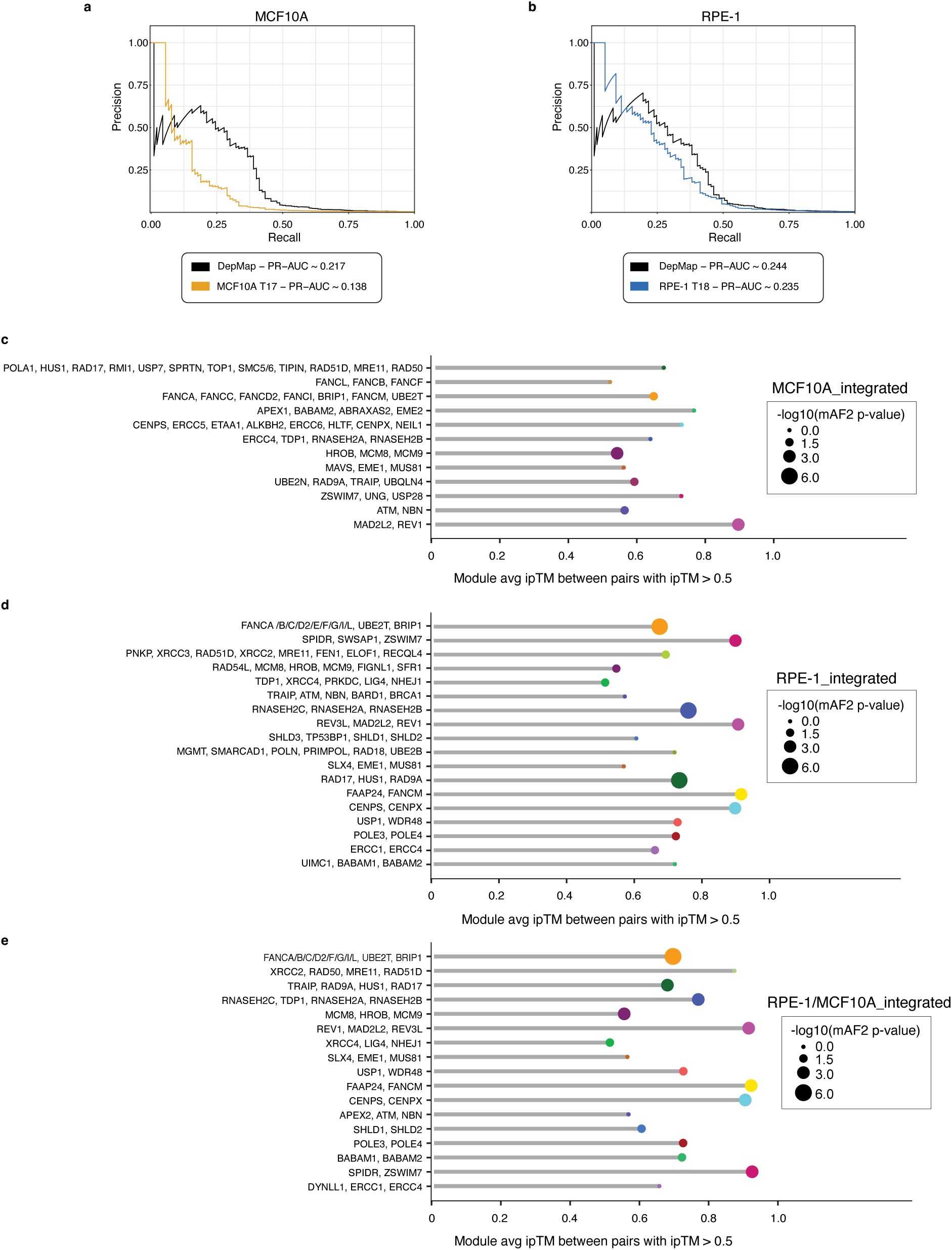
Validation of genetic interaction modules by co-complex benchmarking and predicted physical interactions. **a**, **b**, Precision-recall (PR) analysis evaluating the ability of pairwise GI profile correlations to identify gene pairs that co-annotate to the same protein complex in the Complex Portal dataset. DepMap gene-dependency profiles for the DDR GI library genes were used as a benchmark. PR-AUC stands for area under the PR curve. **a**, PR analysis for the MCF10A_T17 dataset. **b**, PR analysis for the RPE-1_T18 dataset. **c**-**e**, Pairwise AlphaFold2-multimer (mAF2) predictions among members of each genetic interaction module for the indicated integrated datasets (**c**, MCF10A_integrated dataset; **d**, RPE1_integrated dataset; **e**, RPE-1/MCF10A_integrated dataset). Shown are modules with an average interface predicted template modeling score (ipTM) > 0.5, indicating predicted direct physical association between at least a subset of module members. p-values estimating the significance of the number of direct interactions predicted within each module are indicated by circle sizes on the side.

**Extended Data Fig. 10.**
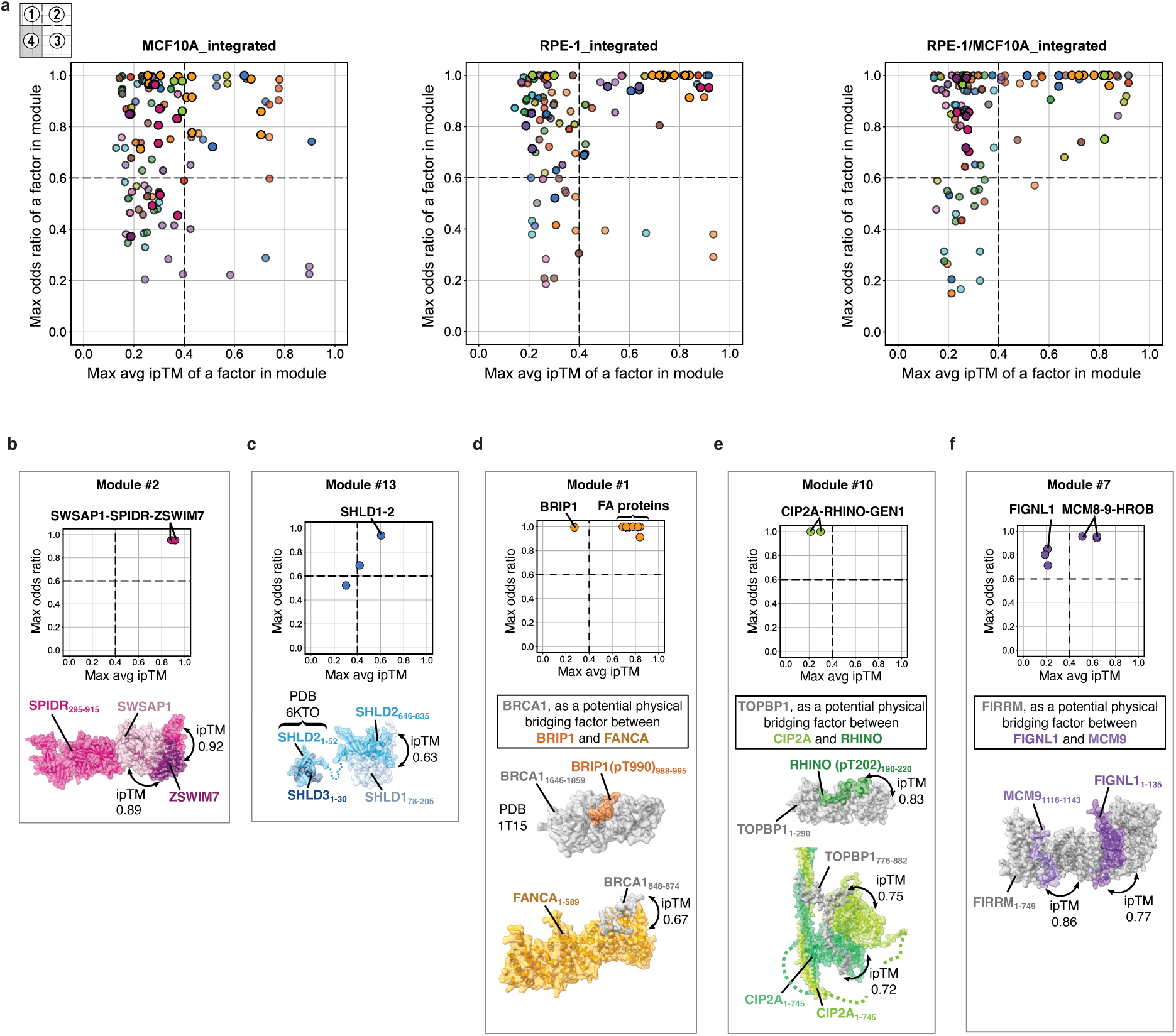
Prediction of potential physical contacts among members of genetic interaction modules. **a**, Scatter plots for all members of genetic modules in the indicated integrated datasets, showing the maximum average (max avg) ipTM score for each factor with another module member (x-axis) versus the maximum odds ratio of that factor with another module member (y-axis). Average ipTM scores were calculated across five AlphaFold2 models for each protein pair. Integrated datasets are indicated on top of the plots. Points in the right top quadrant (Q2) represent DDR factors which are robustly associated to their genetic module and have at least one member of the module that is predicted to interact directly with them, as determined by AlphaFold2 average ipTM score. Points in the left top quadrant (Q1) indicate robust members of gene modules for which a direct interactor was not predicted among the gene members of the same genetic module. The colors of the points indicate members belonging to the same module. Dashed lines in scatter plots indicate empirical cutoffs for strong module specificity (y > 0.6) and interaction confidence (average ipTM > 0.4). **b**-**f**, Selected modules from RPE-1_integrated dataset with each panel displaying a module-specific scatter plot (top) and a representative predicted protein complex structure (bottom). Protein surfaces in structural models are colored according to module identity and annotated with interacting domains and ipTM scores. **b**, Module #2 (SWASP1, SPIDR, ZSWIM7): High ipTM scores support stable complex formation among proteins within this module, consistent with the similarity of genetic interaction profiles (high odds ratios) and prior biochemical evidence^206^. **c**, Module #13 (SHLD1, SHLD2, SHLD3): An experimental structure (PDB 6KTO) demonstrates the direct interaction between SHLD2 and SHLD3, while SHLD1-SHLD2 direct interaction is modeled using AlphaFold2, consistent with previous literature^28,91^. **d**, Module #1 (BRIP1 and FA proteins): BRIP1 is the only member of module #1 for which no direct interaction to any of the module members could be identified. Search for potential bridging factors identified BRCA1 as a protein capable of direct interactions with BRIP1 phosphorylated at T990^207^ (PDB 1T15) and with FANCA as predicted by AlphaFold2, consistent with previously published biochemical evidence^208^. **e**, Module #10 (CIP2A, GEN1, RHINO): None of the members of module #10 exhibit strong predicted interactions. TOPBP1 is predicted to interact with both CIP2A, consistent with previous biochemical evidence^101,209^, and RHINO, as determined by AlphaFold3 modeling guided by the observation that this interaction is dependent on pThr202 phosphorylation^210^. **f**, Module #7 (FIGLN1, MCM8, MCM9, HROB): FIRRM, an established direct partner of FIGNL1^211,212^, is predicted to act as a likely bridge between FIGNL1 and MCM9 based on AlphaFold2 structural modeling, which could account for the robust similarities between FIGNL1 and the MCM8-9-HROB complex.

**Extended Data Fig. 11.**
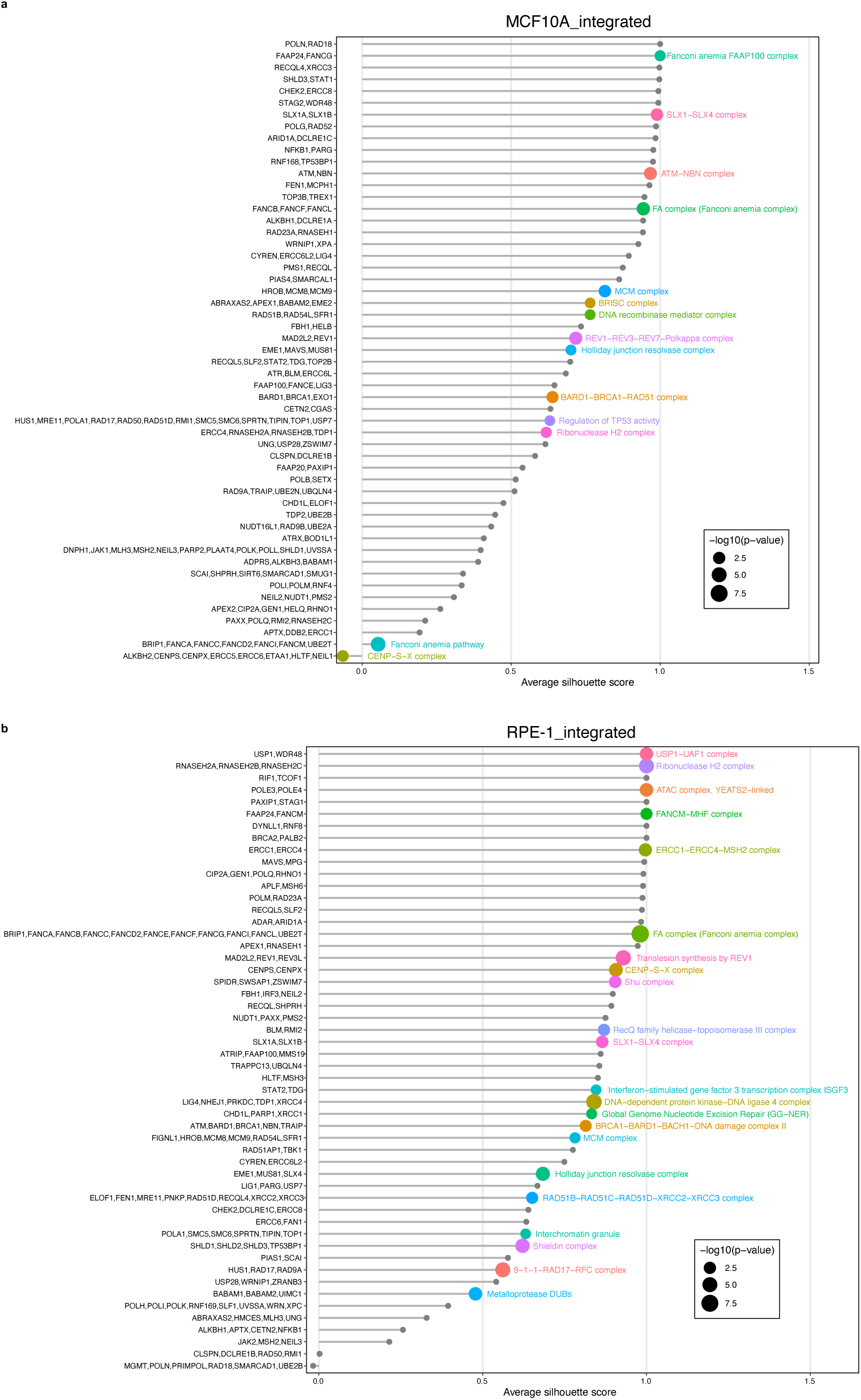
Functional enrichment analysis of GI modules. **a**, **b**, Lollipop plots showing gene ontology analysis results for GI modules in MCF10A_integrated and RPE-1_integrated datasets, respectively. Functional modules are listed on the y-axis, ordered by average silhouette score (x-axis) to quantify cluster cohesion. Point size represents the statistical significance of the most enriched term (-log10FDR) from four different sources: Gene Ontology Cellular Component (GO:CC), CORUM protein complexes, Reactome, and KEGG pathways.

**Extended Data Fig. 12.**
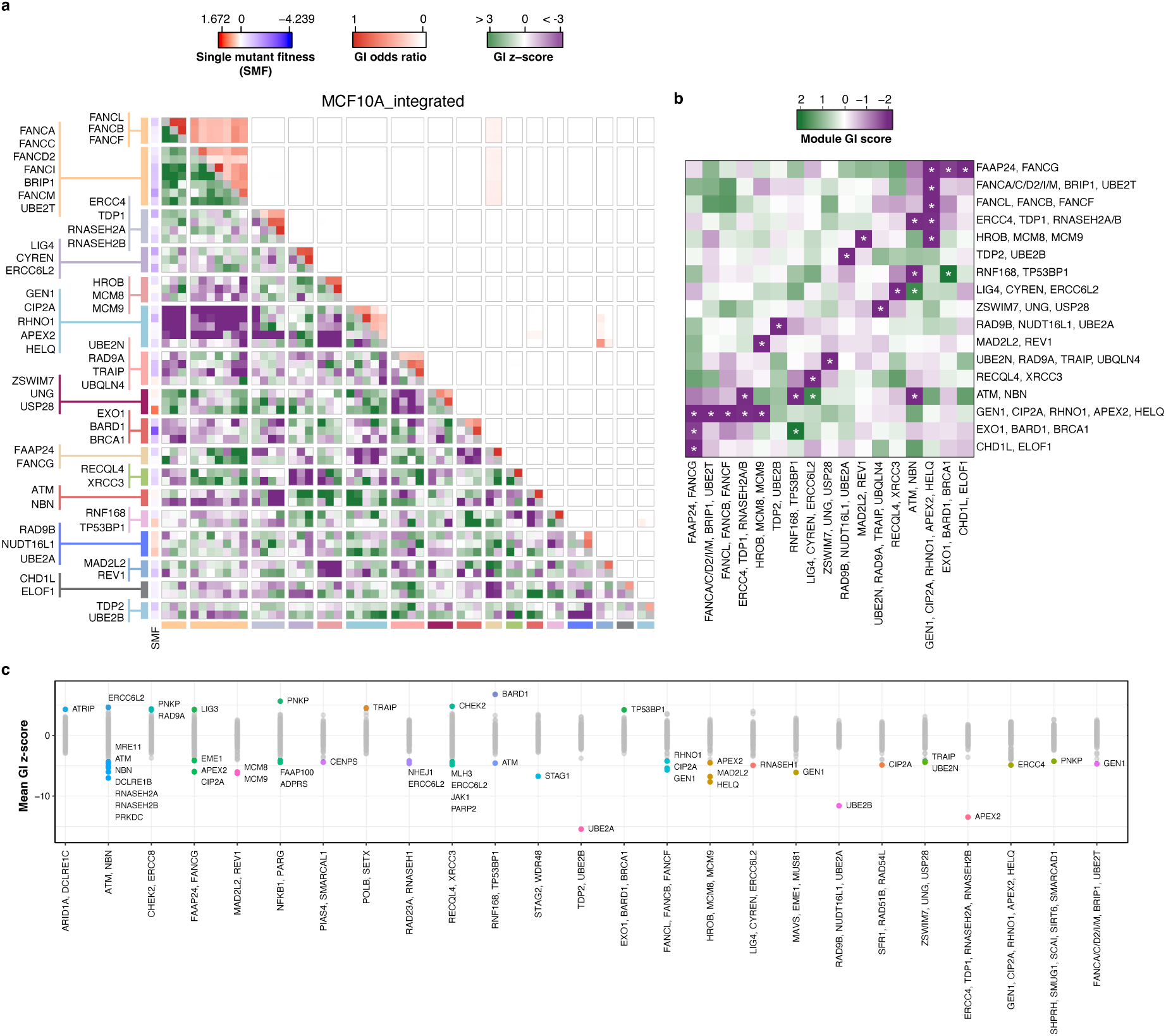
GI landscape of the MCF10A_integrated dataset. **a**, Filtered GI map highlighting genes from 17 high-confidence modules from the MCF10A_integrated dataset. The 233-gene library was filtered to include genes involved in top-scoring module-module interactions. As in the complete GI map, the lower heatmap triangle shows integrated GI z-scores and the upper heatmap triangle shows cluster odds ratios for each gene pair. Interaction of a gene with itself is depicted as grey in the diagonal. To the left of the heatmap, 17 GI modules are shown, together with each gene’s single mutant fitness (SMF). **b**, A reduced heatmap showing the significant module-module interactions from GI map in panel **a**. Significance is indicated (*), corresponding to a nominal two-sided p-value ≤ 0.1 under a normal approximation. **c**, Analysis of gene-module interactions. Each column represents a module (x-axis), and each point represent a single gene’s mean GI z-score with that module (y-axis). Colored datapoints indicate significant interactions (FDR ≤ 0.1). Modules without significant single gene interactors were omitted.

**Extended Data Fig. 13.**
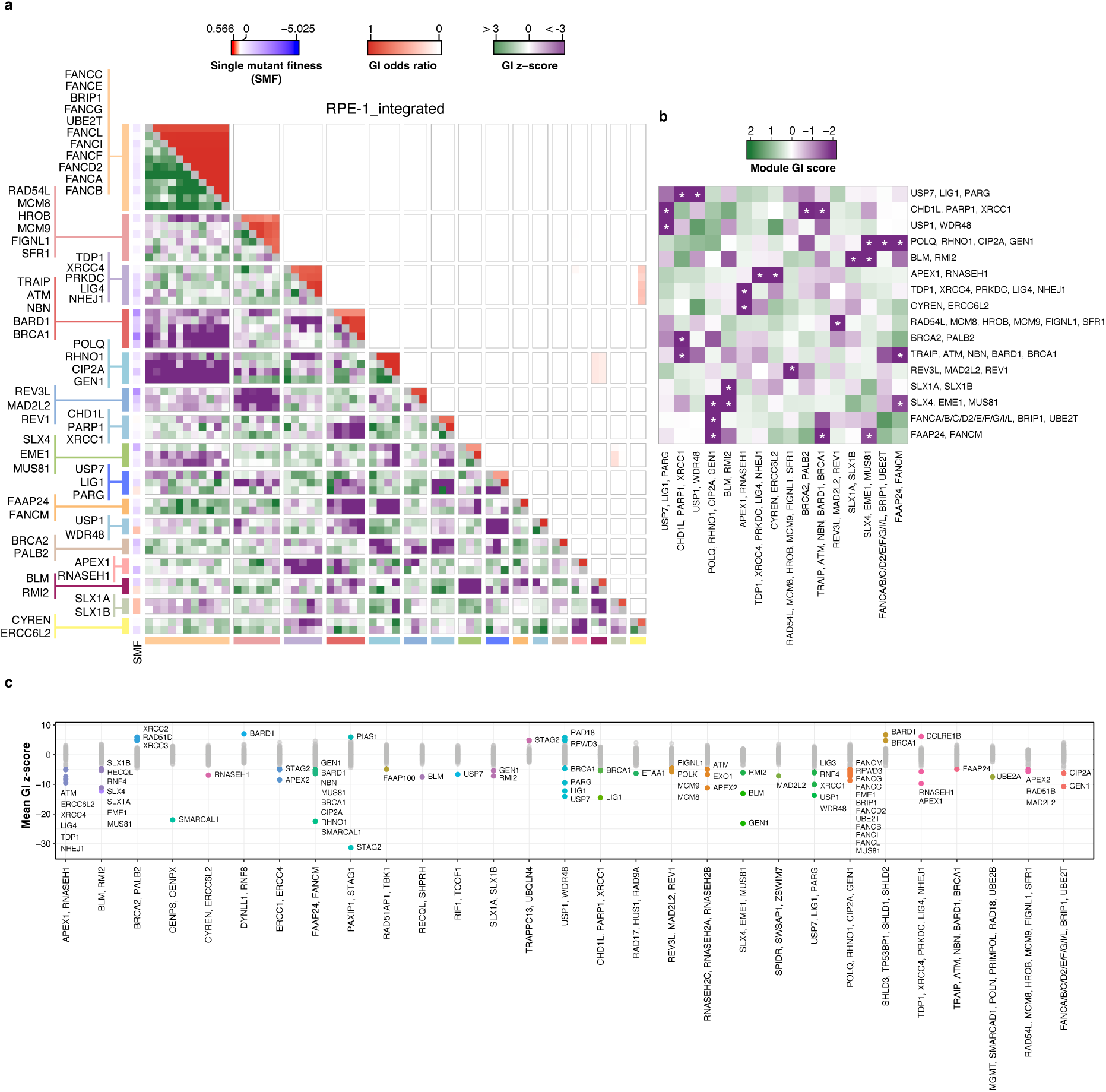
GI landscape of the RPE-1_integrated dataset. **a**, Filtered GI map highlighting genes from 16 high-confidence modules from the RPE-1_integrated dataset. The 233-gene library was filtered to include genes involved in top-scoring module-module interactions. As in the complete GI map, the lower heatmap triangle shows integrated GI z-scores and the upper heatmap triangle shows cluster odds ratios for each gene pair. Interaction of a gene with itself is depicted as grey in the diagonal. To the left of the heatmap, 16 GI modules are shown, together with each gene’s single mutant fitness (SMF). **b**, A reduced heatmap showing the significant module-module interactions from GI map in panel **a**. Significance is indicated (*), corresponding to a nominal two-sided p-value ≤ 0.1 under a normal approximation. **c**, Analysis of gene-module interactions. Each column represents a module (x-axis), and each point represent a single gene’s mean GI z-score with that module (y-axis). Colored datapoints indicate significant interactions (FDR ≤ 0.1). Modules without significant single gene interactors were omitted.

**Extended Data Fig. 14.**
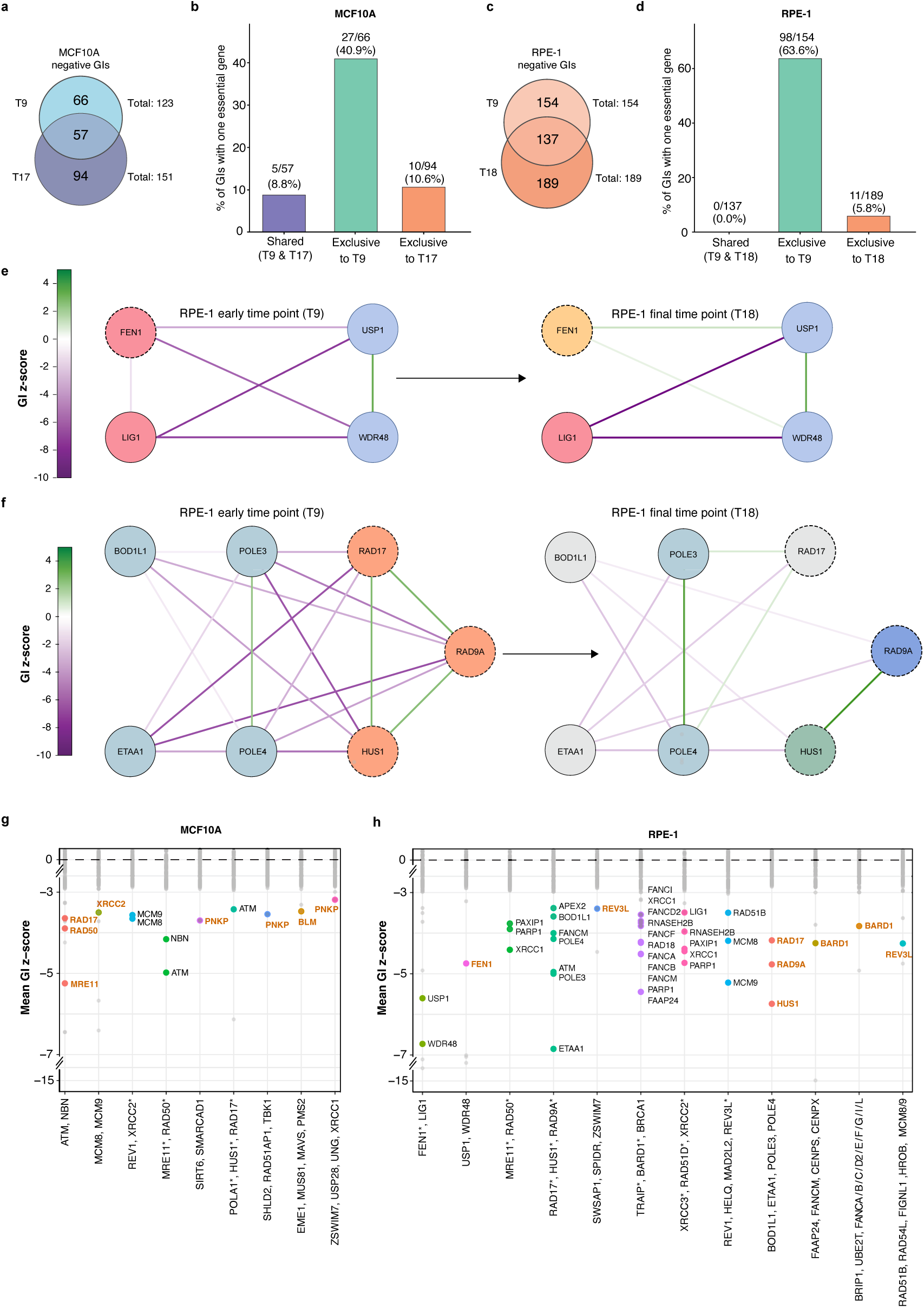
Temporal dynamics of genetic interactions involving essential genes. **a**, **c**, Venn diagrams showing unique and shared negative genetic interactions (GIs) between early (T9) and final (T17/T18) time points in the MCF10A and RPE-1 datasets, respectively. **b**, **d**, Percentage of significant negative GIs (FDR < 0.1) involving at least one essential gene exclusive to each time point or shared between them in the MCF10A and RPE-1 datasets, respectively. **e**, GI network for the *FEN1*, *LIG1* and *USP1, WDR48* modules at T9 in the RPE-1 dataset (left) and the corresponding network at T18 (right). **f**, GI network for the *BOD1L1, ETAA1, POLE3, POLE4* and *RAD17, HUS1, RAD9A* modules in the RPE-1 dataset at T9 (left) and T18 (right). **e**, **f**, Nodes are colored according to module assignments. Grey nodes denote unclustered genes. Nodes for essential genes are shown with a dashed outline. **g**, **h**, Analysis of negative gene-module interactions involving essential genes in the MCF10A and RPE-1 datasets at T9, respectively. Each column represents a module (x-axis), and each point represent a single gene’s mean GI z-score with that module (y-axis). Colored datapoints indicate significant interactions (FDR ≤ 0.1). Modules without significant single gene interactors were omitted. Essential genes are indicated by asterisks on the x-axis and highlighted in orange on the graph.

**Extended Data Fig. 15.**
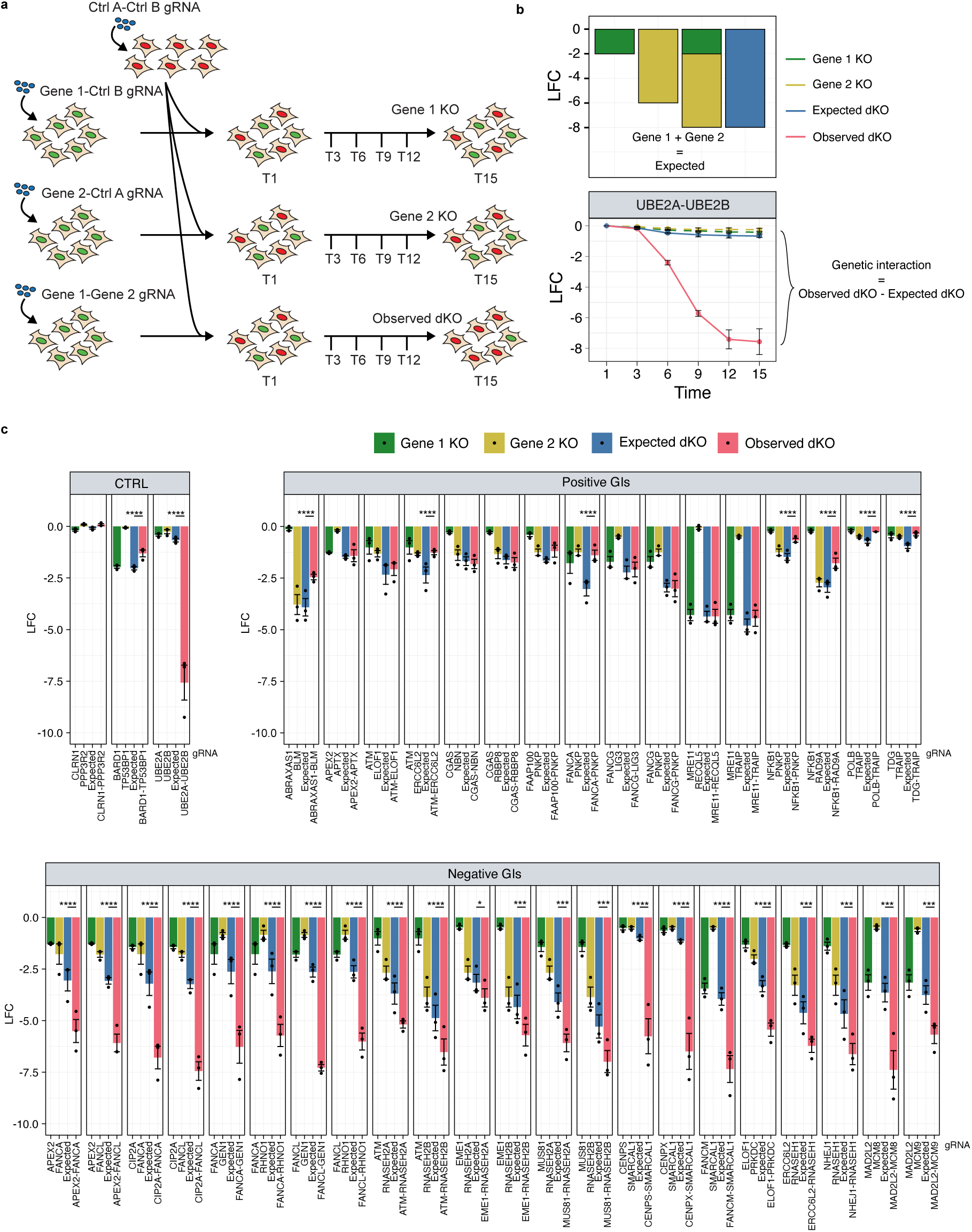
GI validation studies in MCF10A cells. **a**, Schematic of the two-color fluorescent competition assay used for validating hits of the GI screens. In brief, GFP or mCherry enCas12a cells are infected with lentiviral in4mer constructs (2 gRNAs per gene) targeting two control genes or combinations of genes of interest, respectively. Once virus selection is completed, GFP and mCherry cells are counted and mixed at an equal ratio. Mixed cell populations are maintained for 15 days (T15) and the fraction of GFP cells in the population is used to calculate relevant log fold changes (LFCs) vs T1 for each gene pair gRNA construct. **b**, Validation GI scoring method. Expected gene pair LFCs are calculated as the sum of the two relevant individual gene LFCs. GI validation scores are calculated as the difference between the observed LFC and the expected LFC for each gene pair. **c**, Validation LFCs at T15 for the MCF10A dataset. Cells were targeted with the indicated in4mer gRNAs. For each interaction, values are shown for individual single KOs, the dKO (observed), and the corresponding expected values for each dKO. Points represent the mean of three technical replicates and bars show the mean ± SEM (n = 3 biological replicates). Statistical significance was determined by two-way ANOVA comparing expected (blue) and observed (red) values at T15. **** = p-value ≤ 0.0001, *** = p-value ≤ 0.001, * = p-value ≤ 0.05.

**Extended Data Fig. 16.**
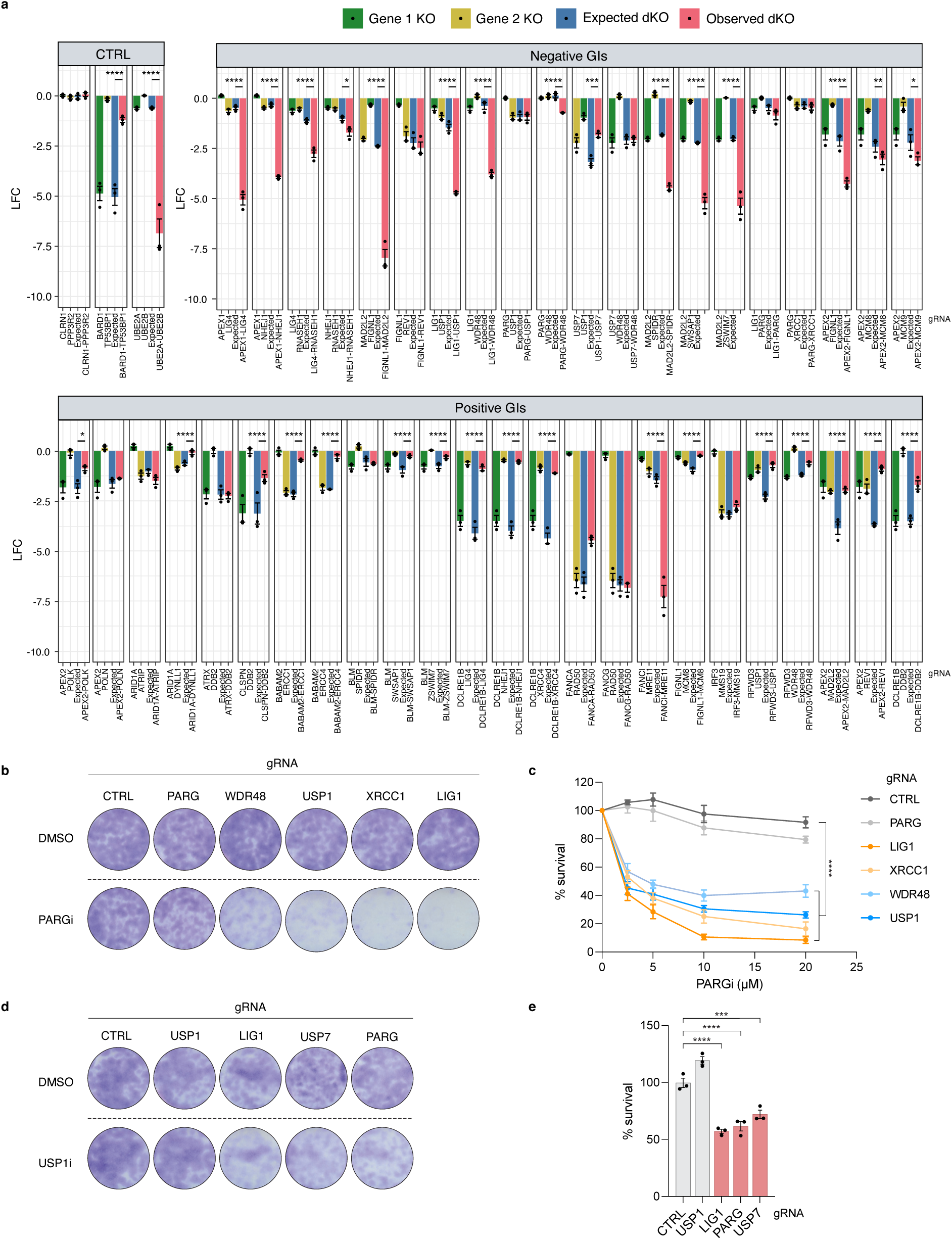
GI validation studies in RPE-1 *TP53^-/-^* cells. **a**, Validation LFCs at T15 for the RPE-1 dataset. Cells were targeted with the indicated in4mer gRNAs. For each interaction, values are shown for individual single KOs, the dKO (observed), and the corresponding expected values for each dKO. Points represent the mean of three technical replicates and bars show the mean ± SEM (n = 3 biological replicates). Statistical significance was determined by two-way ANOVA comparing expected (blue) and observed (red) values at T15. **b**, Representative survival assay of RPE-1 *TP53^-/-^* cells targeted with the indicated in4mer gRNAs and continuously treated for 7-8 days with DMSO vehicle or 20 µM PARG inhibitor (PARGi; PDD0017273). **c**, Quantification of survival shown in panel **b** across increasing concentrations of PARGi, as indicated. *PARG* KO cells and *WDR48* KO cells were used as a negative and positive controls, respectively. Survival is normalized to DMSO-treated controls. Bars represent the mean ± SEM (n = 3 biological replicates). Statistical significance was determined by two-way ANOVA at the highest dose of PARGi (20 µM)**. d**, **e**, Representative survival assay and quantification, respectively, of RPE-1 *TP53^-/-^* cells targeted with the indicated in4mer gRNAs and continuously treated for 7-8 days with DMSO vehicle or 3.125 µM USP1 inhibitor (USP1i; KSQ-4279). *USP1* KO cells and *LIG1* KO cells were used as a negative and positive controls, respectively. Percent survival is calculated as survival after USP1i treatment divided by survival after DMSO treatment for each sample. Bars represent the mean ± SEM (n = 3 biological replicates). Statistical significance was determined by one-way ANOVA. **** = p-value ≤ 0.0001, *** = p-value ≤ 0.001, ** = p-value ≤ 0.01, * = p-value ≤ 0.05.

**Extended Data Fig. 17.**
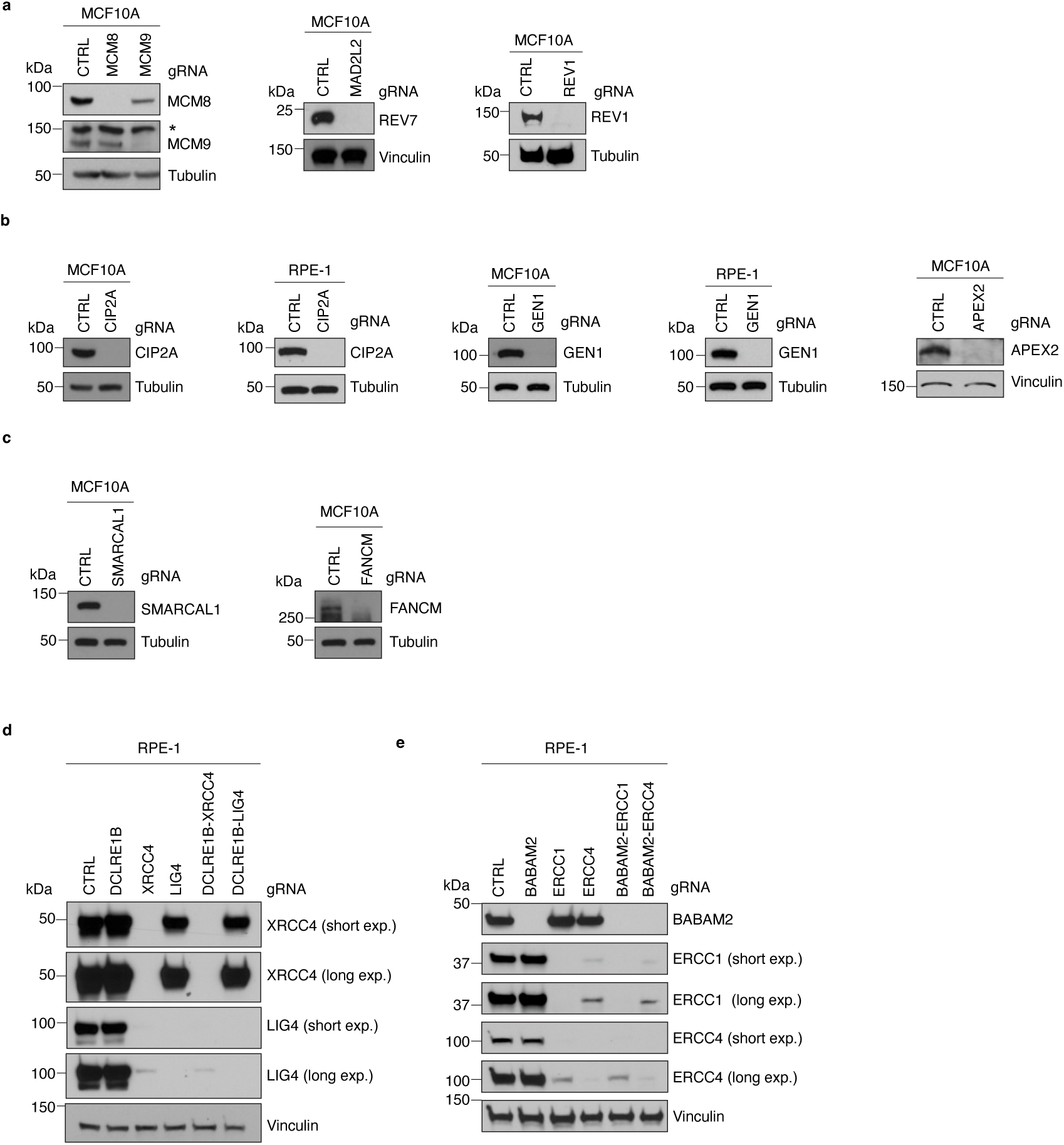
Analysis of Cas12a gene editing by western blotting for negative and positive genetic interactions. **a-f**, Cells were targeted with the indicated in4mer gRNAs, and western blot samples were collected 6-9 days after selection was completed. **a**, Western blots showing MCM8, MCM9, REV7, and REV1 levels in MCF10A cells. Asterisk indicates a non-specific band. **b**, Western blots showing CIP2A, GEN1, and APEX2 levels in MCF10A and RPE-1 *TP53^-/-^* cells, as indicated. APEX2 migrates between 50 and 75 kDa (markers not shown due to distance from band). **c**, Western blots showing SMARCAL1 and FANCM levels in MCF10A cells. **d**, Western blots showing XRCC4 and LIG4 levels in RPE-1 *TP53^-/-^* cells. **e**, Western blots showing BABAM2, ERCC1, and ERCC4 levels in RPE-1 *TP53^-/-^* cells. **a**-**e**, Tubulin or vinculin were used as loading controls. Exp. stands for exposure.

**Extended Data Fig. 18.**
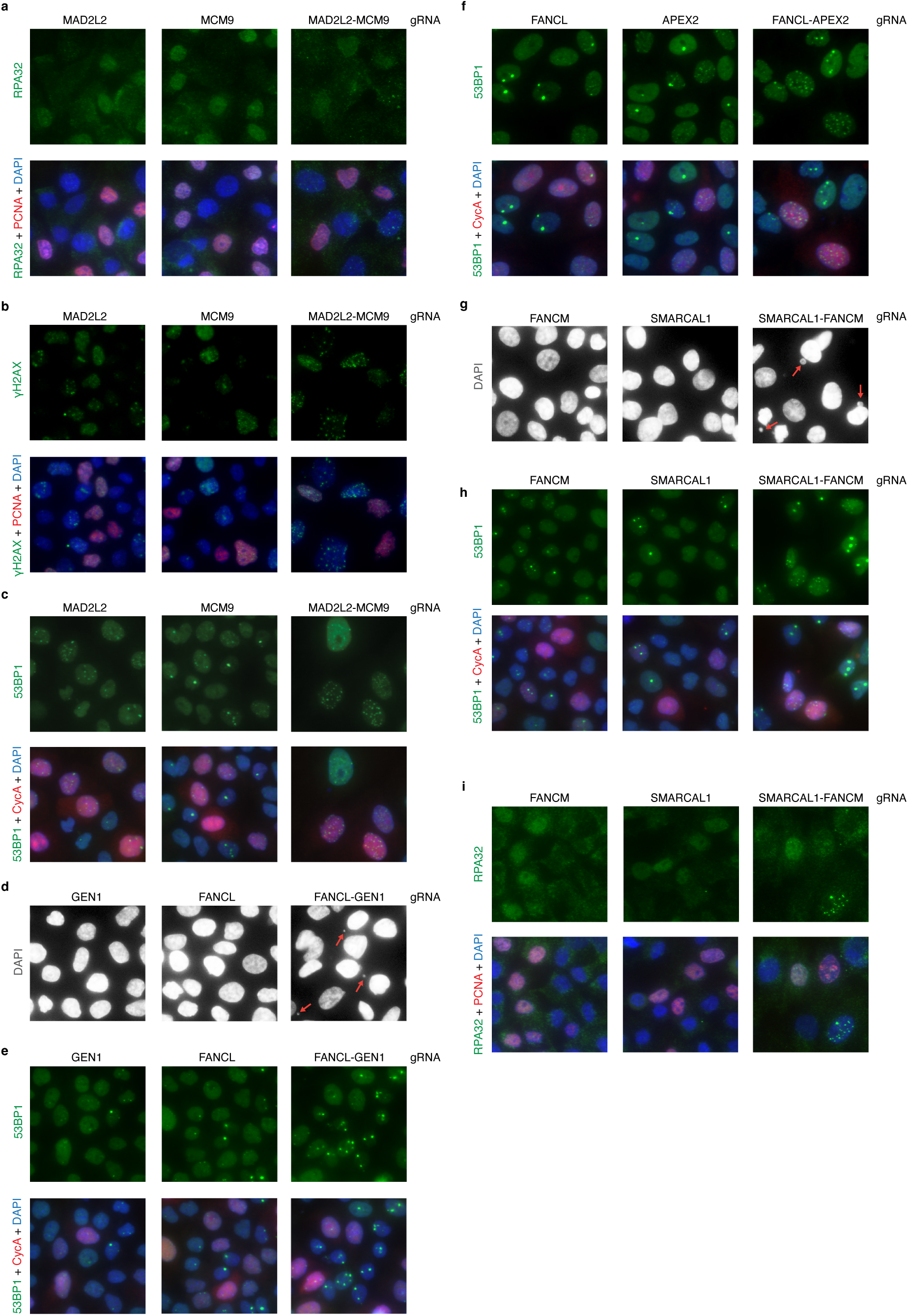
Representative immunofluorescence images of DNA damage markers. **a**-**i**, Cells were targeted with the indicated in4mer gRNAs, and immunofluorescence experiments were carried out 7 days after selection was completed. **a**-**c**, Representative immunofluorescence images of RPA32, γH2AX, and 53BP1 staining (green), respectively, with and without DAPI (blue) and PCNA/cyclin A (CycA) (red) staining for the *MAD2L2*-*MCM9* interaction in MCF10A cells quantified in Fig. 4d-f. **d**, Representative immunofluorescence images of DAPI staining for the *FANCL-GEN1* interaction in MCF10A cells. Red arrows indicate micronuclei quantified in Fig. 5c (middle). **e**, **f**, Representative immunofluorescence images of 53BP1 staining (green) with and without DAPI (blue) and CycA (red) staining for the *FANCL-GEN1* and *FANCL-APEX2* interactions in MCF10A cells, respectively, quantified in Fig. 5c (bottom) and Extended Data Fig. 21c,d. **g**, Representative immunofluorescence images of DAPI staining for the *SMARCAL1*-*FANCM* interaction in MCF10A cells. Red arrows indicate micronuclei quantified in Fig. 6d. **h**, **i**, Representative immunofluorescence images of 53BP1 and RPA32 staining (green), respectively, with and without DAPI (blue) and CycA/PCNA (red) staining for the *SMARCAL1*-*FANCM* interaction in MCF10A cells quantified in Fig. 6e-g.

**Extended Data Fig. 19.**
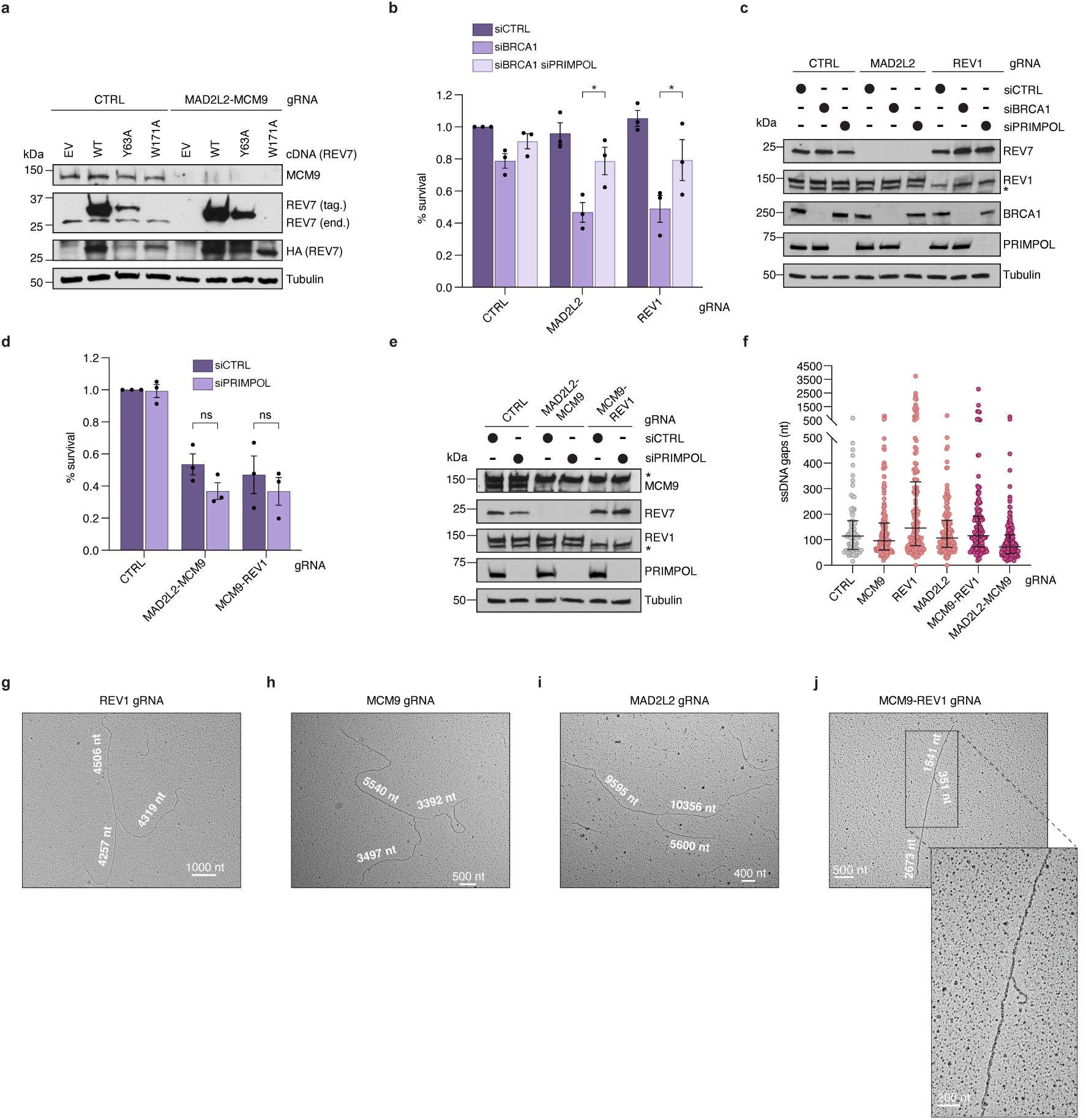
Characterization of the negative interactions between the *REV1/MAD2L2/REV3L* module and *BRCA1* or the *MCM8/MCM9/HROB* module. **a**-**k**, Cells were targeted with the indicated in4mer gRNAs. **a**, Western blot showing the levels of MCM9, endogenous (end.) REV7, and FLAG-HA-tagged (tag.) gRNA-resistant REV7 variants in MCF10A cells examined in the survival assays shown in Fig. 4g,h. The level of endogenous REV7 is reduced in cells targeted with MAD2L2-MCM9 in4mer gRNA, while gRNA-resistant REV7 cDNA constructs remain expressed. REV7 W171A mutant cannot be detected with the anti-REV7 antibody because the mutation disrupts the antibody epitope but is detected with an anti-HA antibody. Tubulin was used as a loading control. **b**, **d**, Quantification of survival assays in MCF10A cells treated with either control (siCTRL), PRIMPOL (siPRIMPOL), BRCA1 (siBRCA1) siRNA, or a combination of siBRCA1 and siPRIMPOL. Bars represent the mean percentage of survival (± SEM) relative to the CTRL sample treated with siCTRL (n = 3 biological replicates). Statistical significance was determined by one-way ANOVA. **c**, Western blot showing the levels of REV7, REV1, BRCA1, and PRIMPOL in MCF10A cells examined in survival assays in panel **b**. **e**, Western blot showing the levels of MCM9, REV7, REV1, and PRIMPOL in MCF10A cells examined in survival assays in panel **d**. **c**, **e**, Tubulin was used as a loading control. Asterisks indicate non-specific bands. **f**, Quantification of ssDNA gap lengths at and behind replication forks (in nucleotides, nt), detected by EM in MCF10A cells. Scatter plot shows individual values (n = 2 biological replicates). At least 41 replication intermediates were examined per sample per experiment. Bars represent the median and interquartile range. **g**-**j**, Representative images of replication forks detected by EM in MCF10A cells. The length of the replication fork branches is indicated in nucleotides (nt). Scale bars are shown in nt. Quantification of EM experiments is shown in Fig. 4k and panel **f.** * = p-value ≤ 0.05.

**Extended Data Fig. 20.**
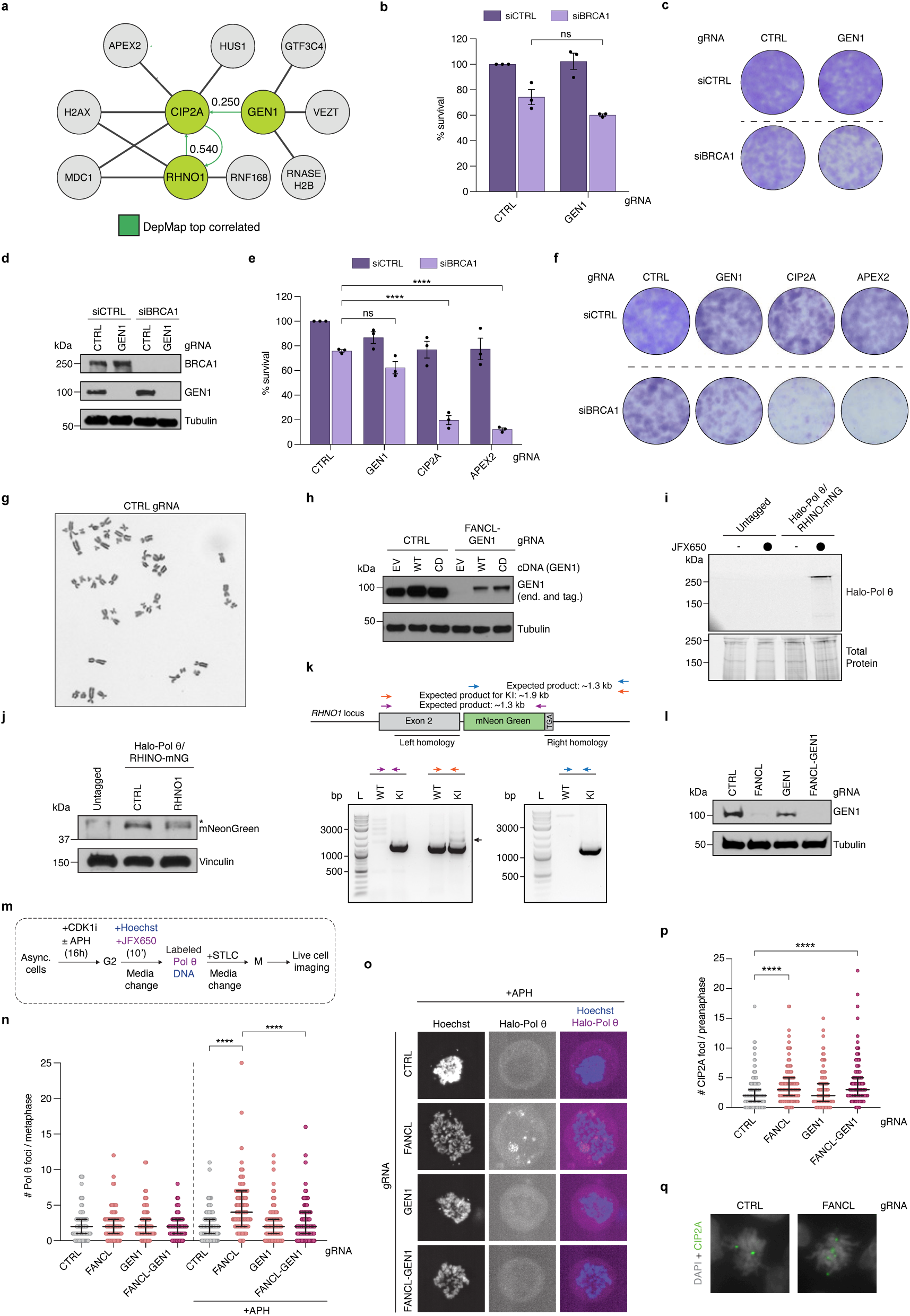
Characterization of the negative interactions between Fanconi anemia genes and the mitotic DNA repair genes *CIP2A*, *GEN1*, and *RHNO1*. **a**, Network representation of top DepMap (24Q4) correlated hits for *CIP2A*, *GEN1,* and *RHNO1*. The top 4 correlated genes for *CIP2A*, *GEN1,* and *RHNO1* are shown, with green edges indicating the top correlated gene for the indicated parent node. Displayed numbers are Pearson correlation scores from DepMap. **b**-**h**, **j**, **l**, **n**-**q**, Cells were targeted with the indicated in4mer gRNAs. **b**, **c**, Quantification of survival assays and representative survival, respectively, of control (CTRL) and *GEN1* KO RPE-1 *TP53^-/-^* cells treated with control (siCTRL) or BRCA1 (siBRCA1) siRNA. Bars represent the mean percentage of survival (± SEM) relative to the CTRL sample treated with siCTRL (n = 3 biological replicates). Statistical significance was determined by unpaired t test. **d**, Western blot showing BRCA1 and GEN1 levels in RPE-1 *TP53^-/-^* cells examined in survival assays in panels **b** and **c**. Tubulin was used as a loading control. **e, f**, Quantification of survival assays and representative survival, respectively, of CTRL, *GEN1, CIP2A,* and *APEX2* KO RPE-1 *TP53^-/-^* cells treated with siCTRL or siBRCA1. Bars represent the mean percentage of survival (± SEM) relative to the CTRL sample treated with siCTRL (n = 3 technical replicates). Statistical significance was determined by one-way ANOVA. **g**, Representative image of a metaphase from CTRL RPE-1 *TP53^-/-^* cells quantified in Fig. 5d. **h**, Western blot showing the levels of endogenous (end.) and FLAG-HA-tagged (tag.) GEN1 variants in MCF10A cells examined in survival assays in Fig. 5h,i. The level of endogenous GEN1 is reduced in cells targeted with FANCL-GEN1 in4mer gRNA, while the expression of gRNA-resistant GEN1 constructs is not affected. Tubulin was used as a loading control. **i**, Western blot confirming expression of Halo-Pol θ in U2OS Halo-Pol θ RHINO-mNeonGreen cells, compared to the parental U2OS cell line (untagged). Total protein levels are shown. **j**, Western blot confirming expression of RHINO-mNeonGreen in U2OS Halo-Pol θ RHINO-mNeonGreen cells, compared to the parental U2OS cell line (untagged), using an anti-mNeonGreen antibody. *RHNO1* was targeted with in4mer gRNA to confirm the specificity of the signal. Asterisk marks a non-specific band. Vinculin was used as a loading control. **k**, Schematic of the *RHNO1* locus showing primer pairs used for PCR (top). Expected size of PCR products is indicated. Genotyping PCR confirming knock-in (KI) of mNeonGreen-tag in the *RHNO1* locus of U2OS Halo-Pol θ cells (bottom). Black arrow marks the KI PCR product. **l**, Western blot showing GEN1 levels in U2OS Halo-Pol θ RHINO-mNeonGreen cells employed in live-cell imaging experiments in Fig. 5j,k and panels **n** and **o**. Tubulin was used as a loading control. **m** Schematic of the protocol used for live-imaging of U2OS Halo-Pol θ RHINO-mNeonGreen cells quantified in Fig. 5j,k (RHINO-mNeonGreen foci) and panels **n** and **o** (Halo-Pol θ foci). Asynchronous (async.) cells were synchronized for 16 h in G2/M with 9 µM CDK1 inhibitor (CDK1i; RO-3306) in the presence or absence of 0.4 µM aphidicolin (APH), followed by 10 min labelling of DNA with Hoechst and Halo-Pol θ with JFX65, and subsequent arrest of cells in mitosis with 10 µM S-trityl-L-cysteine (STLC). **n**, Quantification of mitotic Halo-Pol θ foci in live-cell imaging experiments in U2OS Halo-Pol θ RHINO-mNeonGreen cells. Scatter plot shows individual values (n = 3 biological replicates). At least 24 mitotic cells were examined per sample per experiment. Bars represent the median and interquartile range. Statistical significance was determined by Mann-Whitney test. **o**, Representative images of mitotic Halo-Pol θ foci in APH-treated cells quantified in panel **n**. **p**, **q**, Quantification of spontaneous CIP2A foci and representative images, respectively, in pre-anaphase mitotic MCF10A cells upon disruption of *FANCL* and/or *GEN1*. Scatter plot shows individual values (n = 3 biological replicates). Bars represent the median and interquartile range. At least 39 pre-anaphase cells were scored per sample per experiment. Statistical significance was determined by Mann-Whitney test. **** = p-value ≤ 0.0001.

**Extended Data Fig. 21.**
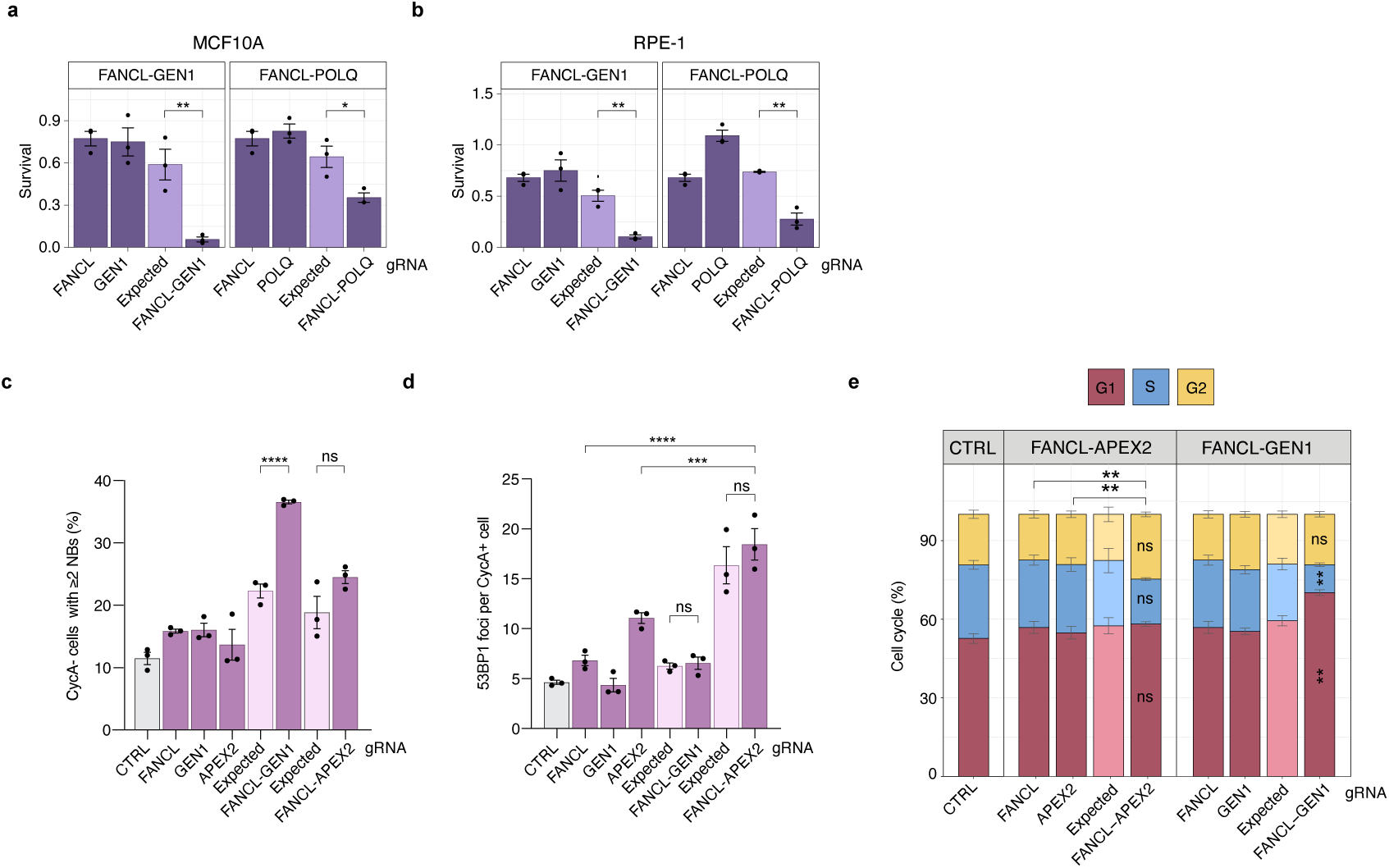
Characterization of the negative interactions between Fanconi anemia genes and *APEX2* or the *CIP2A*/*GEN1*/*RHNO1* module. **a-e**, Cells were targeted with the indicated in4mer gRNAs. For each interaction, values are shown for individual single KOs, the dKO (observed), and the corresponding expected values for each dKO, calculated from single KO phenotypes. Expected values are plotted immediately to the left of their matched observed dKO values for each gene pair. Significant deviations between observed and expected values are indicated. Bars represent the mean ± SEM (n = 3 biological replicates). **a**, **b** Quantification of survival assays for the indicated genetic interactions in MCF10A and RPE-1 *TP53^-/-^* cells. Survival values are plotted relative to survival values of control samples, which are equal to 1 and are not shown on the graphs. Statistical significance was assessed by unpaired t test. **c**, **d**, Quantification of 53BP1 nuclear bodies (NBs) in cyclin A-negative (CycA-, G1) MCF10A cells (**c**) and 53BP1 foci in cyclin A-positive (CycA+, S/G2) MCF10A cells (**d**). **e**, Cell cycle profiles for MCF10A cells. G1, S, and G2 cell cycle phases were determined based on PCNA and DAPI staining. In addition to the statistical comparison between observed and expected values, the significance of the difference in G2-phase cells for the indicated samples is shown. **c**-**e**, Bars represent the mean ± SEM (n = 3 biological replicates). Statistical significance was determined by one-way ANOVA. **** = p-value ≤ 0.0001, *** = p-value ≤ 0.001, ** = p-value ≤ 0.01, * = p-value ≤ 0.05.

**Extended Data Fig. 22.**
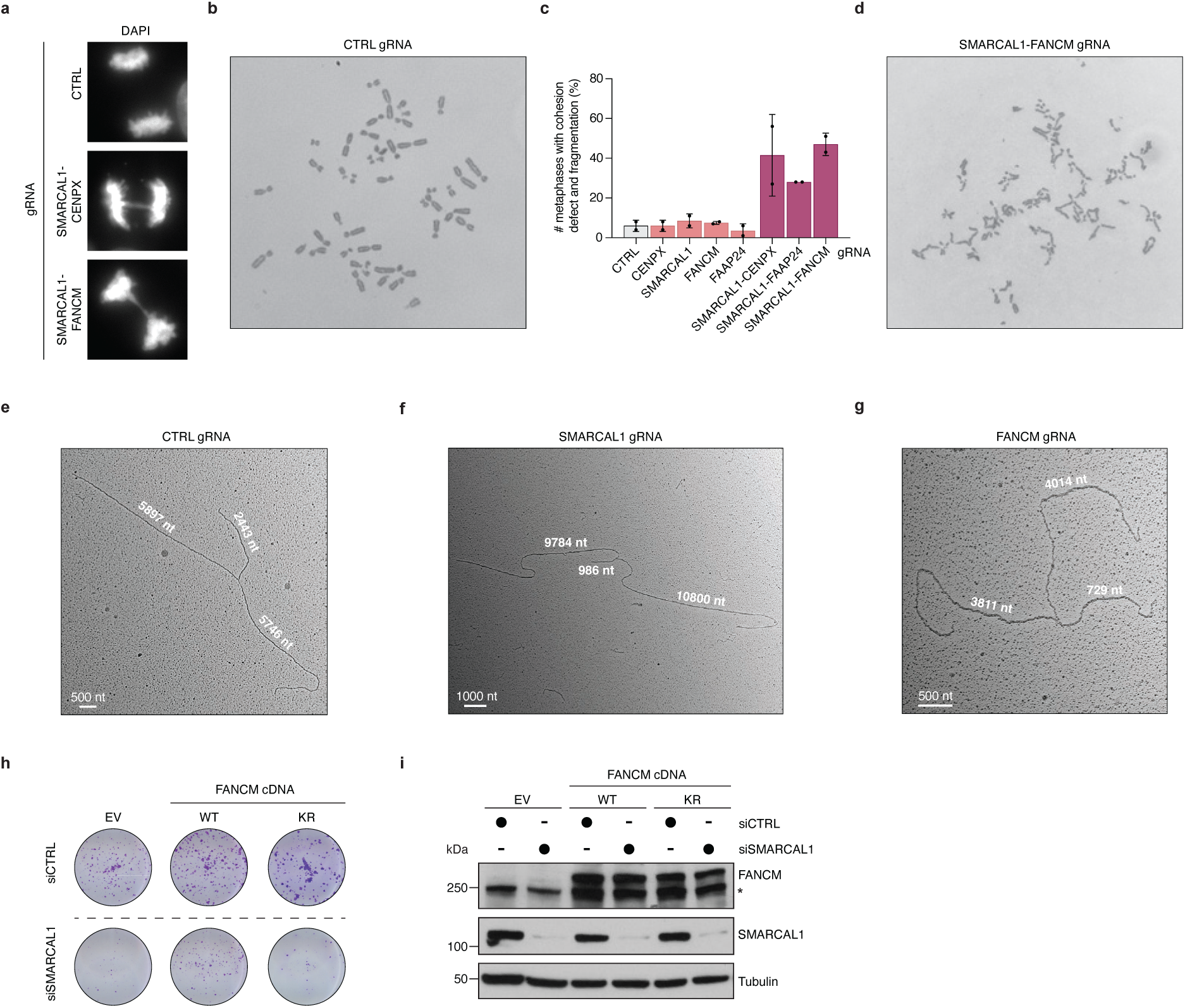
Supporting data for the negative interactions between *SMARCAL1* and genes encoding FANCM and its partners. **a-g**, Cells were targeted with the indicated in4mer gRNAs. **a**, Representative images of anaphases observed in control (CTRL), *SMARCAL1-CENPX*, and *SMARCAL1-FANCM* dKO MCF10A cells quantified in Fig. 6h,i. **b**, Representative image of a metaphase from CTRL RPE-1 *TP53^-/-^* cells quantified in Fig. 6j and panel **c. c**, Quantification of metaphases with cohesion defect and chromosome fragmentation in RPE-1 *TP53^-/-^* cells. Metaphases with this defect were not included in the assessment of metaphase DNA breaks quantified in Fig. 6j. Bars represent the mean percentage of metaphases with the defect (± SEM) relative to the total number of metaphases examined for each sample (n = 2 biological replicates). At least 100 metaphases were analyzed per sample per experiment. **d**, Representative image of a metaphase with cohesion defect and chromosome fragmentation from the *SMARCAL1-FANCM* dKO RPE-1 *TP53^-/-^* cells quantified in panel **c**. **e**-**g**, Representative images of replication forks detected by EM in MCF10A cells and quantified in Fig. 6l. The length of the replication fork branches is indicated in nucleotides (nt). Scale bars are shown in nt. **h**, Representative survival assay of FaDu cells quantified in Fig. 6n. **i**, Western blot showing FANCM and SMARCAL1 levels in FaDu cells examined in survival assays in Fig. 6n. Asterisk marks a non-specific band. Tubulin was used as a loading control.

**Extended Data Fig. 23.**
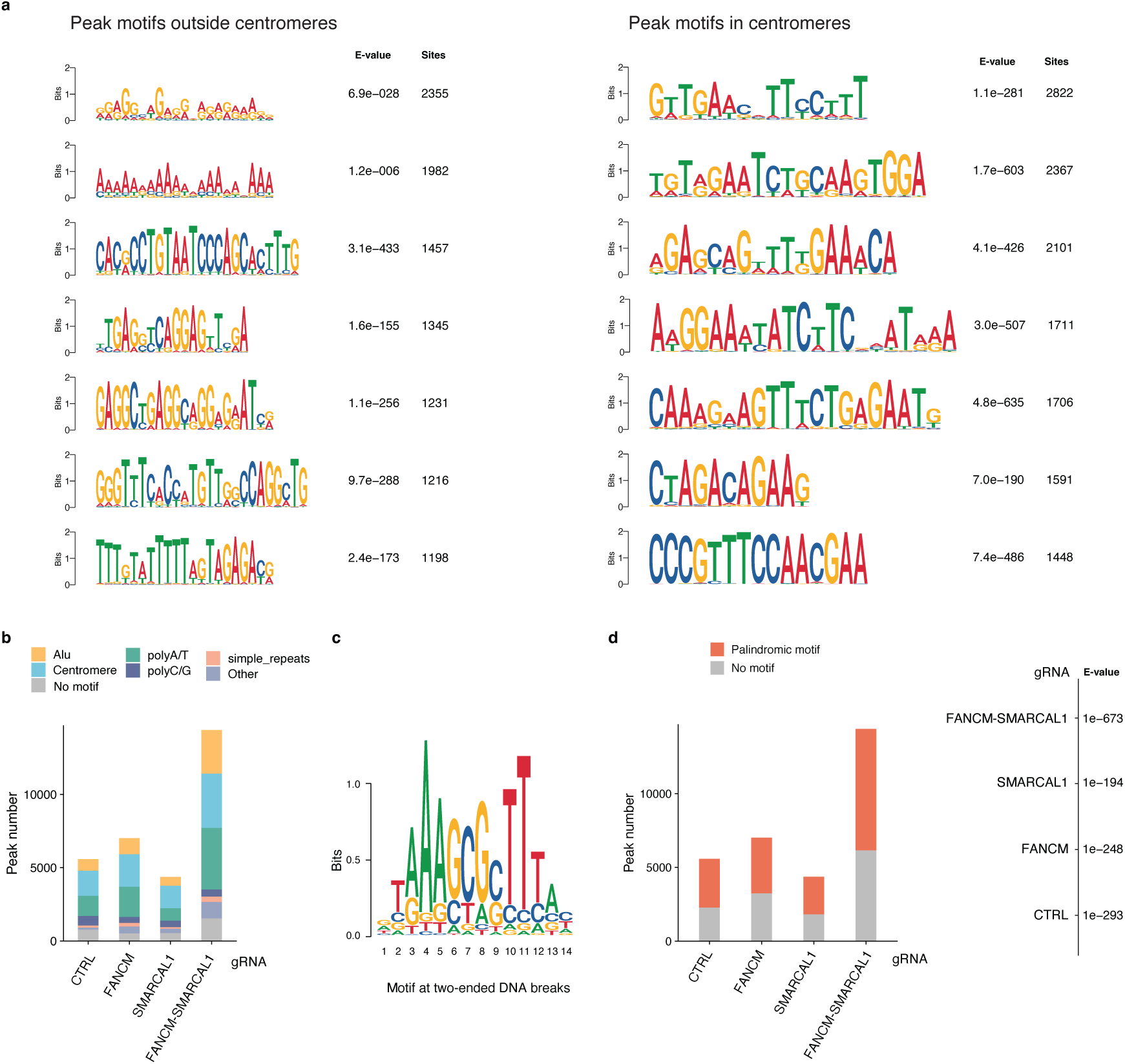
Mapping of DNA double-strand breaks underlying the *SMARCAL1-FANCM* negative interaction by END-seq. **a**, Representative sequence motifs identified by MEME analysis in peaks located at centromeres or other genomic loci in MCF10A cells treated with gRNAs targeting *FANCM* and *SMARCAL1*. **b**, Histogram showing the number of peaks in MCF10A cells treated with the indicated in4mer gRNAs. Peaks identified at Alu elements, centromeres, polyA/T, polyC/G, and simple repeats are shown in different colors. **c**, Palindromic motif enriched at the signal summit of two-ended DNA breaks in MCF10A cells deficient for FANCM and SMARCAL1. **d**, Histogram displaying the number of DNA breaks near the peaks containing the palindromic motif shown in **c** in MCF10A cells treated with the indicated gRNAs.

**Extended Data Fig. 24.**
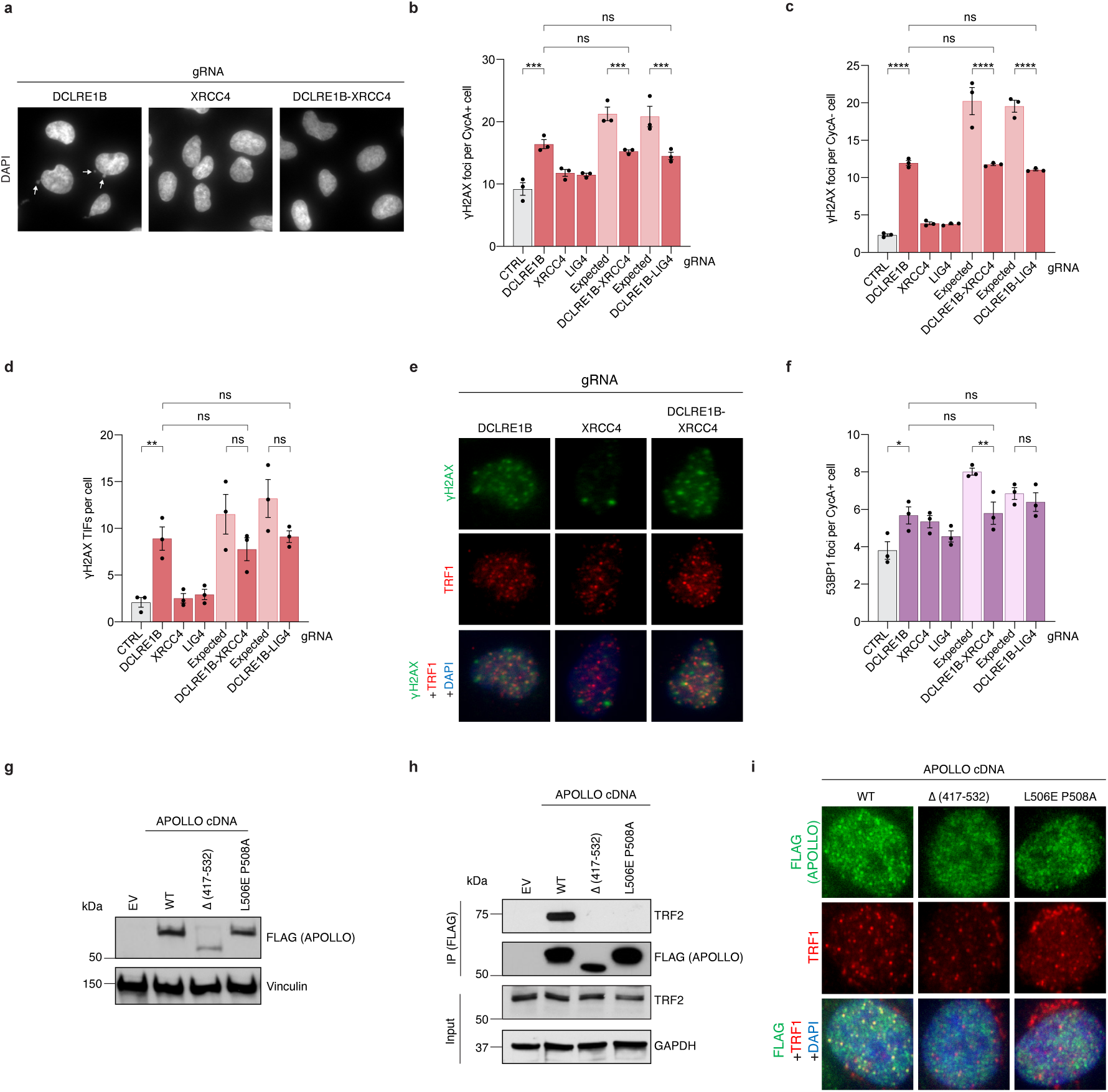
Supporting data for the suppressor interaction between *DCLRE1B* and the *XRCC4/LIG4/NHEJ1* module. **a-f**, Cells were targeted with the indicated in4mer gRNAs. **a**, Representative immunofluorescence images of DAPI staining for the *DCLRE1B-XRCC4* interaction in RPE-1 *TP53^-/-^* cells. White arrows indicate micronuclei quantified in Fig. 7c. **b**-**d, f**, For each interaction, values are shown for individual single KOs, the dKO (observed), and the corresponding expected values for each dKO, calculated from single KO phenotypes. Expected values are plotted immediately to the left of their matched observed dKO values for each gene pair. Bars represent the mean ± SEM (n = 3 biological replicates). Statistical significance was determined by one-way ANOVA. **b**, **c**, Quantification of γH2AX foci in the indicated cyclin A-positive (CycA+, S/G2, **b**) and cyclin A-negative (CycA-, G1, **c**) RPE-1 *TP53^-/-^* cells. Total number of γH2AX foci per cell is quantified in Fig. 7f. **d**, Quantification of γH2AX foci that co-localize with TRF1 foci (TIFs) in RPE-1 *TP53^-/-^* cells. **e**, Representative immunofluorescence images of γH2AX (green), TRF1 (red), and γH2AX together with TRF1 and DAPI (blue) staining quantified in panel **d**. **f**, Quantification of 53BP1 foci in cyclin A-positive (CycA+, S/G2) RPE-1 *TP53^-/-^*cells. **g**, Western blot showing levels of FLAG-tagged APOLLO in RPE-1 *TP53^-/-^* cells targeted with DCLRE1B in4mer gRNA examined in survival assays in Fig. 7j. Vinculin was used as a loading control **h**, Co-immunoprecipitation assay showing the effects of TRF2-binding-deficient APOLLO mutants on the APOLLO-TRF2 interaction in RPE-1 *TP53^-/-^* cells. **i**, Representative immunofluorescence images of FLAG-tagged APOLLO (green), TRF1 (red), and FLAG/TRF1/DAPI (blue) staining of RPE-1 *TP53^-/-^* cells with higher expression levels of APOLLO compared to images in Fig. 7h. **** = p-value ≤ 0.0001, *** = p-value ≤ 0.001, ** = p-value ≤ 0.01, * = p-value ≤ 0.05.

**Extended Data Fig. 25.**
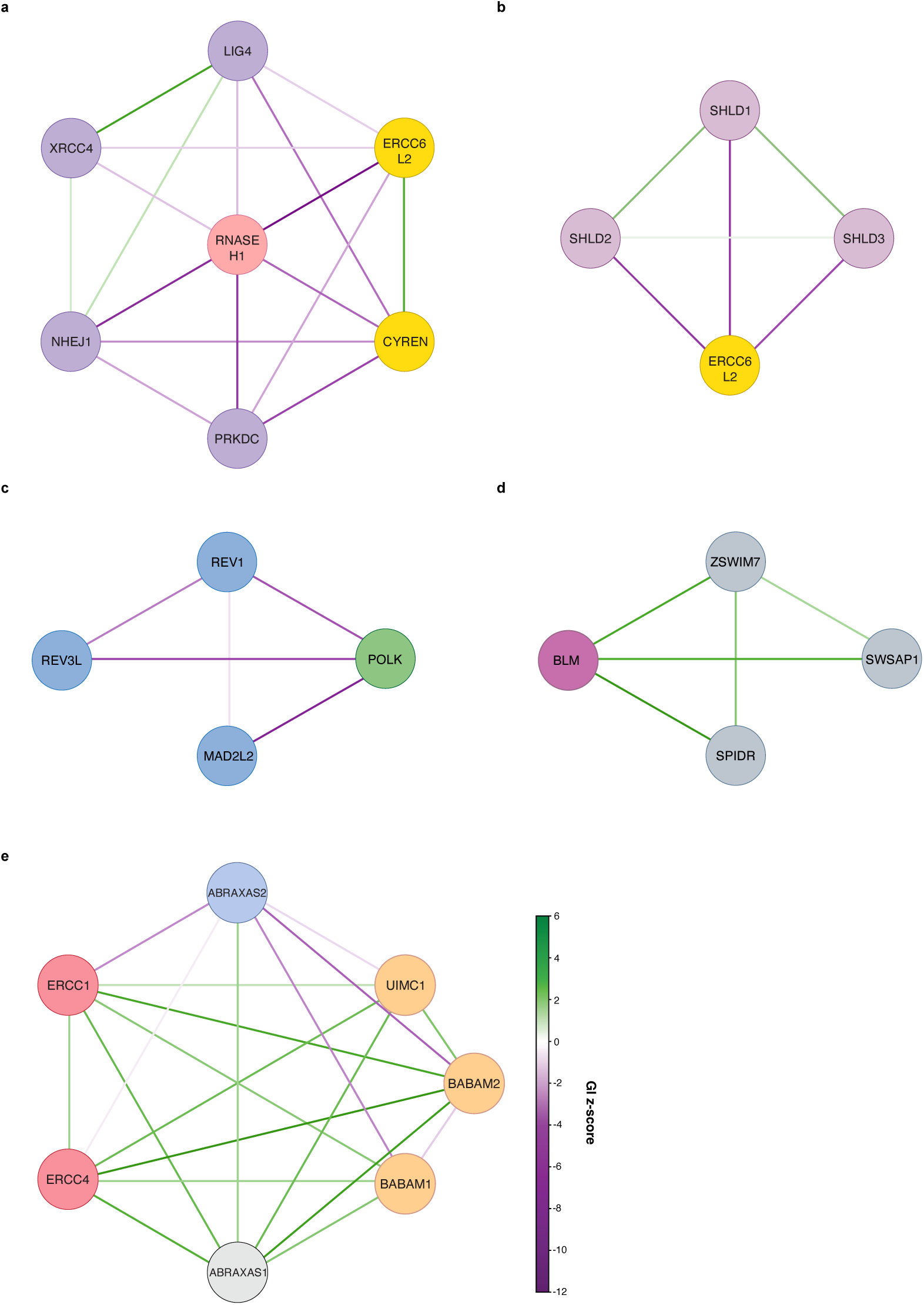
Notable GI networks from the RPE-1_integrated dataset. **a**, GI network for the *RNASEH1* and NHEJ modules (*XRCC4/LIG4/NHEJ1/PRKDC* and *ERCC6L2*/*CYREN*). **b**, GI network for the *ERCC6L2* and the shieldin module (*SHLD1/SHLD2/SHLD3*). **c**, GI network for the *POLK* and the TLS module (*REV1*/*MAD2L2*/*REV3L)*. **d**, GI network for the *BLM* and the Shu complex module (*ZSWIM7/SPIDR/SWSAP1*). **e**, GI network for the *ERCC1/ERCC4* module, the *BABAM1/BABAM2/UIMC1* module, *ABRAXAS1*, and *ABRAXAS2*. **a**-**e**, GI z-scores < 6 and > −12 are shown. Node colors are according to the module coloring scheme in Extended Data Fig. 13a. Grey nodes denote unclustered genes.

